# RNA polymerase II pausing regulates a quiescence-dependent transcriptional program, priming cells for cell cycle reentry

**DOI:** 10.1101/250910

**Authors:** Hardik P. Gala, Debarya Saha, Nisha Venugopal, Ajoy Aloysius, Jyotsna Dhawan

## Abstract

Adult stem cells persist in mammalian tissues by entering a state of reversible arrest or quiescence associated with low transcription. Using cultured myoblasts and primary muscle stem cells, we show that RNA synthesis is strongly repressed in G_0_, returning within minutes of activation. We investigate the underlying mechanism and reveal a role for promoter-proximal RNAPol II pausing: by mapping global Pol II occupancy using ChIP-seq, in conjunction with RNA-seq to identify repressed transcriptional networks unique to G_0_. Strikingly, Pol II pausing is enhanced in G_0_ on genes encoding regulators of RNA biogenesis (Ncl, Rps24, Ctdp1), and release of pausing is critical for cell cycle re-entry. Finally, we uncover a novel, unexpected repressive role of the super-elongation complex component Aff4 in G_0_-specific stalling. We propose a model wherein Pol II pausing restrains transcription to maintain G_0_, preconfigures gene networks required for the G_0_-G_1_ transition, and sets the timing of their transcriptional activation.

## Introduction

In mammalian tissues, adult stem cells exist in a state of reversible arrest or quiescence. This “out of cycle” or G_0_ phase is characterized by the absence of DNA synthesis, highly condensed chromatin, and reduced metabolic, transcriptional and translational activity. By contrast to their terminally differentiated counterparts, quiescent cells retain the ability to reenter cell cycle. Indeed, it is this property of reversible arrest that is crucial for stem cell functions such as self-renewal and regeneration. Mounting evidence indicates that the transition into G_0_ includes not only the induction of a specific quiescence program (Coller et al., 2006; Fukada et al., 2007; Liu et al., 2013; Subramaniam et al., 2013), but also suppression of alternate arrest programs such as differentiation, senescence and death (García-Prat et al., 2016; Sousa-Victor et al., 2014). Recent studies highlight the role of metabolic adaptations, epigenetic regulation and signaling mechanisms that enable cells to establish and/or maintain quiescence (reviewed in (Cheung and Rando, 2013)). Collectively, these insights have promoted the view that rather than a passive decline due to the absence of nutrients, mitogens or signaling cues, the quiescent state is actively maintained. Parallels with G_0_ in yeasts (Martinez et al., 2004; Sajiki et al., 2009; Yanagida, 2009) have shown that the quiescence program is evolutionarily ancient (reviewed in (Dhawan and Laxman, 2015)).

Among the adult stem cell populations that are amenable to comprehensive analysis, the skeletal muscle stem cell or satellite cell (Mauro, 1961) holds a special position as it is readily isolated, easily imaged and can be cultured along with its myofiber niche (Bischoff, 1990; Beauchamp et al., 2000; Siegel et al., 2009). However, there is a growing appreciation that perturbing the stem cell niche leads to activation and loss of hallmarks of quiescence, and that freshly isolated stem cells while not yet replicating, have triggered the G_0_-G_1_ transition (Kami et al., 1995; Fukada et al., 2007; Pallafacchina et al., 2010; Zhang and Anderson, 2014; Fu et al., 2015). Thus, culture models employing C2C12 myoblasts that recapitulate the reversibly arrested stem cell state (Milasincic et al., 1996; Sachidanandan et al., 2002; Yoshida et al., 2013) are useful adjuncts for genome-wide analysis of G_0_ as distinct from early G_1_ (Sebastian et al., 2009; Subramaniam et al., 2013; Cheedipudi et al., 2015) that is inaccessible *in vivo*. Importantly, genes isolated on the basis of induction in G_0_ myoblasts in culture mark muscle satellite cells *in vivo* (Sachidanandan et al., 2002; Doles and Olwin, 2015), strengthening their utility as a model of quiescent muscle stem cells.

Several lines of evidence suggest that quiescent cells are held in readiness for cell cycle re-entry by mechanisms that keep the genome poised for action. For example there is an increase in epigenetic modifications (H4K20me3) that promote the formation of facultative heterochromatin and chromatin condensation, and trigger transcriptional repression (Evertts et al., 2013; Boonsanay et al., 2016), while other histone modifications help to maintain the quiescence program (Srivastava et al., 2010; Juan et al., 2011; Mousavi et al., 2012; Woodhouse et al., 2013; Liu et al., 2013). Chromatin regulators induced specifically in G_0_ hold <u>genes encoding</u> key cell cycle regulators in a poised state (Cheedipudi et al., 2015) by preventing silencing. Importantly, epigenetic regulation in G_0_ ensures that the repression of key lineage determinants such as MyoD is reversible, helping to sustain lineage memory (Sebastian et al., 2009).

There is little information on mechanisms that regulate the expression of quiescence-specific genes in the context of global transcriptional repression. The regulation of RNA polymerase II (Pol II) is an integral part of the transcription cycle, with cell type and cell state-specific control (reviewed in (Adelman and Lis, 2012; Puri et al., 2015)). Pol II in quiescent adult stem cells exhibits reduced activation (Freter et al., 2010), Mediator (MED1) is associated with maintenance of quiescence (Nakajima et al., 2013) and transcription factor complex II (TFII) proteins have been implicated in differentiation (Deato and Tjian, 2007; Malecova et al., 2016). However, while components of the Pol II complex contribute to cell state changes, Pol II regulation in reversible quiescence is poorly explored.

In recent years, the view that global regulation of Pol II occurs largely at the level of transcription initiation has undergone a major revision. Elongation control at the level of promoter-proximal Pol II pausing has been implicated in developmental, stress-induced genes, neuronal immediate early genes (IEG) and mitogen-induced activation of groups of genes (Adelman and Lis, 2012; Saha et al., 2011; Levine, 2011; Gaertner et al., 2012). Further, Pol II <u>pausing</u> allows <u>genes to be</u> pois<u>ed</u> for future expression (Kouzine et al., 2013), and appears to mark regulatory nodes prior to a change in developmental state, allowing coordinated changes in gene networks. The temporal dynamics of Pol II pausing during development have been investigated (Kouzine et al., 2013), but the role of Pol II pausing in regulating cell cycle state is not well understood. We hypothesized that withdrawal of cells into alternate arrested states might preconfigure different subsets of genes by Pol II pausing, and that these genes may play critical roles in the subsequent cell fates.

Here, we investigated the role of promoter proximal RNA Pol II pausing in regulating the quiescent state and maintaining G_0_ cells in a primed condition that might enable cell fate transitions. Using ChIP-seq and RNA-seq, we elucidated Pol II occupancy and the extent of transcriptional repression during different types of cell cycle arrest (reversible vs. irreversible), thereby identifying poised transcriptional networks characteristic of G_0_. We used knockdown analysis of G_0_-specific stalled genes identified by our analysis, as well as known regulators of Pol II activity, to delineate the role of <u>pausing</u> in quiescence-dependent functions. Our results suggest a model wherein promoter-proximal Pol II pausing regulates a specific class of repressed genes in G_0_ and creates a state that is primed for cell cycle re-activation. We propose that Pol II stalling may preconfigure gene networks whose repression aids entry into and maintenance of G_0_, and confers appropriate timing to their transcriptional activation during the G_0_-G_1_ transition and subsequent G_1_-S progression.

## Results

### Altered RNA metabolism in quiescent cells: reduced RNA content and low RNA synthesis

Global transcription mediated by all three RNA polymerases is dynamic across the cell cycle (Yonaha et al., 1995; White et al., 1995; Russell and Zomerdijk, 2005), and dampened during cell cycle exit (Hannan et al., 2000; Scott et al., 2001; Russell and Zomerdijk, 2005). To determine if this suppression is common to cells entering alternate states of arrest, we measured total cellular RNA content of C2C12 skeletal myoblasts in different states: proliferating undifferentiated myoblasts (MB), reversibly arrested undifferentiated myoblasts (quiescent/G_0_) and permanently arrested differentiated myotubes (MT). First, using quantitative *in situ* imaging of cells stained with SYTO RNA-select we found that total RNA content per cell was lower in G_0_ than in MB (Figure 1a,b). Interestingly, despite exiting the cell cycle, MT showed no significant reduction in total RNA content, providing evidence that global transcriptional activity distinguishes quiescence from differentiation. A time course of quiescence reversal (30’-18 hr reactivation out of G_0_) showed that a rapid, robust increase in RNA content occurred within 30’ of cell cycle reentry (Figure 1b), and was sustained as cells enter S phase. Flow cytometry of cells stained simultaneously for DNA and RNA confirmed that cells with 2N DNA content in suspension-arrested cultures showed less than half the RNA content per cell than 2N cells in proliferating cultures, validating the results of quantitative imaging and distinguishing between G_0_ and G_1_ (Figure 1c). Flow cytometric analysis of DNA synthesis during reentry is shown for comparison (Figure 1d): cells begin to enter S phase between 12-16 hr after replating. Taken together, the results show that a rapid rise in cellular RNA content occurs minutes after exit from quiescence (marking the G_0_-G_1_ transition) and greatly precedes the G_1_-S transition.

**Figure 1.**
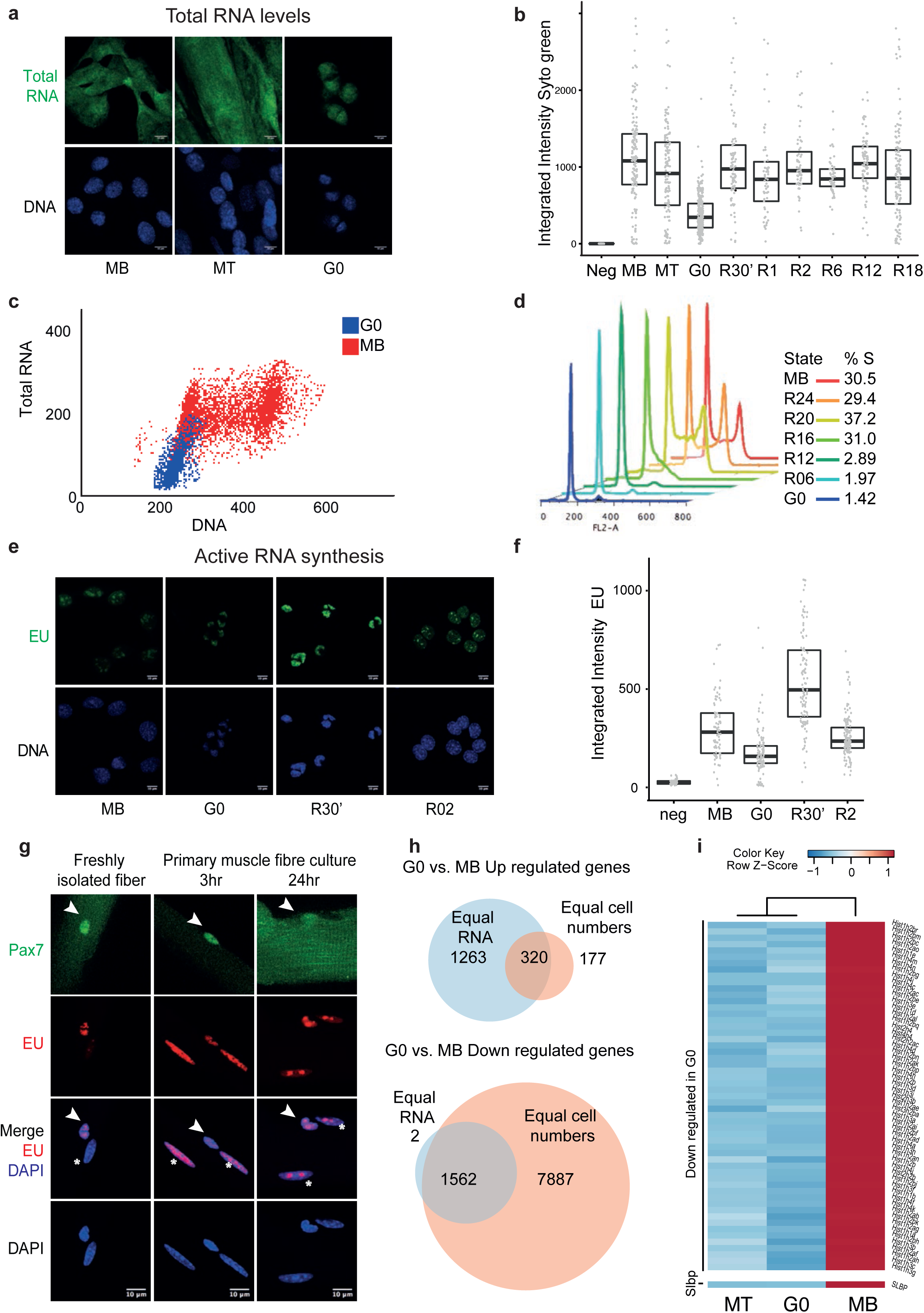
Decreased total RNA content, repressed total RNA synthesis and major down-regulation of the transcriptome in quiescent myoblasts and satellite cells. a. Total RNA in distinct cell states revealed by fluorescence imaging of MB, G_0_ and MT stained for DNA (DAPI) and the RNA-specific dye (SYTO^®^ RNASelect^TM^-green) (scale bar 10μm). All three cellular states show cytoplasmic and nuclear staining. b. Cellprofiler was used to quantitate integrated image intensity values per cell for the 3 cell states as well as across a time course of reactivation from quiescence from 30‵-18hr (R30-R18) (number of cells analyzed >120 for MB, MT, G_0_ and >60 for reactivation time points). The boxplot shows quantification of RNA levels where G_0_ cells show a major reduction of median RNA content compared to MB and MT, and a rapid increase in RNA levels as cells reenter the cell cycle (dark horizontal bar represents median value; the upper and lower limits of box represents 75 and 25 percentiles for each box). G_0_ compared to all other time points shows significance p-value < 2e^−16^ but between MB & MT p-value =0.072 (NS) (Mann-Whitney test). c. Flow cytometric quantification of total RNA (Syto-green) shows reduced RNA content in quiescent (G_0_, blue) compared to proliferative (MB, red) cells. DNA is stained with DRAQ5. Dot plots showing ~50% reduction in RNA in G_0_ compared to G_1_ population of MB. Note that >90% G_0_ cells cluster as a single population with 2N DNA content. d. FACS analysis of cell cycle re-entry from quiescence. Profiles represent a time course of reactivation of asynchronously proliferating myoblasts (MB) out of G_0_ from 30‵-18hr (R30-R18). The numbers represent the % of cells in S-phase. e. Newly synthesized RNA revealed by 5-Ethynyl Uridine (EU) incorporation in a 30‵ pulse: Fluorescence images of MB, G_0_, R30', R2hr: EU (Green) and DNA (DAPI, blue). Scale bar is 10μm. RNA synthesis is low in G_0_ and rapidly enhanced as cells enter G_1_. Note that MB cells display more nucleolar staining, consistent with rRNA synthesis and ribosome assembly, which are reduced in G_0_ and increased again at 30' and 2 hr after reactivation. f. Quantification of EU incorporation by image analysis as in (Number of cells > 80) (a). G_0_ cells display low levels of active RNA synthesis compared to MB. A rapid burst of very strong transcriptional activity within 30‵ of cell cycle activation settles back to levels typical of proliferating cells by 2hr into the G_0_-G_1_ transition. Significant differences were observed (G_0_ compared to MB, R30 and R02: p-value < 2e^−10^, MB / R30 p-value =2e^−14^; but MB / R02 comparison was not significant (Mann-Whitney test). g. Active RNA synthesis in satellite stem cells (SC) associated with single myofibers cultured *ex vivo*. SC marked by Pax7 (green, arrowhead) can be distinguished from differentiated myonucleiwhich are Pax7 negative (MN, asterisk) within the underlying myofiber. In freshly isolated fibers, EU exposure (red) during a 30‵ pulse *ex vivo* shows incorporation only in SC and not in MN, indicating rapid activation of the SC during the isolation protocol. At 3hr and 24hr post isolation, MN also show RNA synthesis to the same level as the SC. h. Transcriptional comparison of G_0_ and MB states using RNA seq shows a major repression of transcript abundance in quiescence. Quantification based on equal cell number (pink) and equal RNA (blue) normalization highlight the bias towards upregulated genes when equal RNA is used for normalization. Overlap of differentially regulated genes between equal RNA (Cufflinks analysis) and cell number normalization methods (DESeq2 package with spike-in RNA controls): size-adjusted Venn diagrams showing overlap of genes between the two normalization methods. Top panel represents up-regulated genes and bottom panel represents down-regulated genes. i. Histone genes are strongly down-regulated in both mitotically inactive states. RNA-seq quantification of 68 histone genes in MB, G_0_ and MT: heat map showing the level of expression for each gene in all the 3 states. The values plotted are row normalized log2 expression scores from cell number normalized datasets. Expression of Slbp a key regulator of replication-dependent histone genes is also repressed in both the arrested states, correlating with repressed histone transcript abundance.

To determine whether reduced steady state levels of RNA in G_0_ result from reduced transcription, we measured active RNA synthesis using pulse-labeling with 5-ethynyl uridine (EU) (Jao and Salic, 2008) (Figure 1e,f). EU incorporation into newly synthesized RNA was strongly suppressed in G_0_ compared to MB and sharply increased within 30’ of activation (Figure 1f) correlating with the rise in RNA content (Figure 1b), and indicating a rapid restoration of the transcriptional machinery concomitant with cell cycle reentry. To evaluate the transcriptional output of muscle stem cells *in situ*, we isolated single myofibers with associated satellite cells and determined their EU incorporation. We found that reversibly arrested satellite cells (SC) *ex vivo* already show robust RNA synthesis within 30’ of isolation, while differentiated myofiber nuclei (MN) in the same sample showed low incorporation of EU (Figure 1g, Figure 1-figure supplement 1a). This observation is consistent with evidence that SC on freshly isolated fibers have already exited from quiescence during myofiber isolation (Kami et al., 1995; Fukada et al., 2007; Pallafacchina et al., 2010; Zhang and Anderson, 2014; Fu et al., 2015; Zammit et al., 2006).

Taken together, the comparison of steady state RNA levels and active RNA synthesis per cell confirm that in G_0_, low global RNA levels result directly from depressed transcriptional activity, which is rapidly reversed in a transcriptional burst within minutes of cell cycle reactivation. Further, since total RNA levels are maintained in irreversibly arrested myotubes, strong repression of RNA biogenesis is not a general function of cell cycle cessation, but rather, a feature of a specific quiescence program.

### Revisiting the quiescent transcriptome using RNA-seq

#### Identifying genes regulated in quiescence using cell number based normalization methods

Previous estimates of altered gene expression in quiescence using differential display (Sachidanandan et al., 2002) in myoblasts, or comparative microarray analysis in a variety of cell types (Venezia et al., 2004; Coller et al., 2006) including myoblasts (Subramaniam et al., 2013) and satellite cells (Fukada et al., 2007; Pallafacchina et al., 2010; Liu et al., 2013) have identified quiescence-induced genes. However, all these studies were based on normalization to equivalent amounts of total RNA which can lead to misinterpretation of global gene expression data (Lovén et al., 2012), particularly when there is sharply varying cellular RNA content across compared samples (as documented above). To quantitatively re-investigate the transcriptome of quiescent myoblasts in contrast to proliferative or differentiated cells, we used RNA-seq analysis, where gene expression was measured by normalization to equal cell number and not equal RNA (enabled by ERCC spike-in analysis as described in Materials and Methods). Post cell number normalization, the similarities between replicates and dissimilarities between samples were computed using Euclidean Distance (Figure 1-figure supplement 1b), confirming that no bias was generated by data processing.

To assess the extent of effects of normalization methods (equal RNA vs. equal cell number) on biological interpretation, we compared gene lists identified as differentially expressed between MB and G_0_ derived from the same RNA-seq dataset but using the two different normalization methods. The Venn diagram (Figure 1h) shows that equal RNA normalization over-represented up-regulated genes compared to the equal cell number normalization (1583 vs. 497) and greatly under-represented down-regulated gene numbers (1564 vs. 9449) (Figure 1 - source data 1). Thus, quiescence is characterized by repression of a much larger number of transcripts than previously appreciated.

#### Repressed RNA biogenesis pathways distinguish quiescent from differentiated cells

To identify distinguishing features between two mitotically inactive states (reversibly arrested G_0_ cells and irreversibly arrested MT), we analyzed gene ontology (GO) terms of differentially regulated genes derived from equal cell number normalization, using gProfiler tools. Consistent with our earlier microarray study (Subramaniam et al., 2013), compared to MB, genes identified as up-regulated in G_0_ are distinct from those in MT (Figure 1-figure supplement 3). GO terms representing response to external stimuli, stress and extracellular matrix organization are over-represented in G_0_, whereas skeletal muscle contraction and ion transport are highlighted in MT. These contrasting terms reflect the suppression of myogenic differentiation in G_0_ myoblasts. Expectedly, the down-regulated genes identified from both mitotically arrested states were enriched for cell cycle processes, DNA replication and cell proliferation (Figure 1-figure supplement 4). Commonly down-regulated gene families that were repressed in both G_0_ and MT included replication-dependent histone transcripts bearing a 3’ stem-loop and lacking a polyA tail (Figure 1i). Slbp which binds the 3’ stem-loop in polyA^-^ histone mRNAs regulating their stability and translation during S phase (Whitfield et al., 2000), was also repressed in both arrested states (Figure 1i).

We focused on the ontologies that were uniquely down-regulated in G_0_ but not in MT, which intriguingly, involved RNA biogenesis itself (tRNA and rRNAs), mitochondria & electron transport chain, proteasome and carbon metabolism. Thus, reversible arrest is uniquely associated with suppression of RNA metabolism, protein turnover and bioenergetics (Figure 1-figure supplement 4). Further, GO terms involved in stem cell proliferation, epithelial differentiation, neurogenesis and cellular growth, were exclusively down-regulated in MT, consistent with the repression of lineage-inappropriate genes in terminally differentiated myotubes (Ali et al., 2008), and retention of progenitor features and plasticity in quiescent myoblasts. Indeed, as has been previously reported, while the satellite cell specification factor Pax7 was induced in G_0_ and repressed in MT, the lineage determinant MyoD was suppressed in G_0_ and induced in MT.

Further, genes identified here in our cultured quiescence model as differentially expressed (quiescent vs. cycling) were corroborated by comparison to genes reported as differentially expressed in freshly isolated quiescent satellite cells (QSCs) vs. activated satellite cells (ASCs) (Liu et al., 2013) (Figure 1-figure supplement 5-a, b). More than 98% of genes expressed in ASCs at 60 hr were expressed in proliferating MB and not in G_0_, consistent with the derivation of C2C12 cells from adult SCs and confirming the global similarity in expression profiles between this cultured cell line and SCs. By contrast with near identity of the ASC-specific gene set, only 17% of the “QSC-specific” gene set of freshly isolated SC were also identified as up-regulated in G_0_ in culture. This observation suggests either that G_0_ *in vivo* employs markedly different programs than *in vitro*, or that the reported “QSC-specific” gene set defines cells that are no longer in G_0_. Taken together with the strong RNA synthesis seen in freshly isolated SC (Figure 1g), this comparative expression data strongly supports the view that disturbance of the SC niche during fiber isolation activates a departure from G_0_. We conclude that the reported “QSC-specific” gene set likely reflects an early-activated cell state, which while still clearly pre-replicative, does not represent undisturbed quiescence.

### The RNA Pol II transcriptional machinery is specifically altered in G_0_

Since we observed strong repression of total RNA levels, reduced RNA synthesis and repression of a large class of Pol II-transcribed genes in G_0_, we hypothesized that Pol II expression itself may also be repressed, to control transcription globally. Indeed, RPB1 the largest subunit of RNA Pol II (N20 pAb) showed decreased levels in both G_0_ and MT (Figure 2a, b, Figure 2-figure supplement 1). Quantitative imaging also revealed that while both G_0_ and MT showed decreased nuclear levels of Pol II (Figure 2b,c), G_0_ cells exhibited more cytoplasmically localized Pol II. Phosphorylation of the CTD of RNA Pol II marks distinct stages in the transcriptional activation cycle, where Ser5-p and Ser2-p reflect initiation and elongation stages respectively (Zhang et al., 2012). G_0_ cells displayed the lowest level of active Pol II marks compared to MB and MT (Figure 2b, c), indicating that overall Pol II processivity in the 3 states would be G_0_ < MT < MB. Taken together, these observations show that despite reduced Pol II levels in both mitotically inactive states, MT exhibit a nuclear localized enzyme which is transcriptionally active, whereas G_0_ cells show predominantly cytoplasmically localized Pol II, and is transcriptionally less active, consistent with reduced EU incorporation.

**Figure 2.**
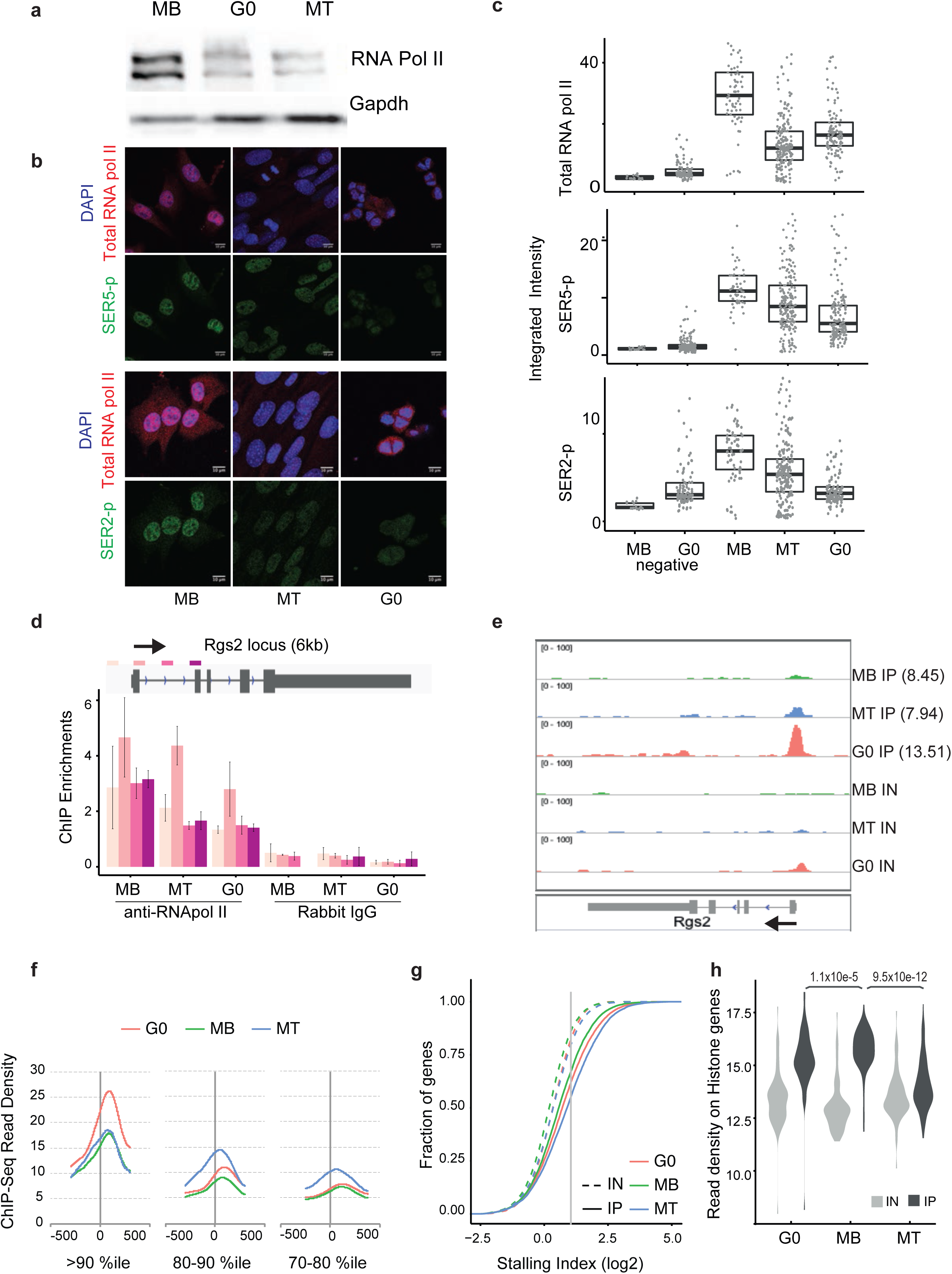
Quiescence is associated with reduced RNA Pol II levels and promoter-proximal RNA Pol II stalling at a specific set of loci. a-c Reduction in levels of RNA Pol II in G_0_ and MT: a. Western blot analysis using antibodies against RNA Pol II large subunit (total) shows two distinct bands (~250kd) corresponding to hyper-phosphorylated and hypo- phosphorylated RNA Pol II isoforms. The cell cycle arrested states G_0_ and MT show reduced levels of RNA Pol II. GAPDH was used as loading control. b. Expression and localization of RNA Pol II: Representative immunofluorescence images for MB, G_0_ and MT stained for total Pol II (large subunit, red) and active Pol II (Ser2-p [green], upper panel) or total Pol II (red) and Ser-5p [green], lower panel). Scale bar is 10μm; number of cells analyzed > 70 for each cell state. MB and MT show nuclear localization of total and active Pol II whereas G_0_ cells show cytoplasmic staining. c. Cell profiler was used to quantitate integrated image intensity values per nucleus for all three states and corresponding negative controls. The boxplot shows reduced levels of Pol II and its active modification marks-representing both initiating (Ser5-p) and elongation (Ser2-p) in G_0_ state. For total RNA pol II levels p-value < 2e^−7^ for all pairwise comparisons. For Ser2-p, G_0_ and G_0_ negative control are not significantly different; whereas p-value is <2e^−7^ when compared pairwise with MB, MT and G_0_ states. For Ser5-p negative control (MB and G_0_) are not significantly different; whereas MB and MT p-value <0.00036 and G_0_ compared to MB/MT p-value < 2e^−7^ (Mann-Whitney test). d, e. RNA Pol II enrichment on the TSS of the Rgs2 locus, a gene up-regulated in G_0_. (d) Targeted ChIP q-PCR, enrichment is represented as percentage of input using primers tiling Rgs2 gene in RNA Pol II IP (N20 pAb) and IgG control. Schematic of the Rgs2 locus is shown above the graph, depicting location of primers upstream (-530), TSS (+100), gene body-1 (+500) and gene body-2 (+1500) used for ChIP-qPCR. Values represent mean + SD n=3, p-value < 0.01. e. ChIP-seq depicted by a genome browser snapshot for Rgs2 (locus is oriented with TSS on right). The read density is normalized per million within each of the sample and plotted at equal scale. The numbers in the brackets indicates the log2 expression score from the cell number normalized RNA-seq dataset. f. Quiescent cells show high enrichment of RNA Pol II on a subset of genes compared to MB and MT. The cumulative density of sequencing reads plotted across +/− 300 bases relative to TSS. <u>RNA pol II enrichment d</u>ensities <u>are represented for</u> genes <u>grouped b</u>as<u>ed on top 3 decile values (ie 1517 genes for each of 70-80, 80-90, 90-100 percentiles group)</u> for each state are plotted. Note the substantial difference in peak height around TSS between G_0_ and MB/MT for the >90th percentile gene set but equivalence in the 70-90 percentile gene set. g. Both mitotically inactive states (G_0_ and MT) show higher stalling indices compared to proliferating MB. CDF plot for stalling indices of both IP and IN samples for the three cellular states are depicted. All pairwise comparisons show p-value <2e^−16^ (except comparison between Inputs G_0_ and MT not significant (Mann-Whitney test). h. RNA Pol II is enriched on histone genes in G_0_ but not in MT. Violin plots of the read density across 68 histone genes show specific enrichment in all three IP samples over the respective input samples. With respect to the corresponding input sample, enrichment is highest in MB (p-value < 10e^−12^), retained at near equivalent levels in G_0_ (p-value < 10e^-11^) and significant but much lower in MT (p-value < 10e^-5^) (Mann-Whitney test).

### Identification of stage-specific Pol II recruitment in myogenic cells

To identify stage-specific programs of gene activation we used Pol II enrichment profiling. We first validated the N20 antibody using targeted ChIP analysis of the Rgs2 locus (called as G_0_-induced by RNA-seq (Figure 1 - source data 1) and by microarray analysis (Subramaniam et al, 2013)), interrogating enrichment at 4 locations across the gene using qPCR (Figure 2d). Specific enrichment of Pol II across the Rgs2 gene was detected in all 3 cellular states, with peak enrichment at the TSS.

To estimate the global recruitment of Pol II on the promoters of all genes, we used ChIP-seq. Libraries were prepared as detailed in Materials and Methods from duplicate samples of MB, G_0_ and MT (Figure 2-figure supplement 2, 3). Only non-overlapping genes of length >2500 bases were used for this analysis (15,170 annotated mouse genes from mm9 build). A snapshot of the Rgs2 locus corroborates the finding from q-PCR based experiments confirming enrichment at the TSS (Figure 2e). Promoter-proximal engagement of Pol II on the Rgs2 gene was strongest in G_0_ correlating with its high expression. Pol II enrichment at the TSS in all 3 states (Figure 2-figure supplement 4) is consistent with the “toll booth” model (Adelman and Lis, 2012) where initiated Pol II may pause, and represents a checkpoint for transcription, presaging stage-specific regulation of elongation.

To compare global cell state specific variations in Pol II recruitment, we <u>first categorized non-overlapping genes in decile groups based on</u> enrichment <u>score</u>, separately for each of the three conditions<u>, and further plotted the normalized enrichment of RNA Pol II around the TSS</u> (Figure 2f). In all cases, enrichment peaked within 30-100 bases downstream of the TSS, consistent with initiated but paused polymerase complexes. Strikingly, the recruitment of Pol II on promoters in G_0_ cells for the >90th percentile Pol II-enriched genes was significantly higher than that of either MB or MT, which were comparable. By contrast, profiles for the 70-90 percentile sub-groups behaved similarly for all 3 cellular states. Thus, a small group of promoters in G_0_ are occupied with much higher abundance of Pol II than other genes, and may represent a G_0_-specific class of genes, with elevated ongoing Pol II recruitment. Since promoter occupancy is observed not only on genes that are highly expressed (“active”), but also on genes that are not expressed and exhibit <u>paus</u>ed polymerases (“poised”), we used the extent of promoter clearance to distinguish these classes.

#### Calculation of polymerase stalling index (SI): more genes are stalled in G_0_ and MT than MB

Promoter clearance can be quantified using the distribution of Pol II abundance across an entire locus. The ratio of Pol II enrichment at the promoter region to that across the gene body region is defined as the Pausing Index or Stalling Index (SI), where higher SI reflects a lower promoter clearance of Pol II and indicates a point of regulation (Zeitlinger et al., 2007; Min et al., 2011; Danko et al., 2013). All genes with annotated gene length >2500 bases were used for further analysis. Genes overlapping with neighboring genes were removed from the analysis. Pol II density is calculated by base pair enrichment in ChIP-seq data, at promoters (-500 to +300 relative to TSS) and gene body (+300 to +2500).

To confirm that the SI is a function of specific enrichment of Pol II, we calculated stalling indices similarly for control input samples: each IP sample exhibited significantly more enrichment/stalling than that of its respective input (Figure 2g, Figure 2-figure supplement 5). Interestingly, >50% of analyzed genes display a SI >1 in any state, confirming that clearance of engaged Pol II from the promoter is a significant common regulatory mechanism. Comparative GO analysis for cell state-specific stalled genes (SI>2.5 in MB, MT and G_0_) shows that metabolic and stress response genes are stalled in all three states (Figure 2-figure supplement 6). Strikingly, the proportion of genes that are stalled in the two non-dividing states (G_0_ and MT) is higher than that in proliferating cells, suggesting that Pol II stalling represents a point of regulation associated with mitotic inactivity. Thus, halted cell division <u>(either due to quiescence or differentiation)</u> is associated with higher rates of polymerase pausing.

#### Short repressed replication-dependent histone genes exhibit Pol II occupancy in G_0_ but not MT

The histone genes transcribed by Pol II were excluded from stalling index calculation, as they do not meet the length criteria set for the analysis. This important gene family comprises ~80 genes that map to chr13 and chr3 in clusters, with a few genes singly located on other chromosomes. To evaluate the relative transcriptional engagement of histone loci, the read density of Pol II across the entire gene region was used. Notably, strong enrichment of Pol II was observed in both MB and G_0_ but not in MT (Figure 2h, Figure 2-figure supplement 7), indicating that the histone loci are transcriptionally silent in terminal differentiation, but not in reversible arrest. Thus, <u>RNA Pol II occupancy on histone gene loci is indicative of transcriptional activity but</u> the low mRNA levels in G_0_ (Figure 1i) likely reflect <u>rapid</u> post-transcriptional <u>turnover</u>, and the sustained Pol II engagement across the entire histone gene family represents a primed gene network preconfigured for the return to proliferation.

### G_0_-stalled genes are specifically repressed in quiescence

Since promoter proximal stalling influences the transcriptional output from a given locus, the repertoire of genes regulated by this mechanism in a given state would yield insights into global Pol II-mediated regulation of that state. Our hypothesis was that control of quiescence might include uniquely stalled genes. Therefore, we analyzed the group of 727 genes that show strong stalling (SI > 2.5) in G_0_ (hereafter referred to as ‘G_0_-stalled genes’ (Figure 2 - source data 1)). The cumulative distribution of SI for G_0_-stalled genes was compared across the three states (SI>2.5 in G_0_ is shown in Figure 3a). Genes that were strongly stalled in G_0_ (high SI) display much lower SI in MB and MT states, indicating that they experience enhanced stalling specifically in quiescence (Figure 3a, Figure 3-figure supplement 1).

**Figure 3.**
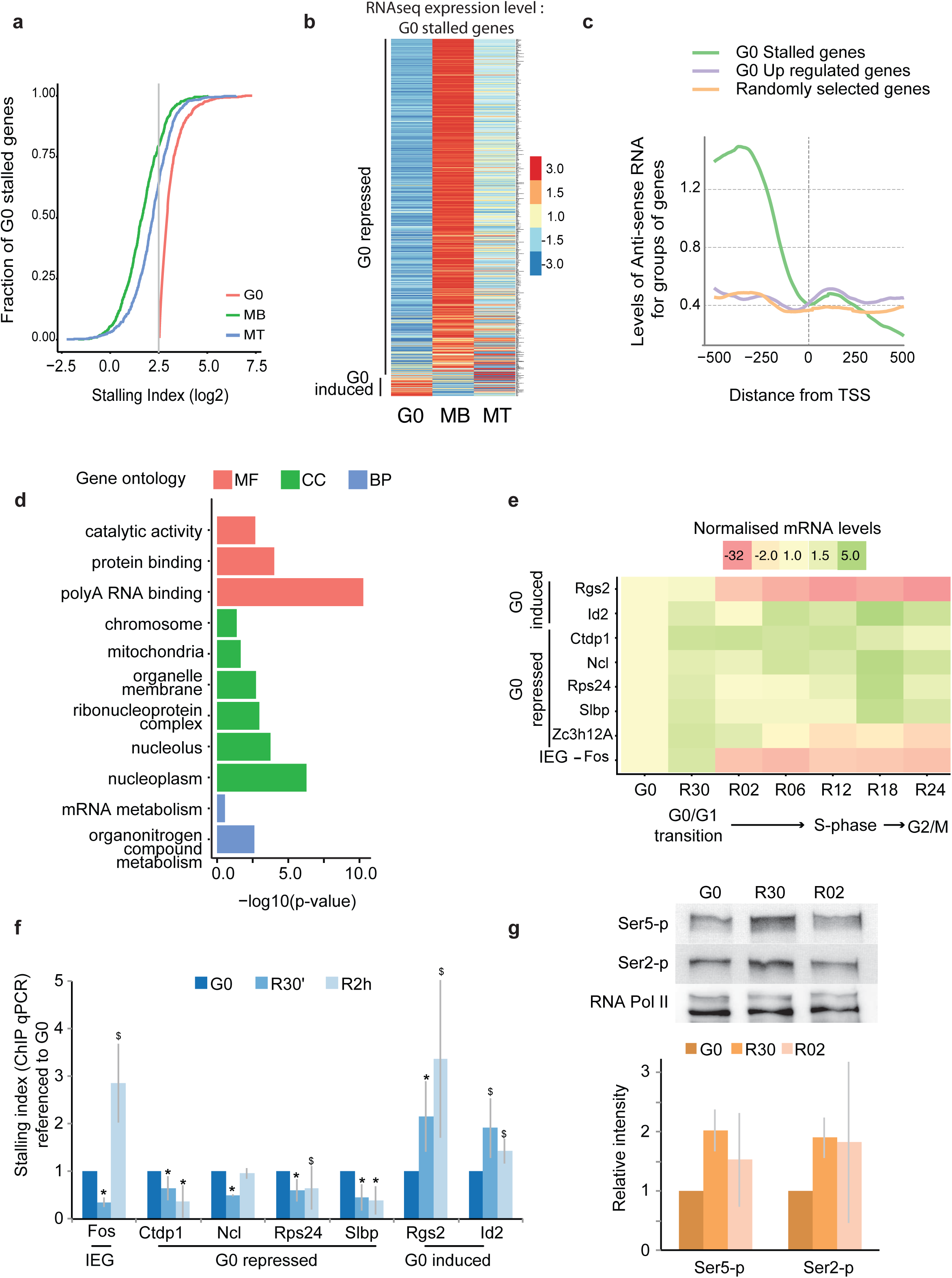
G_0_-specific stalled genes are repressed in quiescence and poised for activation. a. The subset of 727 genes which showing stalling index (SI) >2.5 in G_0_ were selected and cumulative distribution of stalling index is represented. These genes show G_0_-specific stalling since more than 60% of these genes display SI < 2.5 in other states (MB and MT). b. Heat map representing mRNA expression as enumerated from RNA-seq data for genes exhibiting Pol II stalling specifically in G_0_. Note that expression of only ~5% of these genes are induced in G_0_ compared to MB while 95% of stalled genes are repressed in quiescence. c. <u>Normalized read density of antisense RNA flanking +/-500 bases with respect to TSS for G0 state. Equal number of genes (n=410) were used for each of three categories: (i) up-regulated in G0 vs. MB, (ii) G0 stalled genes and (iii) random selected genes as control.</u> <u>d. G_0_-stalled genes were analyzed by PANTHER over-representation test and significantly enriched ontologies (BP – Biological process, CC – cellular component, MF – molecular function) with p-value <0.05 are shown. Notably, ontologies for RNA binding, mitochondria, ribonucleoprotein complexes and mRNA metabolism are enriched in this set of G_0_-stalled genes.</u> e. Quantitative RT-PCR analysis of G_0_-stalled genes in quiescence and during cell cycle re-entry. The expression in reactivation time points is <u>normalized</u> to <u>expression in</u> G_0_, and mean relative expression level is plotted with increasing time as cells exit quiescence (n=3). Fos, a canonical IEG is transiently induced only at 30‵ after reactivation and rapidly repressed. G_0_-stalled but expressed genes Rgs2 (which is repressed during entry) and Id2 (which continues to express as cells exit quiescence). All G_0_-stalled and repressed genes (Ctdp1, Ncl, Rps24, Slbp & Zc3h12A) show induction within 30’ after cell cycle activation<u>, albeit</u> with differential kinetics <u>at later points</u> during G_1_-S progression. f. Stalling index of G_0_-stalled genes decreases during cell cycle reactivation and is associated with expression dynamics. Calculation of stalling index by ChIP-qPCR was done using ratio of ChIP enrichment for primers targeting TSS vs. gene body (+500-1000 bps) in each time point: G_0_, R30’ R2h. Both IEG and G_0_-stalled genes show rapid decrease in stalling index at 30‵. G_0_-expressed genes Rgs2 and Id2 show gradual increase (Rgs2) and no change (Id2) in stalling index respectively, correlating with expression dynamics (significance calculated using Students t-test * = p-value <0.05 and $ = p-values between 0.05-0.15). g. Western blots for total RNA Pol II and active phosphorylated forms at G_0_, and 30’-2hr after reactivation. <u>Densitometric</u> analysis shows an increased proportion of ser2-p/ser5-p modification at R30 and R2 compared to G_0_, correlating with activation. Values represent mean + SD, n=3, ^*^p-value<0.05.

Expression levels enumerated by RNA-seq for G_0_-stalled genes showed that ~95% of these genes were specifically repressed in G_0_ (Figure 3b, Figure 3-figure supplement 2). We conclude that most of the 727 genes <u>show very low</u> transcriptional activ<u>ity</u> in G_0_ but continue to display high promoter proximal Pol II occupancy, indicative of a poised or primed state. Most of these genes are also repressed in MT but do not exhibit stalled Pol II, indicating that other mechanisms are responsible for their reduced expression in differentiated cells. <u>Polymerase pausing has previously been associated with upstream divergent transcription (Flynn et al., 2011). Therefore, we evaluated the level of antisense RNA associated with promoters of genes categorized as G_0_-stalled, G_0_ up-regulated (compared to MB) or a randomly selected group containing an equal number of genes. Interestingly, upstream antisense transcription was observed only for G0-stalled genes</u> (Figure 3c). Gene ontology-based clustering using DAVID revealed distinct terms encompassing G_0_-stalled genes (Figure 3d). These include Pol II-transcribed genes involved in ribosomal machinery, mRNA biogenesis & RNA processing, as well as mitochondria-related genes. <u>We conclude that RNA-pol II pausing is associated with high rates of antisense transcription, and distinguishes specific gene groups in G0</u>.

### Genes that display stalled polymerases in G_0_ are poised for reactivation during G_1_

<u>Considering that quiescent cells display reduced levels of total RNA, the enrichment of GO terms relating to ribosomal machinery and RNA processing genes (Fig.1) led us to hypothesize that the genes regulated by RNA Pol II pausing might be key nodes for global control of RNA metabolism.</u> To <u>validate this</u> promoter proximal stalling, we chose genes involved in RNA regulation <u>which</u> include<u>s</u> genes involved in rRNA biogenesis (nucleolin -Ncl) and ribosomal protein S24 -Rps24), Pol II regulation (Pol II recycling enzyme CTD phosphatase -Ctdp1), and post-transcriptional regulators (stem loop binding protein –Slbp, that controls histone mRNA stability, and Zinc Finger CCCH-Type Containing 12A -Zc3h12A), an RNase that modulates miRNA levels). Enhanced stalling is confirmed as depicted on genome browser snapshots of Pol II occupancy on these genes (Figure 3d, Figure 3-figure supplement 3,4,5,6). <u>Further, all these G_0_-stalled genes are marked by histone modifications typical of active (H3K4me3) and not repressive chromatin (H3K27me3)</u> (Figure 3-figure supplement 7).

<u>Since these genes display active histone marks but are not actively expressed in G0</u> (Figure 3-figure supplement 7), <u>we</u> hypothesi<u>zed</u> that <u>RNA</u> Pol II stalling in G_0_ may poise genes for subsequent activation. Comparable RNA-seq data was not available for reactivation, so we used q-RTPCR to evaluate induction of specific genes during cell cycle reentry. Within the first 30’ of reactivation, we observed a sharp increase in normalized mRNA levels (referenced to G_0_) of the immediate early gene Fos, a hallmark of the G_0_-G_1_ transition (Kami et al., 1995). Id2 (stalled but active in G_0_) was further induced during re-entry. By contrast, the G_0_-stalled active gene Rgs2 showed a rapid decrease in expression during re-entry. <u>Interestingly, only the</u> G_0_-stalled, repressed genes (Ncl, Rps24, Ctdp1, Zc3h12A, Slbp) showed enhanced expression (Figure 3e) <u>whereas genes repressed but not stalled (MyoD, L7, and YPLE5) failed to show rapid activation upon cell cycle re-entry</u> (Figure 3-figure supplement 8). <u>Further, t</u>he increased level of transcripts associated with exit from quiescence reflects in increased protein levels (Ncl and Rps24; Figure 3-figure supplement 9 and 10) <u>in both C2C12 model and primary satellite cells</u>. These data confirm that G_0_-stalled, repressed genes are functionally activated during the G_0_-G_1_ transition. Together, these patterns of expression suggest that while these G_0_-stalled genes are differentially regulated at later stages of the cell cycle, their common early reactivation is consistent with the hypothesis that reversal of stalling contributes to their coordinate restoration to transcriptional competence.

### Transcriptional induction during G_0_-G_1_ is associated with reduced stalling

Since expression of most G_0_-stalled genes was low in quiescence and induced during cell cycle reentry, one might expect a corresponding decrease in the proportion of promoter-proximal stalled Pol II, or an increase in the proportion of elongating Pol II (gene body). We investigated potential changes of Pol II occupancy from G_0_ to early G_1_ (30’-2 hr, the window of induced expression), by computing SI for each candidate gene, using targeted ChIP-qPCR. Consistent with their identification by ChIP-seq as G_0_-stalled, the SI calculated by this targeted analysis was also highest in G_0_, and decreased within 2 hr of exit from quiescence, indicative of increased promoter clearance as cells enter G_1_ (Figure 3f). The IEG Fos, that exhibits strong stalling in G_0_, registered a rapid drop in SI in the first 30’ after activation, at a time when its mRNA expression is high (Figure 3e), consistent with a rapid increase in promoter clearance. All the selected G_0_-stalled repressed genes showed similar trends. Notably, the Rgs2 and Id2 genes (G_0_ stalled but active) showed the opposite trend: the SI increased as cells exited quiescence, suggesting altered regulation at the level of pausing for repressed vs. active genes. Collectively, this analysis confirms that activation of G_0_ stalled genes is accompanied by reduced stalling during the early G_1_ transcriptional burst.

### Alterations in RNA Pol II phosphorylation during the G_0_-G_1_ transition

To analyze whether changes in global transcriptional activity of Pol II across cell states are associated with global alteration in its phosphorylation status, we used western blot analysis for total Pol II and its actively transcribing phosphorylated forms (Ser5-p and Ser2-p; Figure 3g), and calculated the ratio of phospho-Pol II to total Pol II in each state (Figure 3g). At R30’ we observed a sharp increase in the proportion of Ser5-p- and Ser2-p-modified polymerase, indicating a rapid rise in transcriptional competence compared to G_0_ & correlating well with the rapid rise of total RNA and active synthesis shown in Figure 1b and Figure 1f, which stabilizes in 2 hr.

Taken together, we conclude that these repressed G_0_-stalled genes undergo release of Pol II stalling upon exit from quiescence that correlates with their rapid induction during this transition. Further, the data so far suggest that the release of Pol II stalling is associated with exit from quiescence, and this mode of regulation facilitates cell cycle re-entry by controlling G_0_-stalled genes encoding regulators of the protein synthesis machinery, mRNA biogenesis and transcriptional control.

### RNA Pol II stalling mediates repression that controls the quiescent state

At the level of gene expression, the functional consequences of Pol II stalling in G_0_ are repression of transcription (resulting in depressed cellular activity), while stalled loci are poised for activation in response to appropriate signals. Our observations thus far identified several G_0_-specific stalled genes with the appropriate functions to participate in the biology of quiescence and activation, but whether Pol II stalling is directly responsible for establishment or maintenance of the quiescent state has not been demonstrated. Since Pol II pausing on G_0_-stalled genes results in lower mRNA production from the stalled transcription unit, we directly decreased the expression level of individual G_0_-stalled genes in proliferating MB and analyzed the functional consequences for entry into quiescence. We used siRNA-mediated knockdown of the selected G_0_-stalled genes (Ctdp1, Ncl, Rps24, Slbp, Zc3h12A) in cycling MB (Figure 4-figure supplement 1) and evaluated effects on RNA synthesis, DNA synthesis and the cell cycle.

Knockdown cells were first examined for alterations in total RNA content and active RNA synthesis using SYTO-RNASelect staining and EU incorporation respectively (Figure 4a, Figure 4-figure supplement 2). Total RNA content was decreased in Ncl KD and increased for Zc3h12A KD compared to control siRNA treated cells, but not appreciably altered in Ctdp1, Rps24 or Slbp KD (Figure 4a, Figure 4-figure supplement 2). RNA synthesis was decreased by knockdown of four genes consistent with their roles in rRNA biogenesis (Ncl & Rps24), regulation of mRNA synthesis (Ctdp1) and miRNA-mediated mRNA turnover (Zc3h12A). Interestingly, nuclear area was decreased in Ncl & Ctdp1 siRNA conditions, consistent with the increased chromatin compaction seen in G_0_ (Evertts et al., 2013) (Figure 1a, e). These observations demonstrate that in proliferating cells, repression of key genes that normally experience a G_0_-specific stalling, is sufficient to trigger a quiescence-like state as assessed by reduction in RNA content, active transcription and nuclear area.

**Figure 4.**
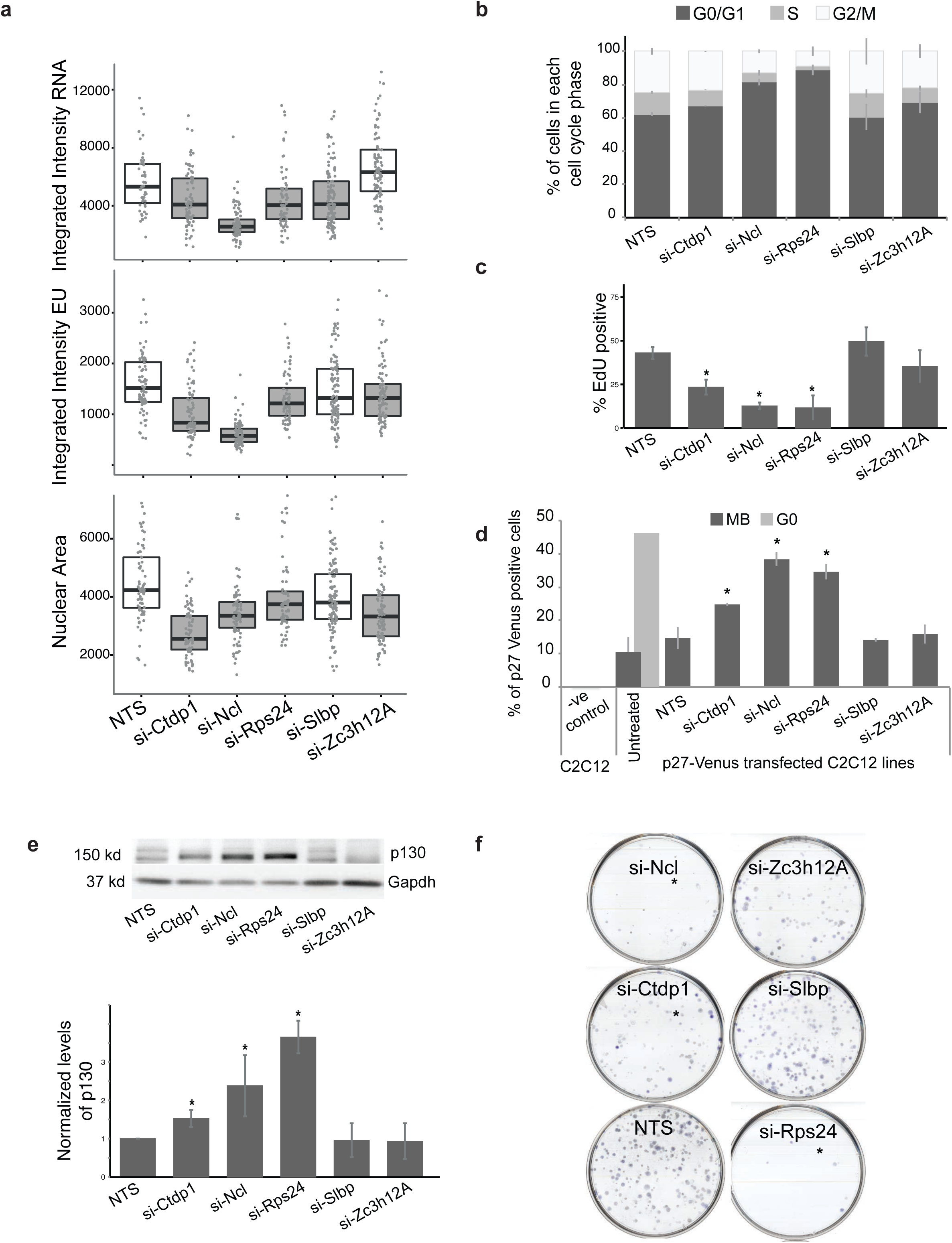
Perturbation of G_0_-stalled genes compromises cell cycle and self-renewal. a. <u>Quantification of integrated image intensity values per cell (number of cells > 75 per condition) to estimate effect of siRNA treatment on total RNA, EU incorporation and nuclear area. Decreased total RNA was observed for Ncl (2.6e-16), Ctdp1, Rps24 and Slbp (< 0.01). Decreased RNA synthesis was observed for Ncl & Ctdp1 (p <1e^-7^), Rps24 (0.0007) and Zc3h12A (0.0128). Nuclear area was decreased for Ctdp1, Ncl and Zc3h12A (<1 x e^-7^), Rps24 (0.0044). (N=3, using Mann-Whitney test, gray shaded box represents p-value < 0.05).</u> b. <u>Change in cell cycle due to the knockdown of G_0_-stalled genes. The stacked bar plots for the proportion of cells in each stage of the cell cycle (G_0_/G_1_, S, G_2_M) for cells treated with siRNA in proliferative conditions. Increase in G_0_/G_1_ phase was observed for siRNA treatments of Ctdp1, Ncl and Rps24 compared to NTS control. The overlay cell cycle traces and statistics are shown in</u> Figure 4-figure supplement 3. c. Proliferation measured by EdU incorporation after knockdown: knockdown of Ctdp1, Ncl and Rps24 (* = p-value <0.05, n=3 t-test) results in cell cycle blockade- increased 2N population and decreased EdU incorporation. d. <u>Flow cytometry shows that %p27m-Venus positive cells increase in proliferating MB treated with siRNA for G_0_-stalled genes. Untransfected C2C12 lines are used as negative control (Figure 4-figure supplement 4). Control cells (p27-mVenus stable C2C12 line) show increased proportion of p27+ cells when shifted to quiescence-inducing conditions (G_0_ ~ 40%) compared to proliferating (MB ~ 10 %), confirming p27 as a marker of quiescence. Further, knockdowns of Ctdp1, Ncl, Rps24 show significantly increased p27+ cells compared to NTS control, indicating that forced reduction in mRNA levels of G0 stalled genes in proliferating conditions is sufficient to induce features of quiescence.</u> e. <u>p130 protein level increases (estimated using western blots) in proliferating MB treated with siRNA for G_0_-stalled genes. The knockdown of Ctdp1, Ncl, Rps24 shows significant increase in quiescence marker P130 and graph represents the normalized densitometry levels. (*= pvalue <0.05, N=3 compared to NTS).</u> f. Knockdown of selected G_0_-stalled genes affects self-renewal: colony-forming ability for Ncl, Rps24 and Ctdp1 knockdown is reduced. Quantification is shown in (Figure 4 -figure supplement 1 - S4b (Values represent mean + SD, p value <0.05, n=3)

The cell cycle in knockdown and control cells was analyzed using flow cytometry and EdU incorporation (Figure 4b,c and Figure 4-figure supplement 3). Of the 5 G_0_-stalled genes analyzed, knockdown led to reduced EdU incorporation and G_1_ arrest for the same 3 genes (Ncl, Ctdp1 & Rps24) that affected RNA synthesis, while 2 (Slbp & Zc3h12A) did not show significant changes in cell cycle profile (Figure 4b,c). As these genes regulate essential processes, there is ~40% decrease in viability in for Rps24 and ~20% decrease in Ctdp1 and Ncl siRNA treated cells compared to ~10% decrease in control siRNA treated cells (Figure 4-figure supplement 4).

<u>We directly tested the onset of quiescence in cells knocked down for G_0_-stalled genes, by evaluating expression of quiescence specific markers p27 (cyclin-dependent kinase inhibitor 1B)</u> (Oki et al., 2014) <u>and p130 (Rb family tumor suppressor)</u> (Carnac et al., 2000; Litovchick et al., 2004) (Figure 4-figure supplement 5,6). <u>Knockdown of the same 3 genes (Ctdp1, Ncl and Rps24) that showed reduced EdU incorporation, also showed a significantly increased proportion of p27^+^ cells, comparable to the increase observed when proliferating cells are shifted to quiescence-inducing conditions</u> (Figure 4d). <u>Similarly, p130 protein was induced when Ctdp1, Ncl or Rps24 were knocked down in proliferating myoblasts</u> (Figure 4e). Taken together, our observations suggest that repression of these G_0_-stalled genes is essential for quiescence, since forced reduction of their expression in proliferating cells leads to <u>induction of markers of G0, indicative of a quiescence-like state</u>.

### G_0_-stalled genes contribute to self-renewal

To investigate the role of these G_0_ stalled genes in self-renewal, we evaluated the effect of knockdown on colony formation after exit from quiescence. G_0_ cells display enhanced self-renewal compared to cycling cells, supporting the view that the quiescence program includes a self-renewal module (Collins et al., 2007; Subramaniam et al., 2013). Knockdown of the same three genes that affected RNA synthesis and G_1_ blockade in proliferating cells (Ctdp1, Ncl, Rps24) also showed a decrease in colony forming ability compared with control cells (Figure 4f, Figure 4-figure supplement 7). Since equal numbers of viable cells were plated for the CFU assay, the effect of knockdown on genes that are normally induced very rapidly (30’-2 hr) would be to dampen their induction upon G_1_ entry, accounting for the sharp decrease in colony formation. This finding further demonstrates the importance of timely reactivation of G_0_-stalled genes during the G_0_-G_1_ transition, failure of which compromises self-renewal.

We conclude that repression of key cellular processes identified by analysis of G_0_-specific Pol II stalling is sufficient to cause cycling cells to <u>acquire</u> quiescence-like <u>features</u>. Consistent with the pre-existing transcriptional repression of G_0_-stalled genes, further repression by siRNA knockdown does not appear to affect the quiescent state itself, but failure to de-repress their expression upon exit (normally mediated by release of RNA Pol II stalling) results in attenuated cell cycle re-entry and diminished self-renewal. Collectively, our results demonstrate that Pol II undergoes a stage-specific regulation to stall on key genes that are required to poise quiescent cells for efficient cell cycle re-entry.

### Aff4, a component of the Pol II super-elongation complex regulates G_0_-stalled genes

Release of paused Pol II is mediated by promoter engagement of P-TEFb, a complex comprised of CDK9 and cyclin T. P-TEFb availability and recruitment is regulated by Hexim1, Aff4, and Brd4, with direct consequences on Pol II elongation (Adelman and Lis, 2012; Jonkers and Lis, 2015). To determine if these key regulators of elongation affected expression of G_0_-stalled genes, we used siRNA treatment (in G_0_) (Figure 5-figure supplement 1), and <u>investigated</u> deregulation of G_0_-stalled genes. While, Hexim1 knockdown led to down-regulation for 4 of 5 selected (all except for Ncl) <u>and</u> Brd4 siRNA led to up-regulation of Ncl and Slbp<u>, unexpectedly, knockdown of Aff4 led to significant up-regulation of all 5 selected G_0_-stalled genes</u> (Figure 5a and Figure 5-figure supplement 2-a,b). Thus, Aff4 is the Pol II elongation control factor that commonly restrains expression of stalled IEG in G_0_ since its knockdown stimulated their transcription.

**Figure 5.**
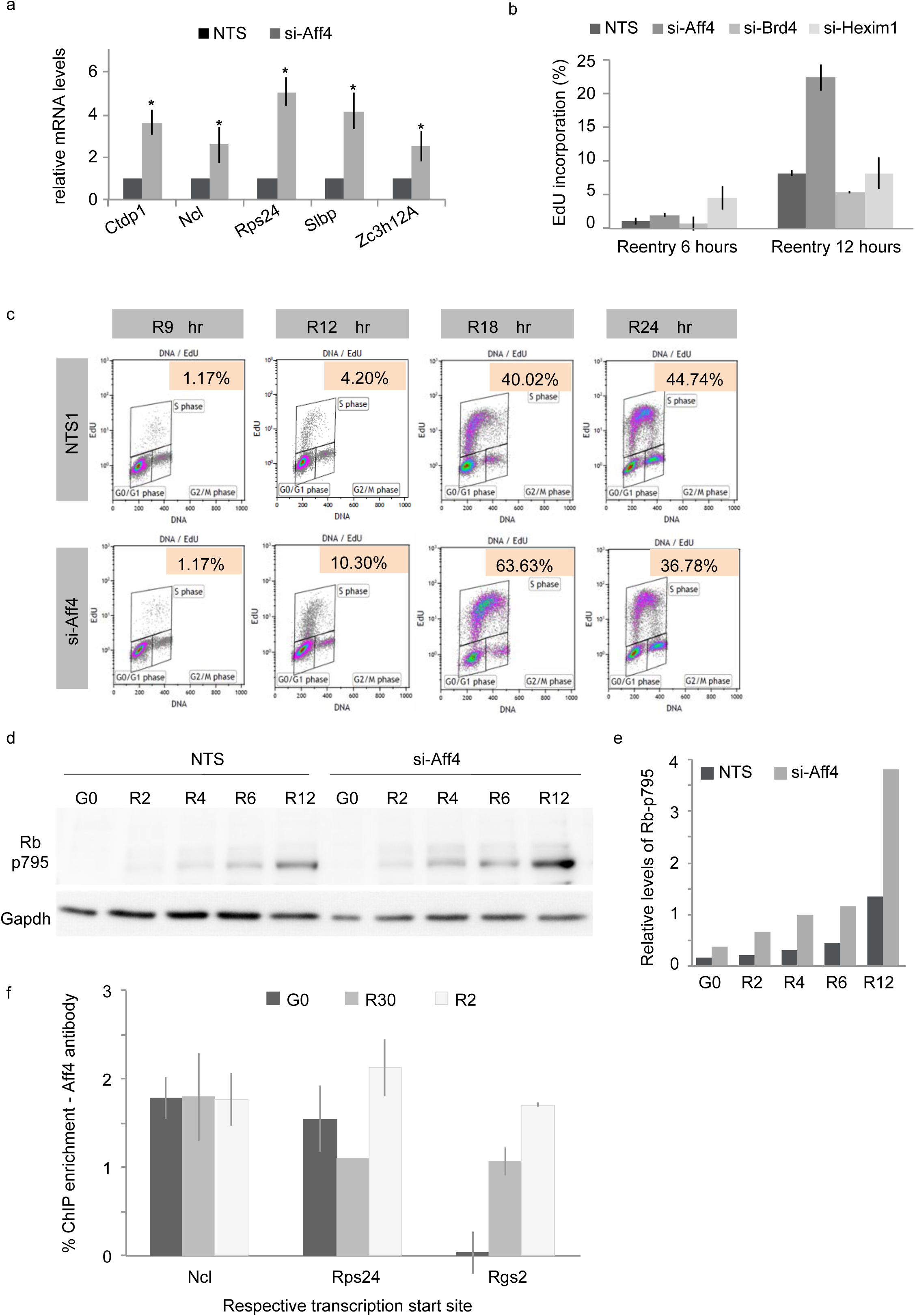
Aff4, a regulator of RNApol II stalling regulates G0 stalled genes and the G0-G1 transition: an unexpected negative role for the SEC complex member. <u>a. Perturbation of Aff4, a regulator of Pol II stalling leads to de-repression of G_0_-stalled genes. qPCR analysis of expression of G_0-_stalled genes after Aff4 siRNA treatment of quiescent cells shows significantly increased mRNA expression compared to NTS treated G_0_ cells (Values represent mean + SD n=3, p value <0.05).</u> <u>b, c. Among regulators of pol II elongation, Aff4 uniquely affects exit from quiescence. Functional effect of knockdown of RNA Pol II regulators on the kinetics of S-phase entry from G_0_: EdU incorporation was estimated at 6hr and 12hr after cell cycle activation (R6, R12). Aff4 KD at R12 shows significantly increased EdU incorporation compared to NTS treated cells. Brd4 KD and Hexim1 KD cells show no difference. (Values represent mean + SD, n=3, p-value * <0.05). Horse-shoe plots of EdU-pulsed re-entry time-course accurately estimates % of S-phase cells in Aff4 knockdown using flow cytometry for cells co-stained for DNA (x-axis) and EdU (y-axis). % of S-phase cells (inset text) is significantly higher at 12-18hr in the Aff4 knockdown, showing more rapid S phase kinetics (quantification depicted in Figure 5-figure supplement 3).</u> <u>d, e. Confirming accelerated cell cycle re-entry in Aff4 knockdown cells compared to NTS control. The early marker for G0-G1 transition -phospho-Rb (S795)- is activated earlier and to higher levels than in control cells indicating that Aff4 knockdown speeds up reactivation. d is representative western blot for p130 and GAPDH (control gene) and e show the normalized densitometry levels of p130 for NTS/ siAff4 treated samples during the quiescence and re-entry time-course.</u> <u>f. Aff4 controls the expression of G_0_-stalled genes during quiescence and reactivation. Aff4 occupancy on promoter of G_0_-stalled genes (Ncl and Rps24) is higher than that of non-stalled gene Rgs2. Aff4 remains associated with promoters during early reactivation, while stalling is reduced.</u>

Since G_0_-stalled genes were induced <u>by release of pausing</u> in G_1_, we hypothesized that the up-regulation of these genes in Aff4 knockdown cells might prime cells for accelerated cell cycle progression. Indeed, EdU incorporation during cell cycle reentry (6 hr and 12 hr after G_0_ exit) showed that only Aff4 depletion led to accelerated S phase kinetics (~22% EdU^+^ at 12hr compared to ~8% EdU^+^ control cells; <u>re-entry</u> kinetics of Brd4 and Hexim1 siRNA treated cells were unchanged (Figure 5 b, c & Figure 5-figure supplement 3). <u>This accelerated exit from quiescence for Aff4 knockdown cells was observed as early as 2 hours after reactivation as estimated using phosphorylation of pRB S795, a marker of the G0-G1 transition</u> (Zarkowska and Mittnacht, 1997) (Figure 5 d, e). The role of elongation regulators on self-renewal was also assessed (Figure 5-figure supplement 4 a,b): perturbation of Aff4 and Hexim1 led to decreased CFU.

<u>To test if Aff4 acts directly on promoters of G_0_ stalled genes to regulate their timely expression during exit from quiescence, we examined the occupancy of Aff4 on the promoters of G_0_-stalled genes using ChIP</u> (Figure 5 f). <u>During quiescence, Aff4 occupancy was indeed located at promoters of G_0_-stalled genes (Ncl and Rps24) but not on a G_0_ expressed genes (Rgs2), which is consistent with a direct role for Aff4. Further, these promoters continue to be bound by Aff4 as cells exit quiescence.</u> Taken together, these results indicate that in quiescence, the SEC component Aff4 <u>unexpectedly</u> acts <u>as</u> repress<u>or</u> o<u>f</u> G_0_-stalled genes, thereby promoting stalling and re<u>strain</u>ing exit from quiescence<u>, in spite of activating chromatin marks</u>.

Collectively, our results demonstrate that in reversibly arrested myoblasts, Pol II stalling contributes to repression of a set of genes that are not thus marked in permanently arrested myotubes, and that primed reactivation of these genes is critical for reversal of quiescence and for the timing of G_1_-S progression (Figure 6).

**Figure 6.**
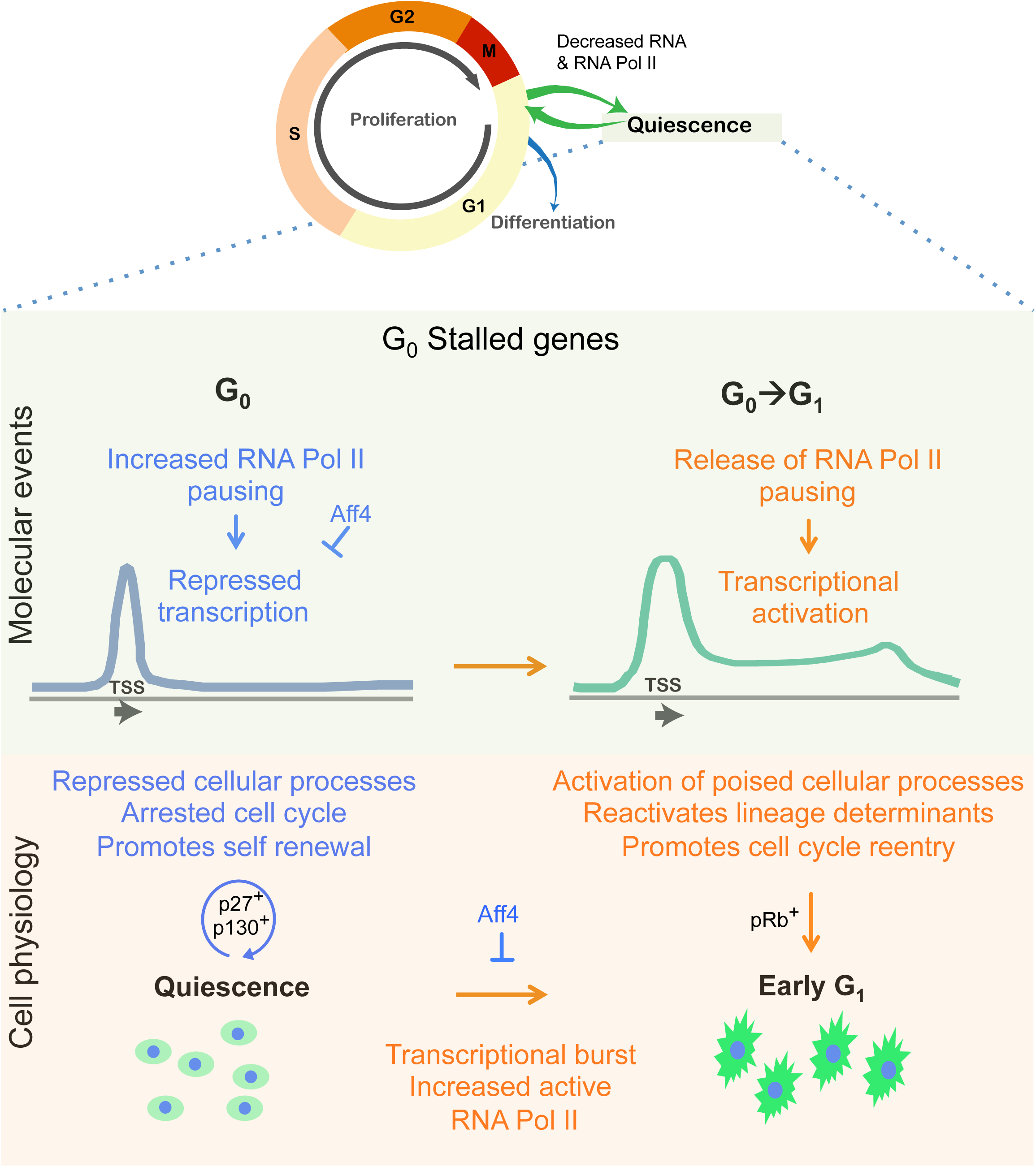
RNA Pol II pausing regulates the quiescent state and G_0_ → G_1_ transition. <u>Proliferating cells can exit G1 into either reversible arrest (quiescence/G_0_) or permanent arrest (differentiation). While both states of mitotic inactivity show enhanced pausing, we identify a distinct regulation of quiescence-specifically stalled genes and its role in cellular fate. Pol II pausing appears to maintain repressed transcription, and thereby contributes to reversibly arrested cellular state and self-renewal. In quiescent cells, Aff4 restrains transcription of stalled genes and blocks cell cycle reentry. During exit from quiescence (G_0_ → G_1_ transition), relief of Pol II pausing leads to coordinated transcriptional activation of poised genes and cellular processes, and reconfigures cellular physiology for timely cell cycle progression.</u>

## Discussion

In this study, we highlight the role of Pol II pausing in the maintenance and exit from the quiescent self-renewing state. While cell cycle exit is known to be associated with reduced transcriptional activity, we show here that this is not uniformly true for all mitotically inactive states: reversible arrest, which is typical of adult stem cells, elaborates global transcriptional control mechanisms distinct from terminally arrested differentiated cells. Consequently, Pol II localization and expression levels, as well as the extent of repression of global RNA content and synthesis differ quantitatively between reversibly and irreversibly arrested cells. In particular, genes regulating RNA biogenesis are among the most strongly repressed in quiescent but not differentiated cells. Strikingly, promoter-proximal polymerase <u>paus</u>ing revealed by ChIP-seq is enhanced in G_0_, genes encoding regulators of RNA biogenesis experience prominent Pol II pausing, and release of pausing on these genes is critical for cell cycle re-entry and self-renewal. Finally, we identify the SEC component Aff4 as a key regulator of quiescence-specific Pol II stalled genes<u>, that also unexpectedly restrains cell cycle re-activation</u>. We conclude that Pol II stalling mediates transcriptional repression that controls quiescence, and that G_0_-stalled genes contribute to competence for G_1_ entry and self-renewal (Figure 6). Given the similarities of gene expression between myoblasts in culture and satellite cells *ex vivo*, we suggest that similar regulatory mechanisms may also function in muscle stem cells.

### Regulation of the RNA Pol II transcriptional machinery distinguishes reversible and irreversible arrest

Reduced transcriptional output is typical of cells whose proliferative activity has slowed, and has been documented in many systems (Bertoli et al., 2013) including quiescent adult stem cells (Freter et al., 2010). Here, we explored the differences in Pol II activity as myoblasts withdrew into alternate arrested states −G_0_ or differentiation- and found several notable differences. First, while the expression of total Pol II was strongly inhibited in both MT and in G_0_, the cytoplasmic localization of Pol II was increased in quiescent cells rather than predominantly nuclear as in MT. Second, the levels of active phospho-Pol II (both Ser5p indicative of initiated Pol II and Ser2p indicative of elongating Pol II) were lower in G_0_ than in MT, suggesting quiescence-dependent regulation at the level of signaling to the CTD kinases/phosphatases. Finally, consistent with the severely depressed total RNA levels in G_0_ compared to either MB or MT, the extent of active RNA synthesis (EU incorporation) was much lower in G_0_ than in MB, but rapidly reversed in response to cell cycle reactivation. Importantly, active RNA synthesis differed in muscle cells *ex vivo*- analysis of freshly isolated skeletal myofibers showed enhanced EU incorporation in activated SC nuclei compared to the differentiated myonuclei. Together, these findings suggest that not only are quiescent cells in culture marked by specific mechanisms that control RNA biogenesis, but also that analysis of RNA synthesis may be useful in distinguishing global cellular states *in vivo*.

### Revisiting the quiescent transcriptome using RNA-seq reveals major differences in RNA biogenesis compared to permanent arrest

Several studies have reported global gene expression profiles of quiescent cells and revealed a core quiescence signature as well as cell type-specific programs (Fukada et al., 2007; Liu et al., 2013; Hausburg et al., 2015). All studies along these lines so far have been comparative analysis using microarrays, where the experimental design forces the use of cRNA derived from equal total RNA. However, our study clearly shows that the altered RNA metabolism in G_0_ myoblasts leads not only to lower RNA synthesis, but also strongly reduced total RNA content. The marked reduction in total RNA content led us to re-evaluate the quiescent transcriptome using RNA-seq: by normalizing transcript levels to cell number and not equal RNA, we reveal a revised picture of the quiescent transcriptome. A major new insight offered by this analysis is the strong repression of a much larger number of genes than previously appreciated, bringing the repression of RNA biogenesis into sharp focus. Interestingly, these pathways are not repressed in differentiated cells, consistent with the observations that Pol II regulation distinguishes two mitotically inactive states, leading to the elaboration of distinct global controls.

The transcriptome studies carried out here also show a distinct quiescence signature in the repertoire of genes up-reglated in G_0_. This observation contrasts with studies of c-MYC overexpression or activation of naïve B-cells (Lovén et al., 2012; Kouzine et al., 2013), where the basal transcription of a set of genes seen at low levels in the resting phase is induced upon overexpression of c-MYC or B-cell activation: i.e. the same set is expressed to a higher extent, thereby increasing the RNA content per cell and causing transcriptional amplification. By contrast, in G_0_ myoblasts, we find a new set of genes is induced in quiescence, and involves distinct mechanisms of global control at the level of Pol II.

### Poising of the replication-dependent histone gene network in reversible arrest

A key new finding from our study concerns the differential transcriptional competence of the histone genes in arrested cells. Although replication-dependent histone mRNA expression is tightly S-phase linked, histone genes are transcribed constitutively, while mRNA processing and degradation is regulated by Slbp (Kaygun and Marzluff, 2005). We now show that while Slbp expression is repressed in both arrested states, Pol II is retained on histone loci only in G_0_ and not MT. Transcriptional control of Slbp has not been previously reported, and Pol II pausing on this regulator might be the pivot for priming the entire histone gene family. Since the replication-dependent histone loci while stalled and repressed, remain primed during quiescence, and stalling of Pol II on the Slbp gene is specific to G_0_, our finding provides a new node for reversible control of replication in G_0_ that is silenced in MT.

Further, the expression of histone variants implicated in marking transcriptionally active or poised regions in the genome (H3f3a and H3f3b (Tang et al., 2013)) is unchanged in quiescence but repressed in MT. These variants (Yuen and Knoepfler, 2013) are translated from polyA^**+**^ transcripts and are required for normal development and growth control (Tang et al., 2013). It is tempting to speculate that expression of specific replication-independent histones is critical for quiescence, and that the condensed chromatin structure (Evertts et al., 2013) and reduced nuclear size peculiar to G_0_ cells (Figure 1e, 4c) may require input of specific histone isoforms.

### A new quiescence-specific regulatory node: increased global Pol II <u>pausing</u>

Accumulating evidence points to quiescence as a poised state where diverse regulatory mechanism at the level of chromatin (Liu 2013; Cheedipudi, 2015), mRNA mobilization (Crist, 2013), and signaling (Rodgers, 2014) work to prime G_0_ cells for cell cycle reactivation. Extending this concept, our study addresses the hypothesis that genes important for cycling cells are kept poised by Pol II pausing, permissive for future expression. Indeed, we identify a distinct class of genes that are regulated by pausing, and allow quiescent cells to boost the activity of the core cellular machinery rapidly after activation. <u>It is quite likely that Pol II pausing in G0 may enable the retention of open chromatin around key (inactive) promoters during quiescence, permitting their activation during cell cycle reentry.</u> Genes exhibiting quiescence-specific stalling are also transcriptionally repressed specifically in G_0_, suggesting a functional relationship between promoter-proximal pausing and low transcriptional output in this state. As this repressed gene set includes RNA metabolism and regulatory proteins, high in the hierarchy of cellular control, Pol II mediated repression of these genes would lead to a reduction in overall cellular throughput, contributing to quiescence. Further, ontologies that include Ca^+^ channels and neuro-muscular signaling were found to be specific for myotubes, suggesting that Pol II stalling in differentiated cells may control responses to neuronal stimulation. Overall, our finding of state-specific <u>paused</u> networks in myoblasts supports the emerging idea of Pol II elongation control as a major regulatory step in eukaryotic gene expression.

### Quiescence-repressed genes are poised for activation in G_1_

An important hallmark of quiescent cells concerns their extended kinetics of S-phase entry compared to that of a continually cycling population. This additional phase - the G_0_-G_1_ transition- has long been appreciated to integrate extracellular signaling (Pledger et al., 1978), but the regulation of its duration is still incompletely understood (Coller, 2007). Our study suggests that Pol II pausing and its reversal may play a role in determining these kinetics. Unlike genes such as Rgs2 and Id2 that are up-regulated in quiescent cells, 95% of G_0_-stalled genes, are repressed. The activation of some G_0_-stalled genes preconfigures changes in cellular processes that are essential for cell cycle reentry. Transcriptional activation of this gene set is accompanied by loss of Pol II stalling, increased active phosphorylation marks (Ser2-p and Ser5-p), and a rapid burst in active RNA synthesis very early in G_1_. The timing of these events suggests that the increase in active Pol II marks is linked to the exit from quiescence. Taken together, <u>we suggest that</u> release of polymerase stalling and the burst of active transcriptional output during reversal of quiescence promotes cell cycle entry and fine-tunes the transcriptional cascade required for de-repressing a variety of cellular processes essential for progression from G_0_ to G_1_ to S.

### Repression of G_0_-stalled genes contributes to blocked proliferation and sustained cell potency

Our studies support the hypothesis that transcriptional repression of G_0_-stalled genes contributes to quiescence entry since direct knockdown of these genes in proliferating cells leads to arrest and compromised self-renewal. Perturbation of ribosomal biogenesis genes (Ncl and Rps24) and the Pol II recycling phosphatase (Ctdp1) led to cell cycle arrest of cycling myoblasts, consistent with the findings that these genes are anti-tumorgenic in other systems (Ugrinova et al., 2007; Badhai et al., 2009; Zhong et al., 2016). Interestingly, Ctdp1/FCP1, which controls the restoration of active Pol II during the transcription cycle, was earlier identified in a genetic screen for quiescence-regulatory genes in yeast (Sajiki et al., 2009). Further, in mammalian cells, Ctdp1 has been implicated in controlling global levels of polyA transcripts as well as in rapid induction of heat shock genes (Kobor et al., 1999; Cho et al., 2001; Mandal et al., 2002; Fuda et al., 2012), both functions ascribed to <u>regulatory functions of</u> Pol II <u>mediated transcription</u>. Thus, stalling <u>control</u> of the Ctdp1 gene itself and its reduced expression in G_0_, may suggest a feed-forward mechanism by which RNA Pol II activity is further depressed in G_0_.

The ribosomal machinery genes, Ncl and Rps24 are involved in the very first processing step that generates pre-rRNA (Ginisty et al., 1998; Choesmel et al., 2008). Rps24 mutations result in Diamond-Blackfan Anemia characterized by ribosome biogenesis defects in HSCs and erythroid progenitors (Choesmel et al., 2008; Song et al., 2014). Stalling of genes involved in ribosomal RNA processing in G_0_ suggests an upstream blockade on ribosome biogenesis, and by extension, the translational machinery. Thus, G_0_-specific <u>Pol II paus</u>ing participates in keeping the major biosynthetic pathways in check to maintain the quiescent state.

### The SEC component Aff4 regulates G_0_-stalled genes to control the timing of S-phase entry

Quiescence-specific regulation of Pol II has not been previously reported. Regulators of stalling such as DSIF and NELF are widely involved and likely to participate in all states. Therefore, we investigated specific components of the P-TEFb regulatory system complex as good candidates for the release of stalling of specific target genes during G_1_ (Lu et al., 2016). Release of stalled polymerase is brought about by coordinated regulation of P-TEFb mediated by SEC, Brd4, Hexim1 (Lu et al., 2016; Puri et al., 2015). Hexim1 haplo-deficiency in mice increases satellite cell activity and muscle regeneration, highlighting a role in self-renewal (Galatioto et al., 2010; Hong et al., 2012). While Hexim1 knockdown did not affect the timing of cell cycle re-entry, an effect in quiescence cannot be ruled out and the repression of some G_0_-stalled genes warrants further investigation. The pause-release regulator that had the most interesting phenotype in our study was Aff4. As a SEC component, Aff4 is known to display target gene specificity (Luo et al., 2012), promoting release of paused polymerase on heat shock genes and MYC (Kühl and Rensing, 2000; Luo et al., 2012; Schnerch et al., 2012). By contrast, we identified a <u>new</u> repressive role for Aff4 in G_0_, such that knockdown led to enhanced expression of G_0_-stalled genes, and accelerated exit from quiescence. Therefore, it is tempting to speculate that Aff4 mediates differential regulation (activation and repression) of stalling on different target genes, thereby regulating cell cycle progression or arrest. Moreover, the accelerated S-phase entry in knockdown quiescent cells indicates that Aff4 guards against premature activation of quiescent cells, thereby regulating the timing of cell state transitions.

In conclusion, by combining a genome-wide analysis of Pol II occupancy with its functional transcriptional output in distinct mitotically arrested states, we uncover a repressive role for Pol II pausing in the control of RNA metabolic genes that is central to the entry into quiescence. We also define a new regulatory node where the SEC component Aff4 acts to restrain expression of these genes in G_0_ to set the timing of their activation during G_1_ entry and affects S-phase progression. Overall, this study extends the role of Pol II pausing in regulating not only developmental lineage transitions (Scheidegger and Nechaev, 2016), but also distinct cell cycle states and in particular, sheds light on the type of arrest typical of adult stem cells.

## Acknowledgements

We thank Karen Adelman (NIH), Aseem Ansari (U. of Wisconsin), Boudewijn Burgering (Utrecht), Dasaradhi Palakodeti (InStem) and Aswin Seshasayee (NCBS) for helpful discussions, Michal Mokry (Hubrecht Inst) for input to customize the ChIP-seq analysis pipeline, Dawn Cornelison (U. of Missouri) and Judy Anderson (U. of Manitoba) for teaching us myofibre methods, Shahragim Tajbakhsh (Institut Pasteur) for providing Tg:Pax7-nGFP mouse, and Ramkumar Sambasivan (InStem), Suchitra Gopinath (THSTI), Purnima Bhargava (CCMB), Surabhi Srivastava (CCMB), and Sanjeev Galande (IISER-Pune) for critical reading of the manuscript. We gratefully acknowledge facilities at the Bangalore Life Sciences Cluster (NGGS facility for sequencing, Animal Facility and CIFF for imaging and flow cytometry at C-CAMP, inStem,/NCBS Bengaluru), and at Advanced Imaging Facility and the Animal Facility at CCMB, Hyderabad. We acknowledge graduate fellowships from Govt. of India Council for Scientific and Industrial Research (CSIR) (H.G., D.S.), DBT (N.V.) and DAE-NCBS (A.A.). H.G. was also supported by a short-term fellowship from CSIR-CCMB. J.D. is supported by core funds to CCMB from CSIR, an Indo-Australia collaborative grant and an Indo-Danish collaborative grant from DBT. Animal work was partially supported by the National Mouse Research Resource (NaMoR) grant (BT/PR5981/MED/31/181/2012;2013-2016) from the DBT to NCBS. We gratefully acknowledge an InStem collaborative science chair fellowship to Boudewijn Burgering and support from Utrecht for exchange visits between JD and BB labs.

## Author contributions

HG and JD designed research, interpreted data and wrote the paper; HG performed research, acquired and analysed data, and wrote custom scripts for NGS analysis and quantitative image analysis; DS, NV and AA performed research. The authors declare no conflict of interest.

## Materials and Methods

***Cell Culture.*** C2C12 myoblasts (MB) were obtained from H. Blau (Stanford University) and a subclone A2 (Sachidanandan et al., 2002) were used in all experiments. Myoblasts were maintained in growth medium (GM; DMEM supplemented with 20% FBS and antibiotics) and passaged at 70-80% confluency. ***Differentiation***. Myotube (MT) differentiation was induced in proliferating cultures at 80% confluence after washing with PBS and incubation in differentiation medium (DM: DMEM with 2% horse serum), replaced daily for 5 days. Multinucleated myotubes appear after 24 h in DM. ***Quiescence induction:*** G_0_ synchronization by suspension culture of myoblasts was as described (Sachidanandan et al., 2002). Briefly, subconfluent proliferating MB cultures were trypsinized and cultured as a single-cell suspension at a density of 10^5^ cells/mL in semisolid media (DMEM containing 1.3% methyl cellulose, 20% FBS, 10 mM HEPES, and antibiotics). After 48 h, when ~98% of cells have entered G_0_, arrested cells were harvested by dilution of methyl cellulose media with PBS and centrifugation. G_0_ cells were reactivated into the cell cycle by replating at a sub-confluent density in GM and harvested at defined times (30‵ - 24hr) after activation. Upon replating, G_0_ cells undergo a G_0_–G_1_ transition to enter G_1_ ~6 h after reattachment; S-phase peaks at 20-24 h.

### Muscle fiber isolation and culture

Animal work was conducted in the NCBS/inStem Animal Care and Resource Center and in the CCMB Animal Facility. All procedures were approved by the inStem and CCMB Institutional Animal Ethics Committees following norms specified by the Committee for the Purpose of Control and Supervision of Experiments on Animals, Govt. of India.

Single muscle fibers were isolated and cultured using the Anderson lab method with some modifications (Shefer and Yablonka-Reuveni, 2005; Siegel et al., 2009). Briefly, the EDL muscle was dissected from 8-12 week Tg:Pax7-nGFP mice (Sambasivan et al., 2009). Isolated muscles were digested with Type I collagenase (Worthington) in DMEM at 37ᴼC, till single fibers dissociated. All dissociated fibers were transferred into fresh DMEM medium and triturated to release individual fibers using fire polished pasteur pipettes. Fibers were cleaned by multiple transfers through media washes and then to fiber culture media (DMEM (Gibco), 20% Fetal bovine serum (Gibco), 10% Horse serum (Gibco), 2% chick embryo extract (GENTAUR), 1% penicillin streptomycin (Gibco), 5ng/ml bFGF (SIGMA)).

For quantification of active RNA synthesis, 10 μM EU was added to media. Freshly isolated or cultured fibers were analyzed by immuno-staining. Single fibers were fixed in 4% PFA for 5min, washed 3X with PBS, picked and placed on charged slides (Thermo-Fisher) for immuno-staining.

### Knockdown of target genes using RNAi

C2C12 cells were cultured in growth medium until 80% confluent. The cells were then trypsinized and ~0.3 × 10^6^ cells were plated on 100 mm tissue culture dishes. Approximately 12-14hr post plating, the cells were transfected with siRNA (400 picomoles of siRNA with 40 μl lipid) using Lipofectamine RNAiMAX (Invitrogen) as per manufacturer's instructions. The cells were incubated with the RNA-Lipid complex for at least 18-24hr following which they were tested to evaluate the extent of knockdown by qRTPCR and western blot and then used for further experimental analysis. Typically, siRNA-mediated knockdown was in the range of 50-90% at RNA level. siRNA used in this study are detailed in Table S3.

### Immunofluorescence and microscopy

Cells plated on cover slips or harvested from suspension cultures were washed with PBS, fixed in 2% paraformaldehyde at room temperature, and permeabilized in PBS + 0.2% TritonX100. Primary antibodies (Table S4) were diluted in blocking buffer (1XPBS, 10% HS, 0.2 % TritonX100). Secondary antibody controls were negative, no cross reactivity of secondary reagents was detected. For total RNA quantification, fixed cells were incubated with 10 μM Syto^®^ RNASelect^TM^ for 30‵. For detection of DNA/RNA synthesis, cells were pulsed for 30‵ with EdU/EU, fixed with 2% PFA for 15’ at RT and labeling was detected using Click-iT^®^ imaging kit (Invitrogen) as per manufacturer’s instructions. Samples were mounted in aqueous mounting agent with DAPI. Images were obtained using a Leica TCS SP8 confocal microscope (63X, 1.4 NA). Minimum global changes in brightness or contrast were made, and composites were assembled using Image J. The automation of intensity calculation and counts was carried out using a custom pipeline created on the Cellprofiler platform (Source Code File - Cellprofiler_processing_images, Source Code File - Cellprofiler_processing_pipeline) (Carpenter et al., 2006; Kamentsky et al., 2011).

### Cell cycle analysis using flow cytometry

Adherent cells were trypsinized, washed in PBS and pelleted by centrifugation. Suspension-arrested cells were recovered from methyl-cellulose by dilution with PBS followed by centrifugation as described earlier. Cell pellets were titurated in 0.75 ml of PBS, fixed by drop wise addition into 80% ice-cold ethanol with gentle stirring. Following fixation (cells could be stored up to 7 days at -20^0^C), cells were briefly washed with PBS and re- suspended in PBS at 1 million cells per 500 μl PBS containing 40 μM DRAQ5^TM^ (Cat. No DR50050, Biostatus) and 10 μM Syto^®^ RNASelect^TM^ (cat. No S-32703, Invitrogen). Cell cycle analysis was performed on a FACS Caliber^®^ cytometer (Becton Dickinson), DRAQ5^TM^ (wide range of absorption / emission maxima = 697 nm) and Sytox^®^ Green (absorption / emission maxima = ~490 / 530 nm). 10,000 cells were analyzed in each sample. CelQuest^®^ software was used to acquire <u>For ***cell cycle and EdU co-staining*** -Cells were pulsed with EdU and fixed as described, washed twice in PBS and resuspended in PI staining solution for 30 minutes in dark. For analysis of S-phase cells during cell cycle reentry, EdU pulsed cells were fixed in 4% paraformaldehyde for 10 min, washed and stored in PBS at 4C. For staining, cells were permeabilized and blocked in PBS having 10% FBS and 0.5% TX-100 and labeling was detected using Click-iT^®^ imaging kit (Invitrogen) as per manufacturer’s instructions. Cells were co-stained with PI (50 μg/ml, RNaseA 250 μg/ml). Cell cycle analysis was performed on Gallios cytometer (Beckman Coulter). Kaluza software (ref) was used to acquire and analyze the data.</u>

#### Analysis of p27 induction

<u>A C2C12 line expressing p27mVenus (a fusion protein consisting of mVenus and a defective mutant of p27-CDKI, (p27K(−)(Oki et al., 2014)) was generated by stable transfection. Adherent myoblasts expressing p27-mVenus (excitation/emission= 515/528), +/− knockdown of various G0-stalled genes were trypsinized and fixed in 4% paraformaldehyde for 10 minutes at room temperature. Suspension cultured cells were recovered as described above. Following fixation, cells were analyzed for mVenus expression using a Gallios cytometer (Beckman Coulter). Kaluza or FlowJo software was used to acquire and analyze the data.</u>

### Western blot analysis

Soluble lysates of adherent cultures or suspension cells (2×10^6^) were obtained after harvesting PBS washed cells by centrifugation, resuspended in 500μl of lysis buffer (10mM HEPES, pH 7.9, 1.5mM MgCl2, 10mM KCl, 0.5mM DTT, 1.5mM PMSF and protease inhibitor cocktail (Roche). Protein amount in lysates was estimated using Amido Black staining and quantification of absorbance at 630 nm. Proteins were resolved on 8% Acrylamide gels. and transferred to PVDF membrane. The membrane was incubated with primary antibody overnight at 4°C, then washed in 1X TBS + 0.1% Tween for 10’ each followed by incubation with secondary antibody conjugated with HRP (Horse radish peroxidase) for 1 hr. After a brief wash with TBS-T for 10’, ECL western blotting detection reagent for HRP was used for chemiluminescent detection using gel documentation system (Vilber Lourmat). Details of antibodies and dilutions used in this study are given in Table S4.

### Isolation of RNA from cultured cells and RNA sequencing library preparation

Accurate determination of cell number in different states: Attached cultures or suspension cells were counted 3 times (at 2 or more dilutions) using trypan blue exclusion to generate a live cell count using a Countess^®^ Automated Cell Counter (Invitrogen). For MT samples, being multinucleate cells, accurate counts were not possible since cells with varying number of nuclei were present; therefore nuclei count was used as cell number counts.

Cells were washed twice with cold PBS, residual PBS removed and cells lysed using 1 ml RLT plus buffer containing β-mercaptoethanol (RNeasy Plus Mini Kit, Qiagen). The lysate was vortexed vigorously to shear genomic DNA and then stored at –70^0^C until further processing. RNA was isolated from cultured cells using RNeasy Plus Mini Kit (catalog no 74134, Qiagen) as per manufacturers’ protocol, using RNeasy MinElute spin column and eluted twice in 30 μl water, quantified by NanoDrop ND-1000UV-Vis spectrophotometer (NanoDrop Technologies, Wilmington, DE) and checked by gel electrophoresis and q-RT PCR for marker transcripts. For RNA-seq, RNA quantification and quality check was done using Agilent RNA 6000 Pico Kit on Agilent 2100 Bioanalyzer system.

RNA sequencing libraries were prepared from duplicate MB, G_0_ and MT samples and by quantifying the exogenously added “spike-in mix”, we determined that each of the 6 libraries were referenced to RNA content per million cells, correcting for sampling biases generated during sequencing. 3 μl per million cells of ERCC spike-in RNA (1:10 dilution) was added to each sample. Since the RNA-seq experiment was carried out in duplicates, ERCC Mixes were added to distinguish each replicate sets (MB, MT, G0) i.e Mix1 added for MB_1, MT_1, G0_1 (similarly for replicate2 and mix2). The spiked total cellular RNA was DNase treated to remove trace amounts of DNA (DNA-freeTM kit, AmbionTM Cat. No AM1906) and 4 μg of DNase-treated spiked RNA was further processed to remove ribosomal RNA (RiboMinusTM Eukaryotic kit V2, AmbionTM cat no A15020). The Ribominus RNA was quality checked using BioAnalyser. Finally, the purified mRNA was used for cDNA synthesis and library preparation using NEBNext^®^ UltraTM Directional RNA Library Prep Kit for Illumina^®^ (New England Biolabs^®^ cat. No. E7420). The library thus prepared was quality checked on BioAnalyser and verified by q-PCR for marker transcripts prior to carrying out paired-end sequencing on Illumina HiSeq1000.

### Chromatin immunoprecipitation (ChIP) and ChIP-seq library preparation

***Cell harvest and sonication:*** Wild type C2C12 (8×10^6^) were detached from the substratum by cell dissociation buffer (Sigma) collected in 15 cc tubes and washed with PBS. Cell pellets were re-suspended in growth medium with final concentration of 1% HCHO (Sigma). Cells were incubated for 10’ at 37°C followed by two washes with cold PBS and finally re-suspended in 1.6 ml of SDS lysis buffer (1% SDS, 10 mM EDTA and 50 mM Tris HCl pH 8.1). Lysates were incubated on ice for 20’ and spun at 12,000 rpm for 10’ at 4°C. 200 μl samples were subjected to sonication for 60 sec on/off for 16 cycles for G_0_ and MB samples and 18 for MT (Bioruptor ^®^ sonicator). The aliquots of sample were stored at -80^0^C. ***Immuno-precipitation:*** Chromatin Immunoprecipitation (ChIP) Assay Kit (milliore catalog no 17-295) was used to carry out immunopreicitation as per manufactures protocol. Briefly 200 <u>μl</u> of sonicated lysates were diluted up to 2 ml by adding ChIP dilution buffer and pre-cleared with Dynabeads^TM^ A beads (agarose beads from the kit were not used). Cleared supernatants were recovered and incubated with 7.5 μg antibody (anti RNA Pol II; N20clone/ rabbit IgG) for 16hr at 4°C on a rotary shaker. Dynabeads A/G were added to collect immune complexes. Washes (1ml for each) in a series of buffers were performed as follows: low salt immune complex wash buffer, High salt immune complex wash buffer, LiCl wash buffer and twice in TE (10 mM Tris HCl and 1 mM EDTA). All washes were done at 4°C on a rotary shaker and collected by centrifugation at 7000 rpm for 5’ each. Finally, the beads were re-suspended in 250 μl freshly prepared elution buffer (1% SDS, 0.1 M NaHCO3) and incubated at RT for 15’ each. The elution process was repeated twice and 20 μl of 5 M NaCl was added to 500μl of combined eluate for reverse cross-linking at 65°C for 16hr. Reverse cross-linked eluates were subjected to proteinase K digestion at 45°C for 1 hr. Qiagen nucleotide removal kit was used to purify DNA. Purified DNA was subjected to real-time PCR with primers as shown (Table S5). Each DNA sample was analyzed in triplicate, in at least three independent experiments.

***ChIP-sequencing library preparation*:** 20 ng of pull-down/input DNA was used to carry out sequencing library preparation using NEBNext^®^ DNA Library Prep kit for Illumina^®^ (New England Biolabs^®^, cat. No. E6000) along with NEBNext^®^ Multiplex oligos for Illumina^®^ (Index Primers set 1) (New England Biolabs^®^, cat. No. E7335), as per manufacturer’s protocol. The library thus prepared was quality checked on BioAnalyser and qPCR prior to carrying out paired-end sequencing on Illumina HiSeq1000.

### Sequencing data analysis

The ChIP-seq and RNA-seq raw data from this publication have been submitted to the GEO and can be accessed at link (http://www.ncbi.nlm.nih.gov/bioproject/343309)

#### RNA-seq

Sequenced libraries were aligned using Tophat to *Mus musculus* reference genome (mm9 build with 92 spike-in mRNA sequences added as pseudo-chromosomes). Reference annotation Version M1 (gtf file, from July 2011 freeze, NCBIM37 - Ensembl 65 from GENCODE consortium) along with ERCC transcripts was used.

##### For equal RNA normalization

Cufflinks was used for transcript abundance and differential expression analysis (Cuffdiff) (Trapnell et al., 2012; Goff L et al., 2013). Cummerbund was used for sub-selection of differentially expressed gene list (Figure 1 - source data 1) and for representation of expression using heatmap (Figure 1-figure supplement 2).

##### For Equal cell number normalization

HTcount packages (Anders et al., 2015) was used to quantify reads per gene for each of the samples using reference annotation file. This was further normalized using DEseq2 package using custom R-scripts generated specifically for this normalization (Source Code File - RNA_Seq_DEseq_processing). The sizeFactor is estimated for the Group B counts of ERCC spike-in RNA mix since the group B genes must not vary across all samples of both mixes. The calculated sizeFactor for each sample using Group B as reference gene set is further used to normalize other genes in the respective library including those in the other ERCC spike in mix groups (Groups A, C, D). This approach ensures that the RNA-seq libraries are scaled to reference spike-in controls, and not to the sequencing depth of the library. For each of the sub groups A, C and D, we found that the mix sets 1 and 2, cluster as distinct groups and the levels of intensity within each of the mix sets is comparable. Also, the fold differences between the mix sets 1 and 2 were observed to be as expected as per the manufacturer’s descriptions for all groups. Differentially expressed gene was then identified for each of pair wise comparisons (Figure 1 - source data 1). The heatmap of ERCC spike in mixes groupB and groupA transcripts after normalization step (Figure 1 -figure supplement 2).

***Cross-comparison between normalization methods:*** Overlap of gene identity for differentially expressed genes identified from both normalization methods (Equal RNA and equal cell number) (>2 log2 fold change and p-value<0.05) is represented using size adjusted Venn diagrams separately for both up-regulated and down- regulated genes (Figure 1h).

##### Correlation between replicates and samples

Dissimilarity between the sample and replicates were computed using the normalized gene count matrix to get the sample to sample distance using Euclidean Distance Method using DEseq packages. The distance matrix is further is clustered hierarchically and heatmap is plotted (Figure 1-figure supplement 1).

##### Gene ontology studies

Panther tools (Thomas et al., 2003) and gProfiler (g:Cocoa) (Reimand et al., 2011) was used for over representation analysis of ontologies for single and multiple gene lists respectively. For GSEA, cell number normalized expression values (for MB and G_0_) used as expression dataset and gene sets (QSCs (437) and ASC36h (355) ASC60h (1588)) as identified from earlier published dataset (Liu et al., 2013) and enrichment analysis carried out as described in manual (http://software.broadinstitute.org/gsea/doc/GSEAUserGuideFrame.html).

### ChIP-seq

After quality assessment, the raw ChIP-seq reads were aligned using paired option using bowtie2 to mouse genome (NCBIM37/mm9).

#### RNA Pol II enrichment at TSS

Only non-overlapping gene >2500 bases were used for analysis (except for histone genes panels). Using these criteria 15170 genes were used for analysis. The read density counts around TSS (+/- 300) was enumerated for all samples. Independently, for each cell state (MB, MT, G_0_), the TSS regions were sub group into greater than 90 percentile and 70-90 percentiles based on read density score sorted lowest to highest value. Thus for each cell state (both input and IP) the average distribution of read densities were plotted for the each subgroups (Figure 2 f).

#### Stalling index

Only non-overlapping gene >1500 bases were used for analysis (except for histone genes panels). Using these criteria 16095 genes were used for analysis. To calculate RNA Pol II stalling index, the number of ChIP-seq reads at promoter region (-500 to +300 bp from TSS) and gene body (+300 to +2500 for genes longer than 2.kb and +300 to +1500 for genes between 1.5kb to 2.5kb) of each gene was enumerated using Seqmonk. Stalling index for all genes were plotted in Figure 2g. The statistical significance of RNA Pol II stalling index was assess using the Benjamini-Hochberg method corrected Fisher's exact tests and to control the false discovery cut off was (p-value<0.005) and Stalling index greater than 2.5 in G_0_ IP samples were used in Figure 3a.

##### Statistical analysis

R-packages were used to carryout statistical significance tests. Mann-Whitney test and Student T-test were used as mentioned in the respective figure legends. For the box plots the dark horizontal bar represents median value; the upper and lower limits of box represents 75 and 25 percentiles for each box.

---

**Figure 1-figure supplement 1a.**
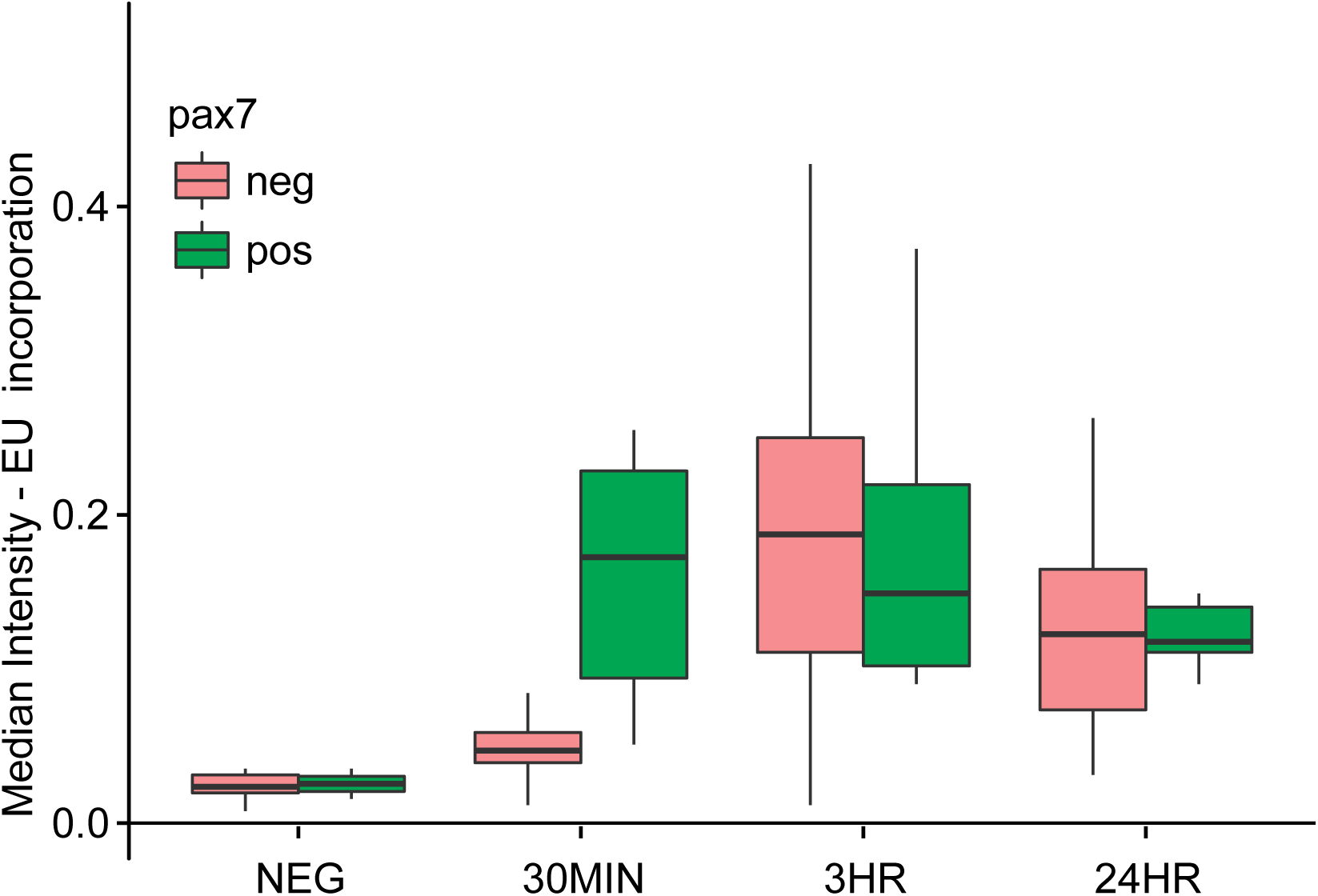
Satellite cells associated with freshly isolated myofiber are activated rapidly during the isolation protocol. Boxplot for median intensity of EU incorporation: Pax7 positive (green) satellite cells & myonuclei (red) for ex vivo cultures at various time points (30min, 3hrs and 24hrs post isolation).

**Figure 1-figure supplement 1b.**
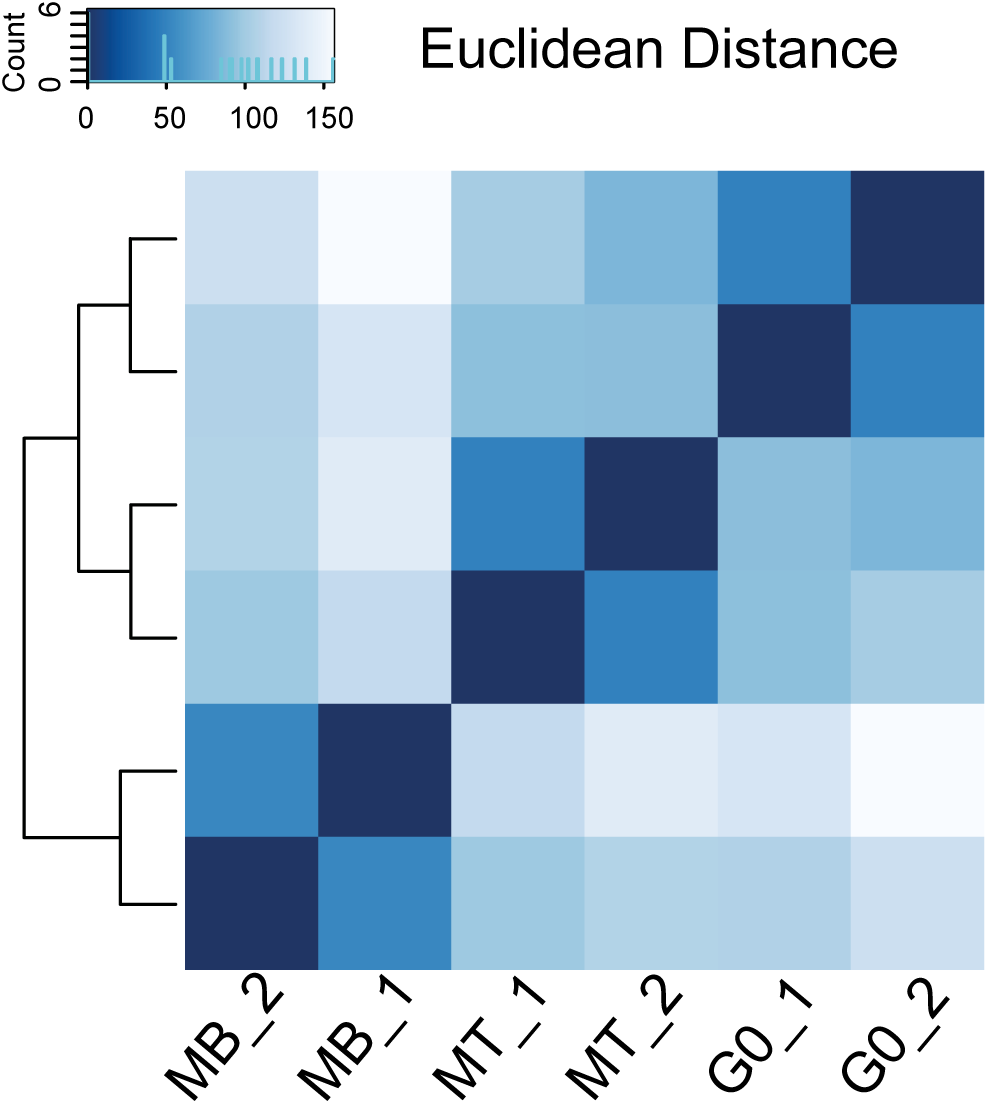
Cell number normalization of the RNA-seq dataset does not affect quantification as the replicates cluster together. In order to confirm that the modified normalization method employed in this study does not affect the similarity between samples and replicates, the Euclidean distances between the samples was calculated from the regularized log transformation (Deseq package). The distance is color coded and shown in the key scale in the heat map. The dendogram shows the distance between samples and their replicates, indicating that the replicates behave similarly, while the three states are indeed distinct from each other.

**Figure 1-figure supplement 2.**
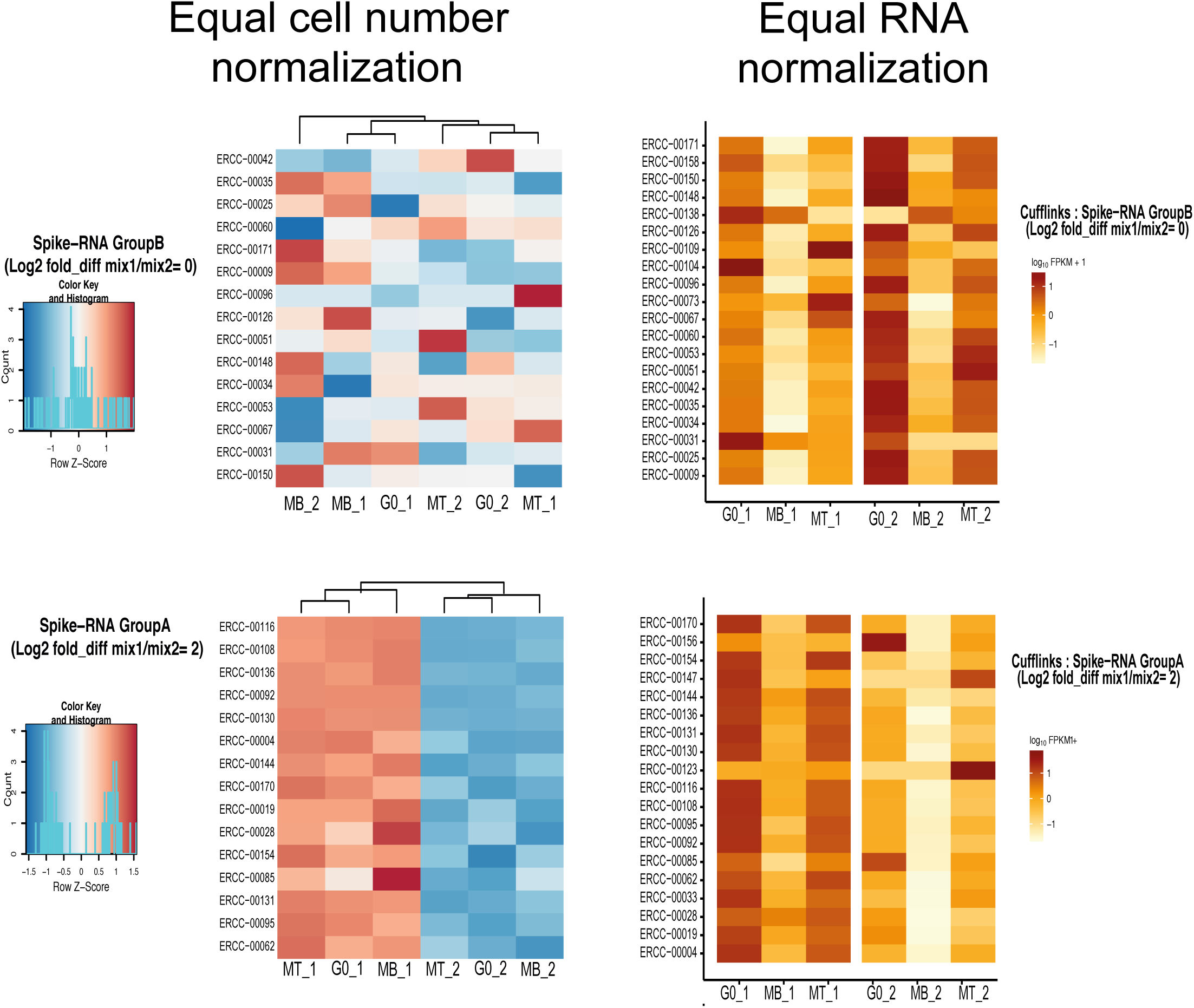
Effect of normalization method on Spike in RNA group B (unchanged between mixes) and group A (mix1: mix2 = 4 fold). Heat map showing the extent of representation of ERCC RNA in each of the MB, G0 and MT libraries. The value for each RNA is scaled within the row and relative intensity is color coded as shown on adjacent scale bar. Right panel = Equal RNA normalization: Extent of representation of group A and B Spike in transcripts in left panel is similar ie G0 > MT >MB for both replicates Left panel = Equal cell number normalization: Group B transcript used as reference for normalization and histogram shows their value is close to zero. Group A transcripts show average difference of 2 (log2 scale) between the replicate set 1 (ERCC mix1) and replicate set 2 (ERCC mix 2).

**Figure 1-figure supplement 3.**
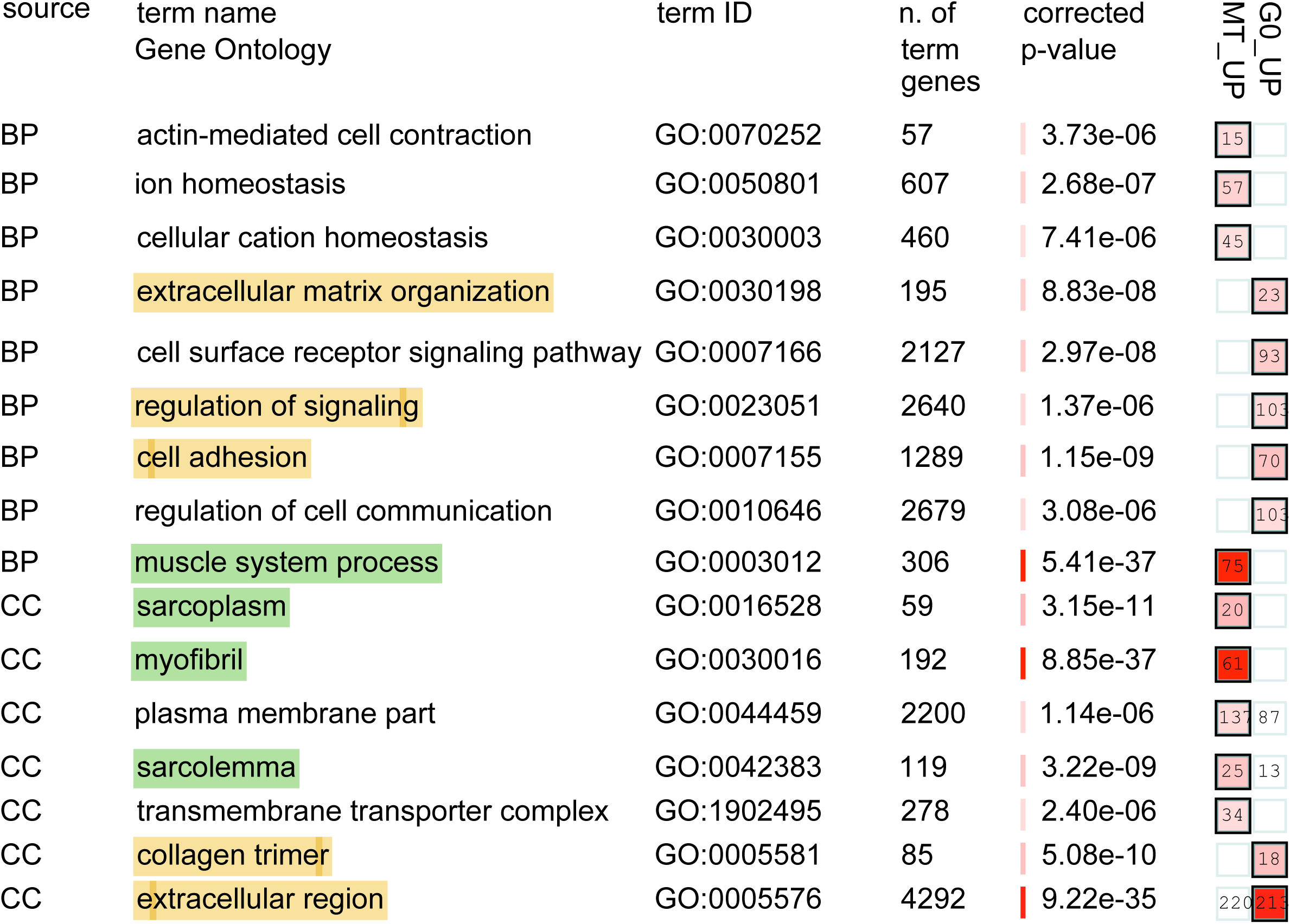
Gene ontology analysis of genes differentially regulated between two mitotically arrested states (up-regulated genes). Biological process comparison for two mitotically arrested states (Up regulated genes) gProfiler analysis carried out for genes up-regulated in G0 vs MB and MT vs MB comparisons. The output is graphically represented here using moderate option for Biological process and cellular component. The terms enriched in each state are listed on the left. The intensity of color in red square on right indicates the significance of each state. The most significant among the 2 gene lists are highlighted with bold boundaries around square and corresponding p value is shown next to it and shaded in red square. Yellow highlight terms are enrich specifically in G0 and green are represented from MT state.

**Figure 1-figure supplement 4.**
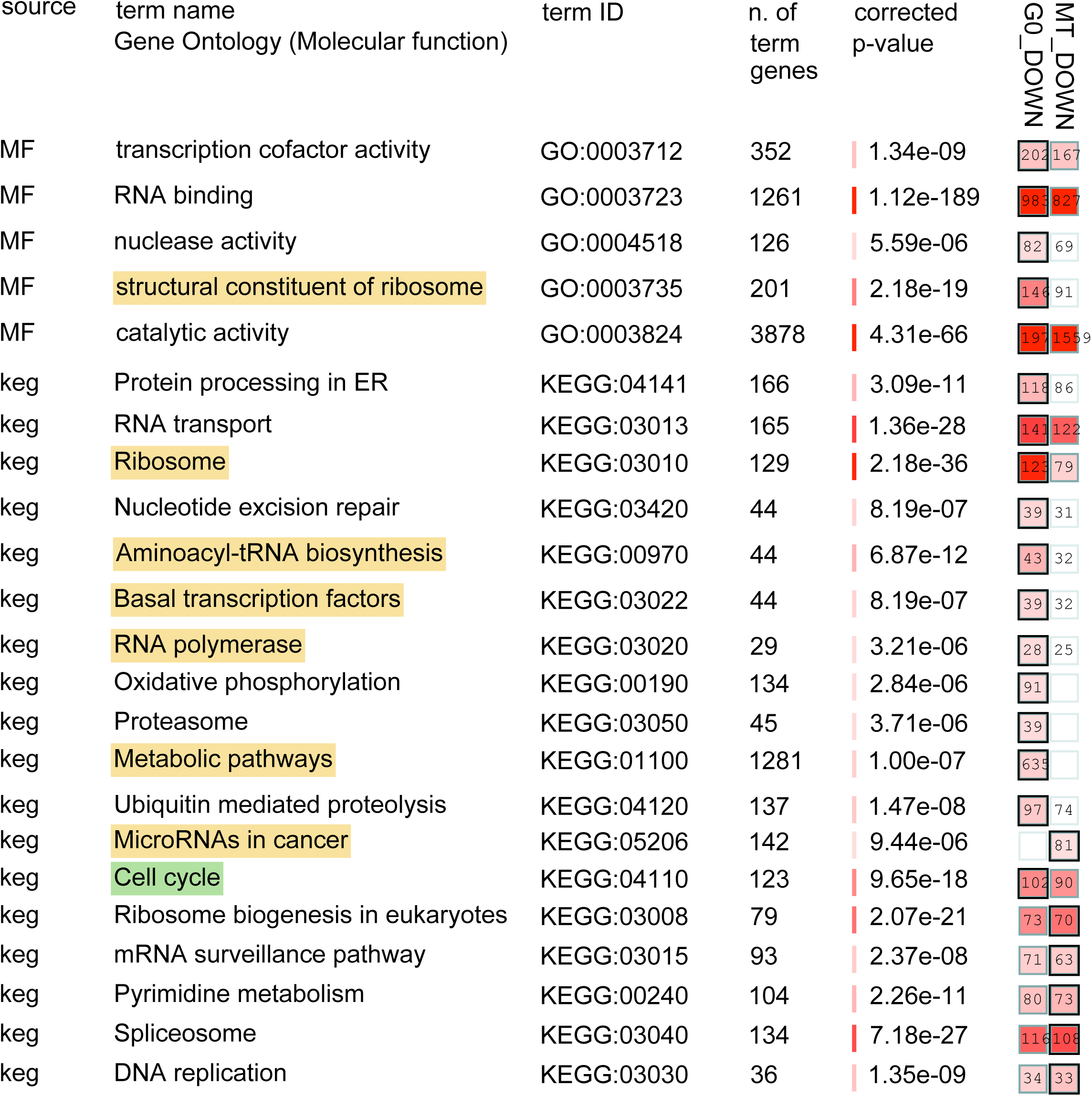
Gene ontology analysis of genes differentially regulated between two mitotically arrested states (Down-regulated genes). Biological process comparison for two mitotically arrested states (Down regulated genes) gProfiler analysis carried out for genes down-regulated in G0 vs MB and MT vs MB comparisons. Yellow highlight terms are enrich specifically in G0 and green are represented from Both G0 and MT states.

**Figure 1-figure supplement 5.**
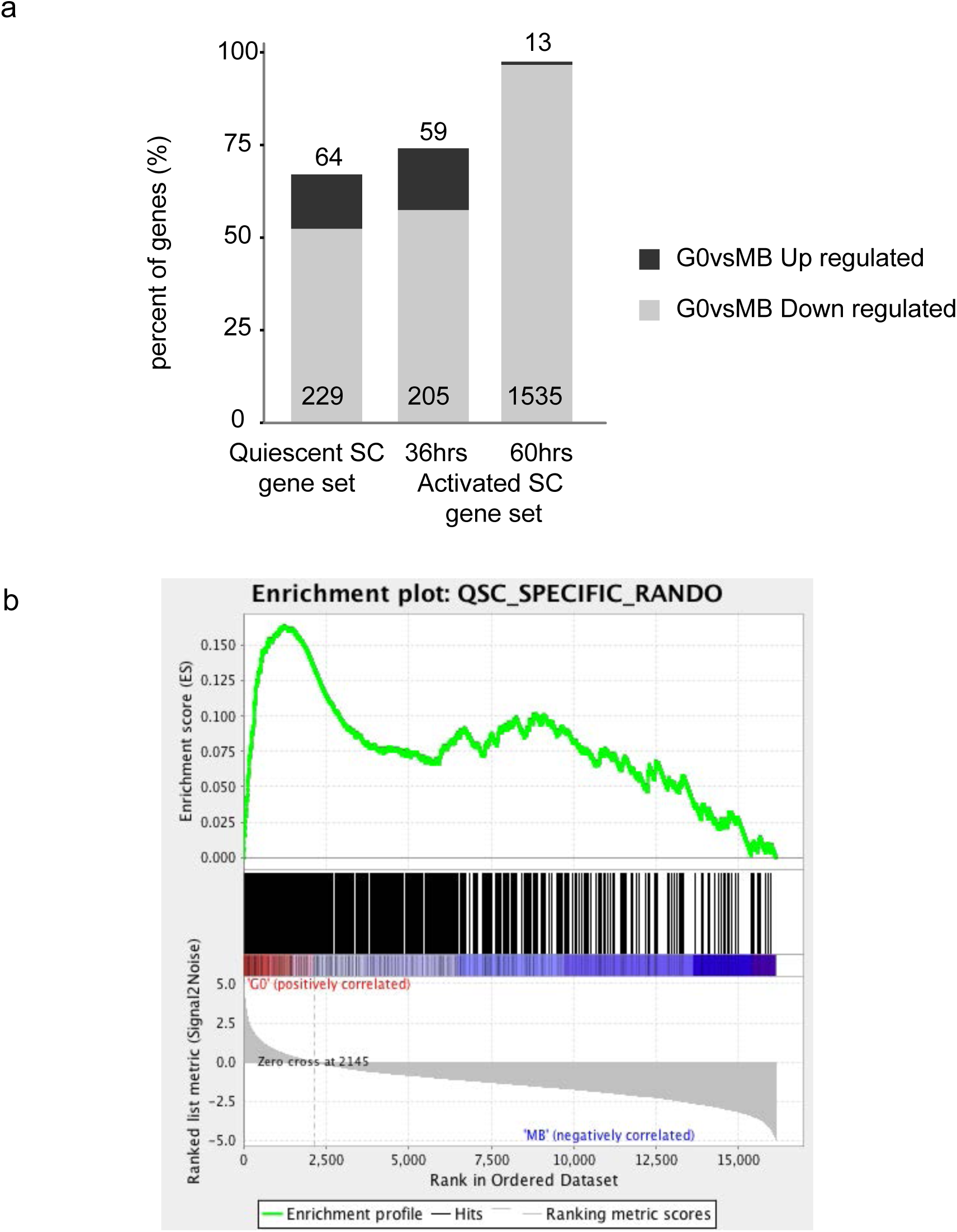
Identifying similarity between G0 in cultured C2C12 myoblasts (this study) and freshly isolated satellite cells. a. Percentage of overlapping genes between differentially expressed genes in G0 vs. MB comparison (Up/Down regulated) for each of gene sets QSCs (437) and ASC36h (355) ASC60h (1588) gene set identified in (Liu et al., 2013). Note that although Quiescent SC has genes higher fraction of genes that are Down regulated in G0 vs MB comparisons suggesting QSC gene list depicts partially activated cell state. ASC 60hrs overlap almost entirely with G0 down regulated genes ie ASC60 hrs is similar to MB. b. Leading edge analysis using Gene set enrichment analysis (GSEA) comparing G0 vs. MB: Geneset used here is 437 gene identified as QSCs specific genes (Liu et al., 2013) and expression dataset generated in our study was used as reference. Note that these genes partition into bimodal distribution one node enriched for both G0 and MB statex

**Figure 1-figure supplement 6.**
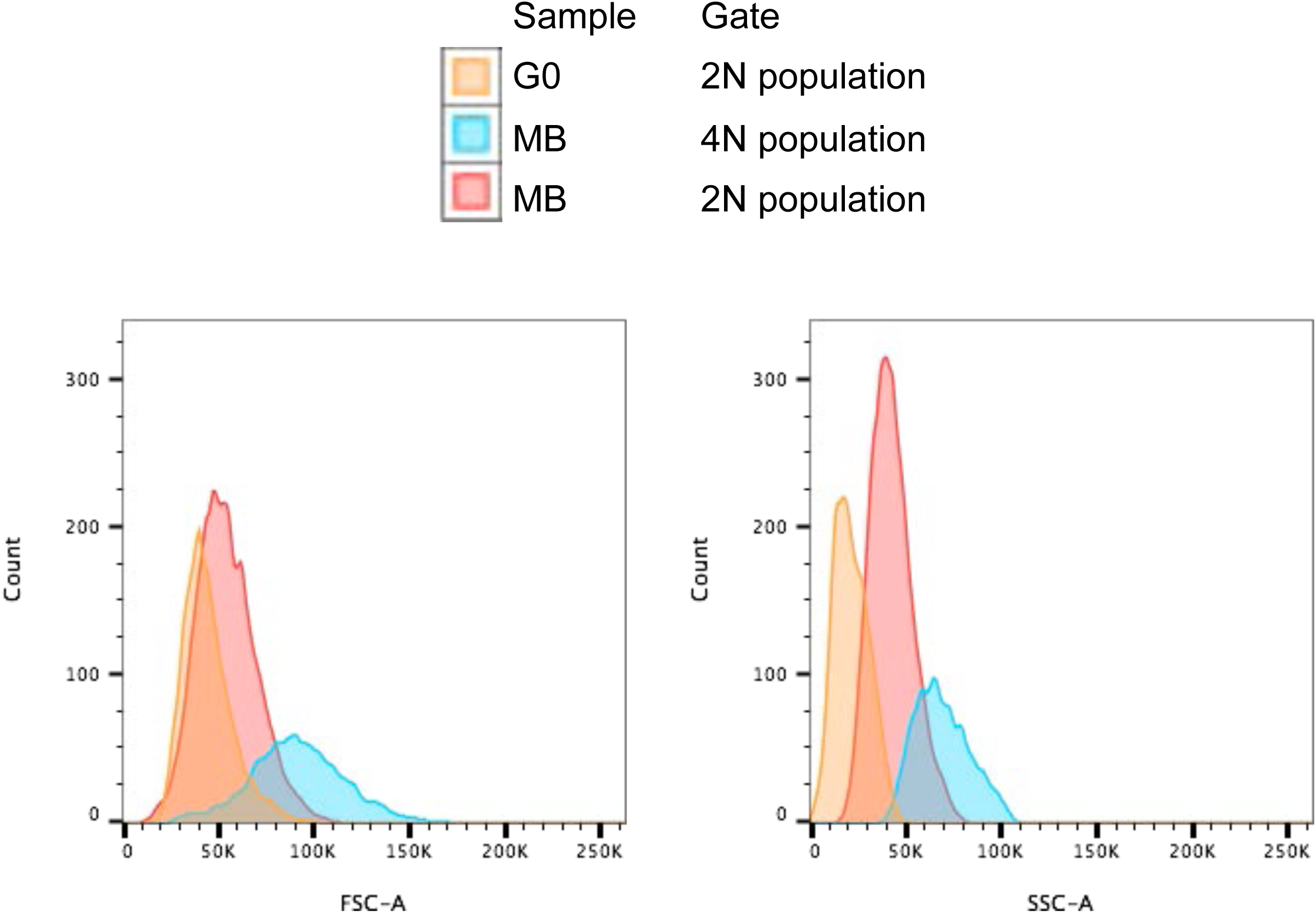
Volume estimation of MB and G0 cell containing 2N and/or4N DNA content. FSC and SSC didstribution for gated cellls is used as indicator for volume. G0 celll are observed to be markedly reduced in volume compared to MB 2N or 4N population.

**Figure 2-figure supplement 1.**
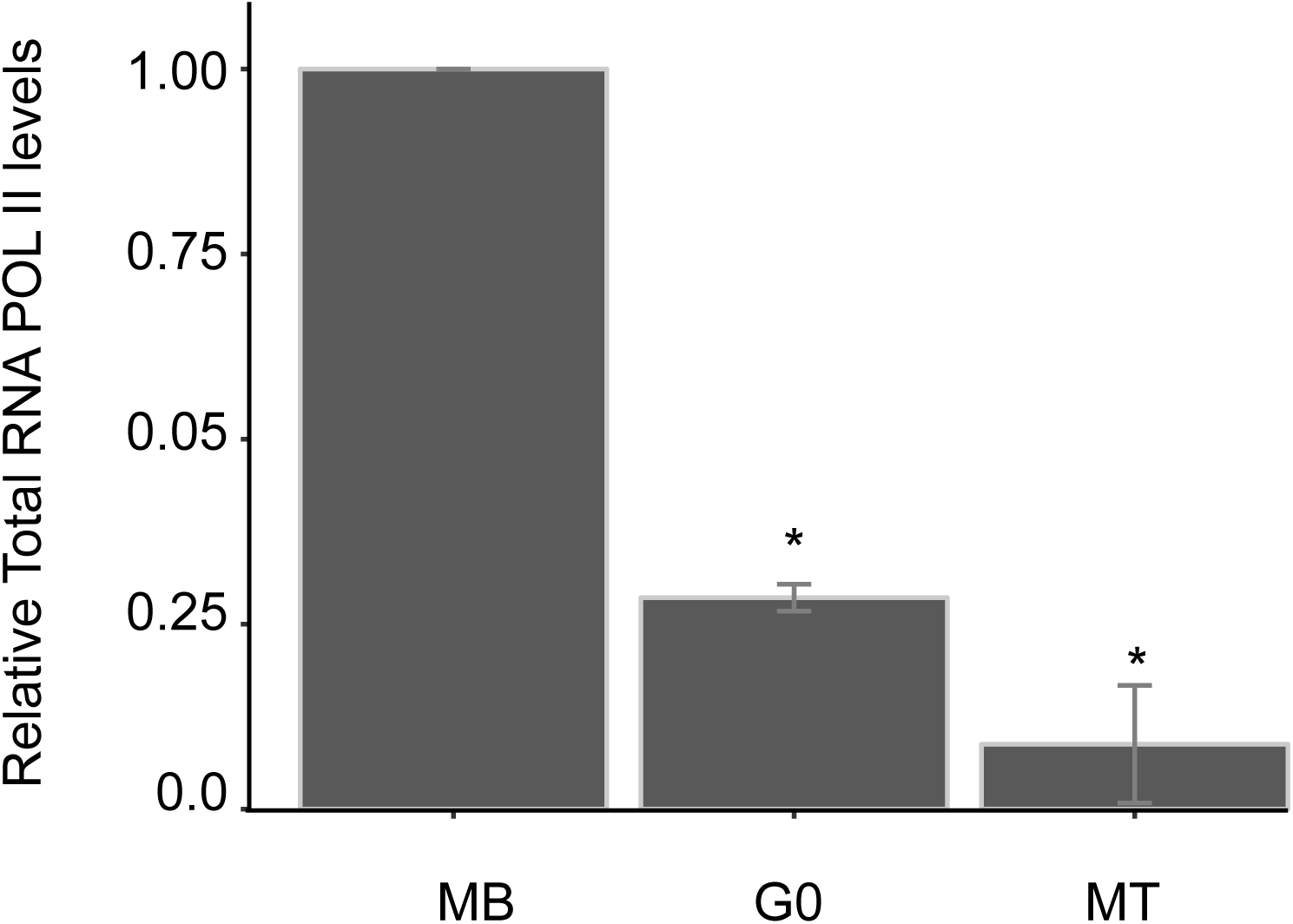
Reduction in levels of active RNA Pol II in G0 state: Densitometric analysis of western blot data shows an decreased proportion of RNA Pol II G0 and MT compared to MB. Normalized RNA Pol II levels referenced to MB and GAPDH as loading control for each state.

**Figure 2-figure supplement 2.**
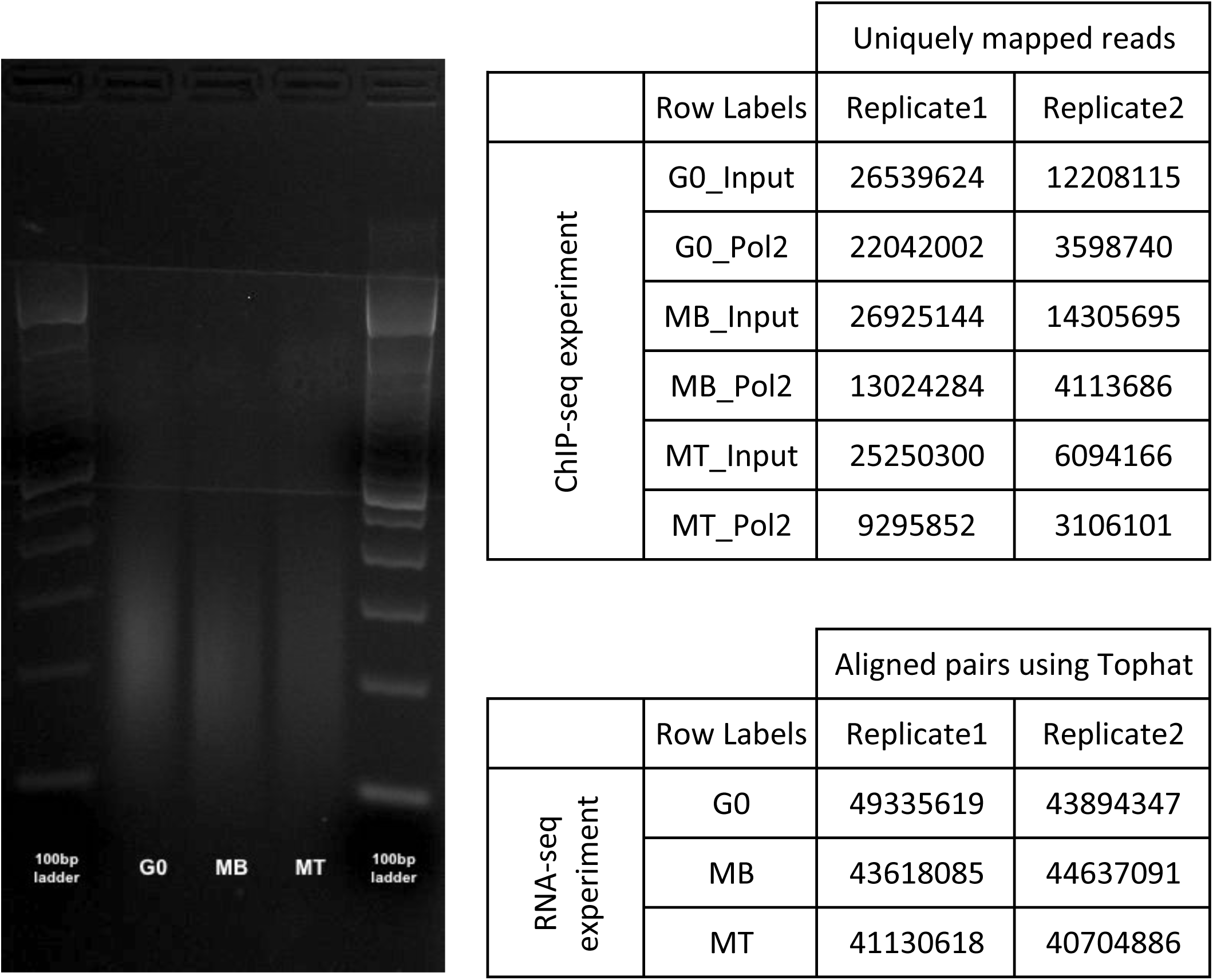
DNA gel depicting sonication profiles for all 3 states, the average sonication size is observed to be around 200 bases pairs. The table enumerated the sequencing depth of libraries of each replicate used in this study.

**Figure 2-figure supplement 3.**
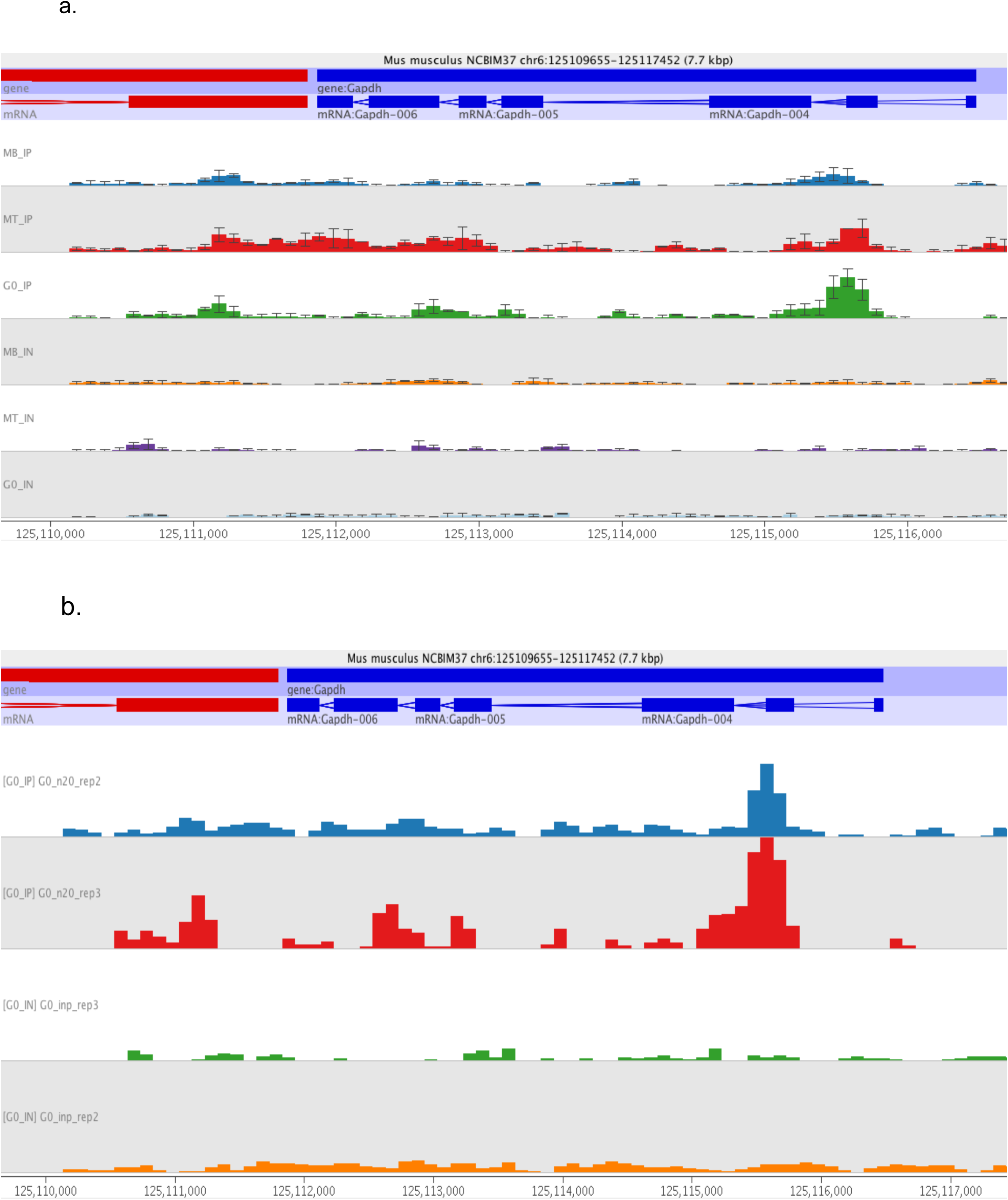
a. Mean Sequencing coverage (2 replicates) at GAPDH gene shown for all three cellular state (pull down and input). Error bar show spread between highest and lowest value observed across replicates Individual replicate track for G0 shown in panel below (b).

**Figure 2-figure supplement 4.**
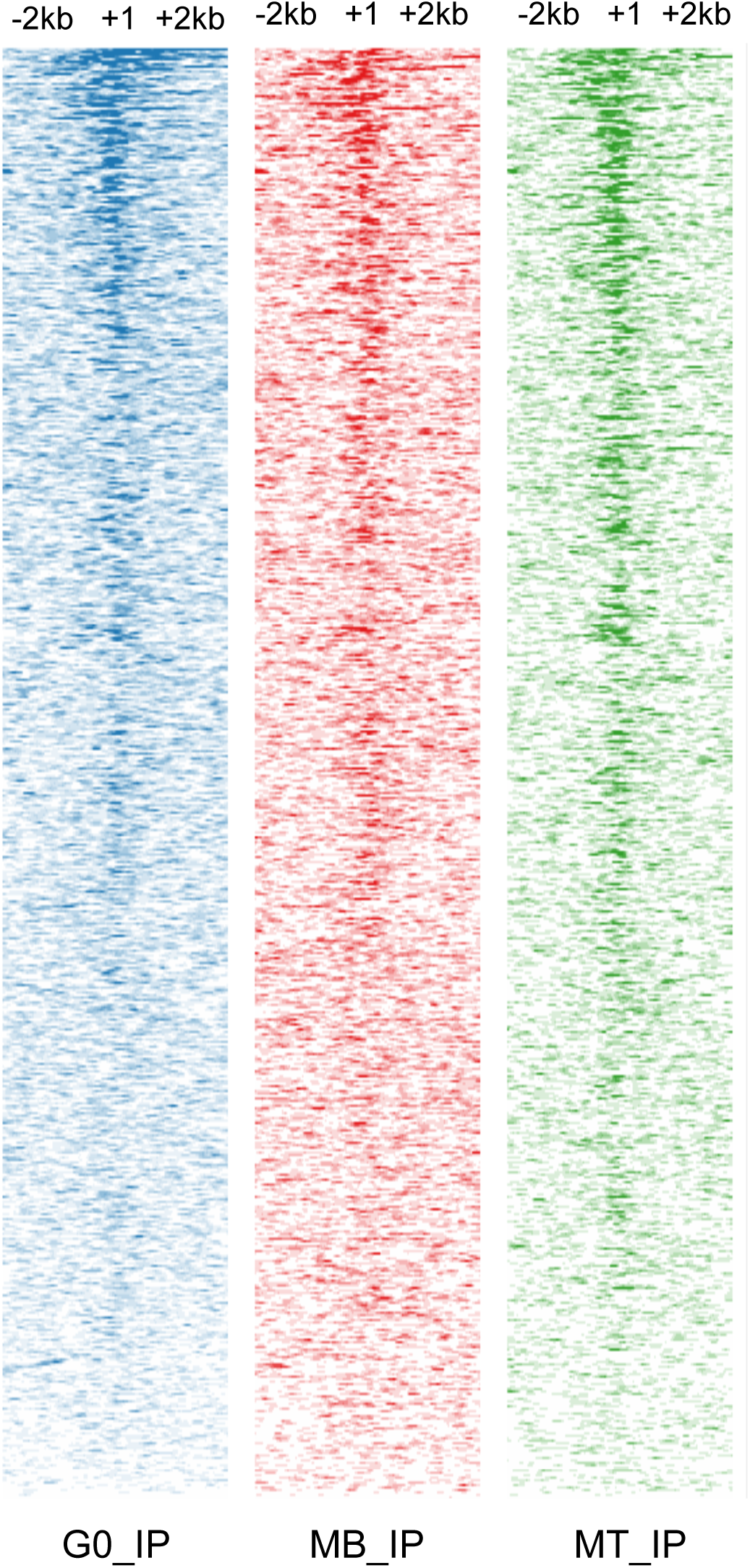
RNA pol II read density across TSS: Intensity of RNA pol II enrichment is represented around TSS (+/-2kb) for non overlapping genes. The dark shade represent higher intensity as observed in the ChIPseq data set for all three cellular states.

**Figure 2-figure supplement 5.**
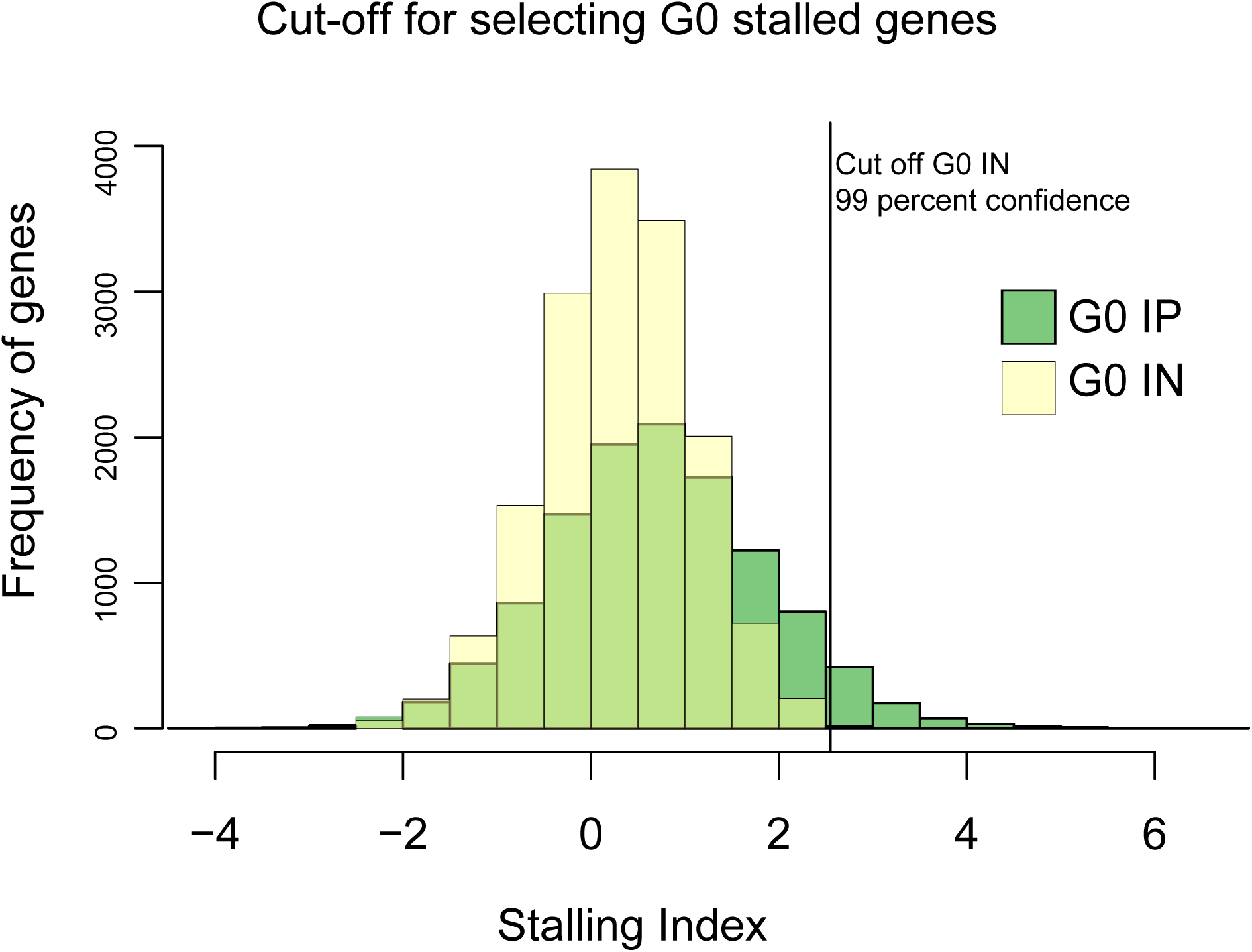
For calculation of stalling index the input is used as reference to set stringent cutoffs. The pulldown sample has rightward tail of the stalling index compared to Input and non overlapping region with >95percent confidence is used to identify the stalled gene. The vertical line shows the cut off value for G0 sample.

**Figure 2-figure supplement 6.**
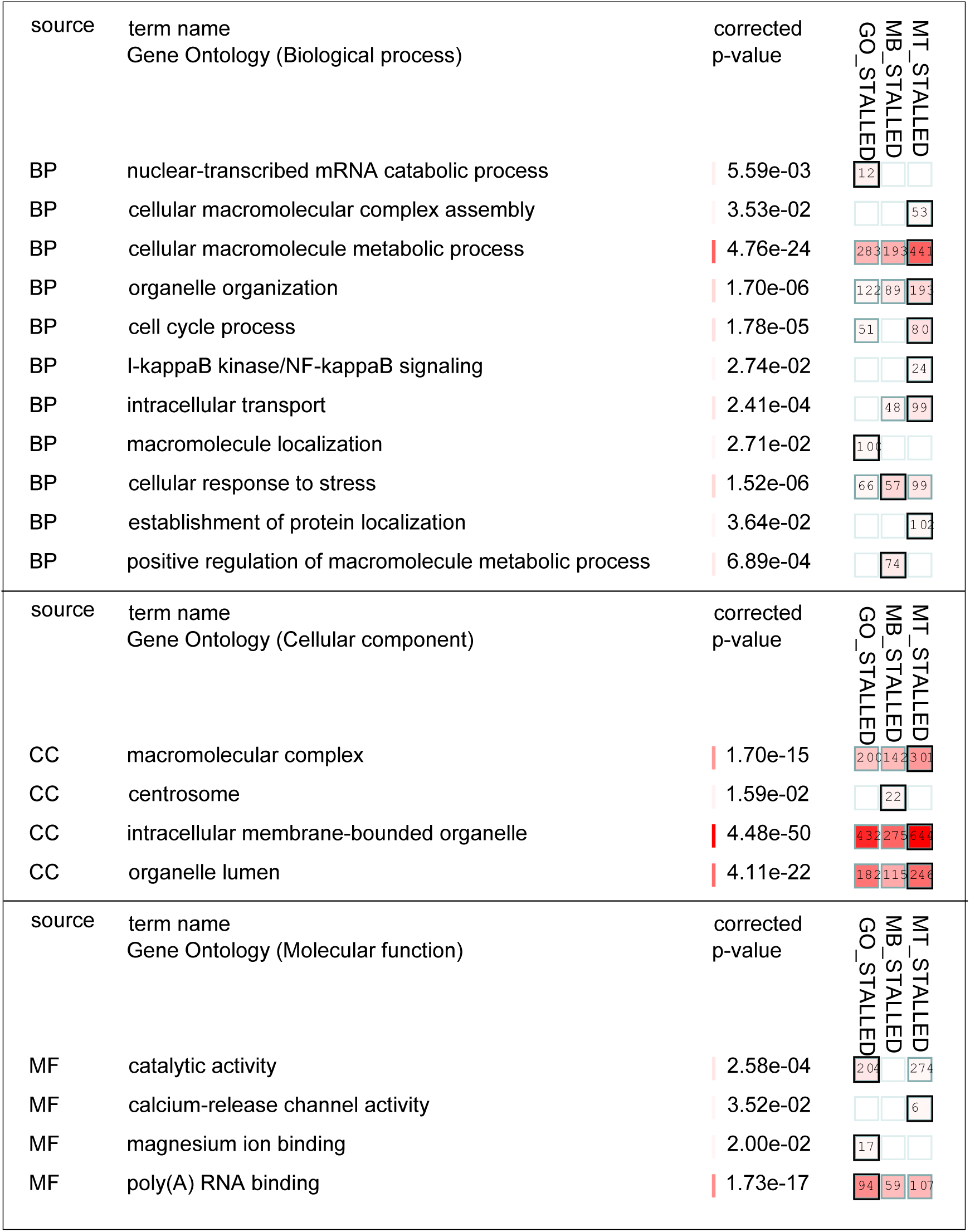
Gene Ontology analysisof Stalled genes Genes with stalling index >2.5 were enlisted for each MB, G0 and MT states Genes with stalling index >2.5 were enlisted for MB, G0 and MT states separately. gProfiler separately. gProfiler based comparison for the identified list was carried out. The based comparison for the identified list was carried out. The output is graphically represented here using “moderate” filtering option for Biological represented here using “moderate” filtering option for Biological process, Cellular componemt and molecular functions. The terms in each state is listed to the left. The square with red intensity indicates the significance of each states is The enriched terms in each state indicates the significance of each state. The most significant among the 3 highlighted with boundaries around square and corresponding p value is shown next to it.

**Figure 2-figure supplement 7.**
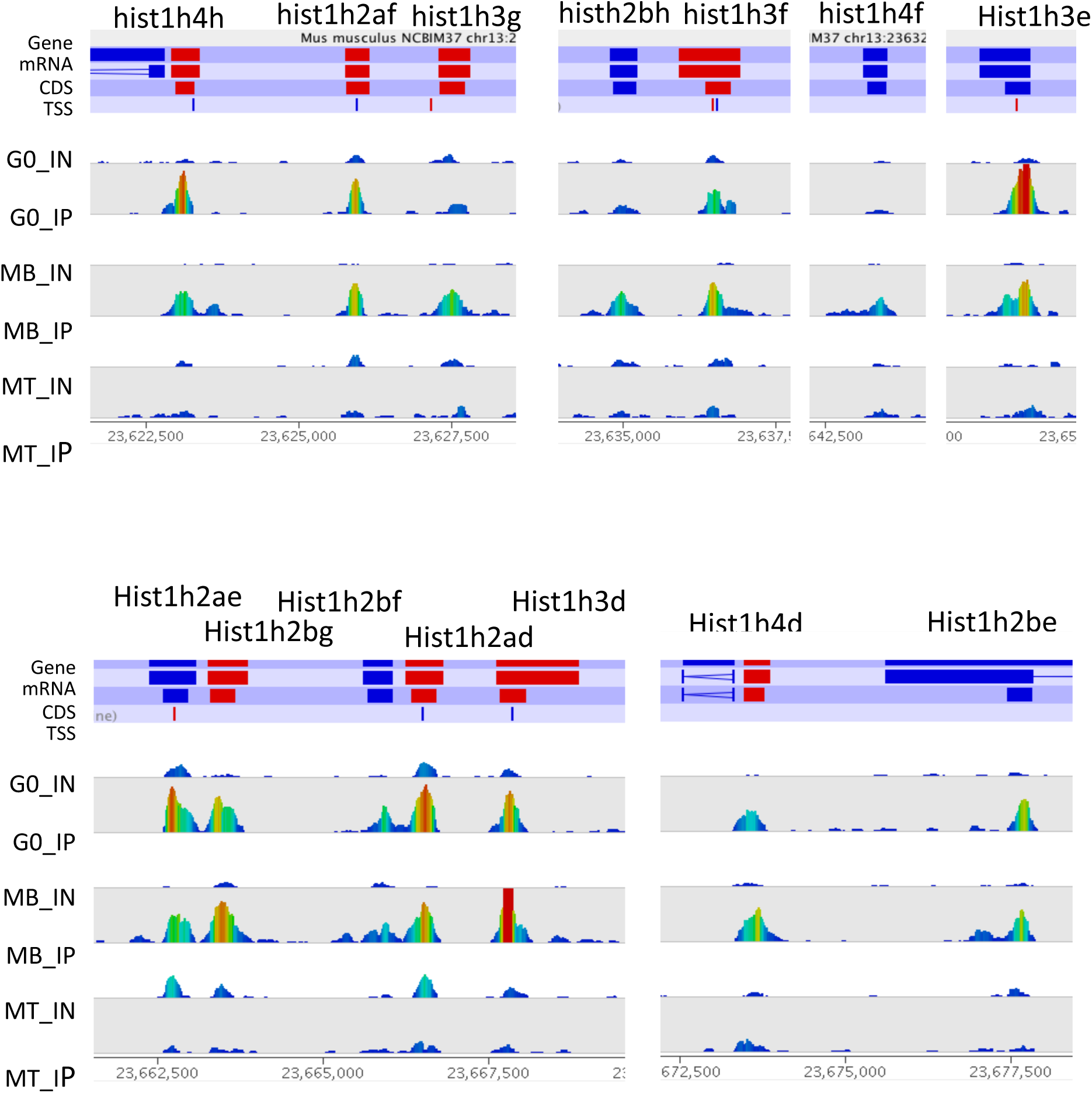
RNA Pol II abundance at histone 1 cluster is higher in G0 and MB and not in MT Genome browser snapshot was generated for the Histone 1 gene cluster at coordinates (chr 13:23622400-23677600) on mm9 build. The top panel shows the gene location on chromosome and its orientation (red indicates forward and blue in reverse). The read density is normalized per million within each sample and plotted with the color code where red signifies higher density and blue indicates lower density. The intergenic regions are removed in order to represent the density at higher resolution. Strong enrichment of Pol II seen in MB persists in G0 but not MT, despite down regulation of transcripts in both arrested states.

**Figure 3-figure supplement 1.**
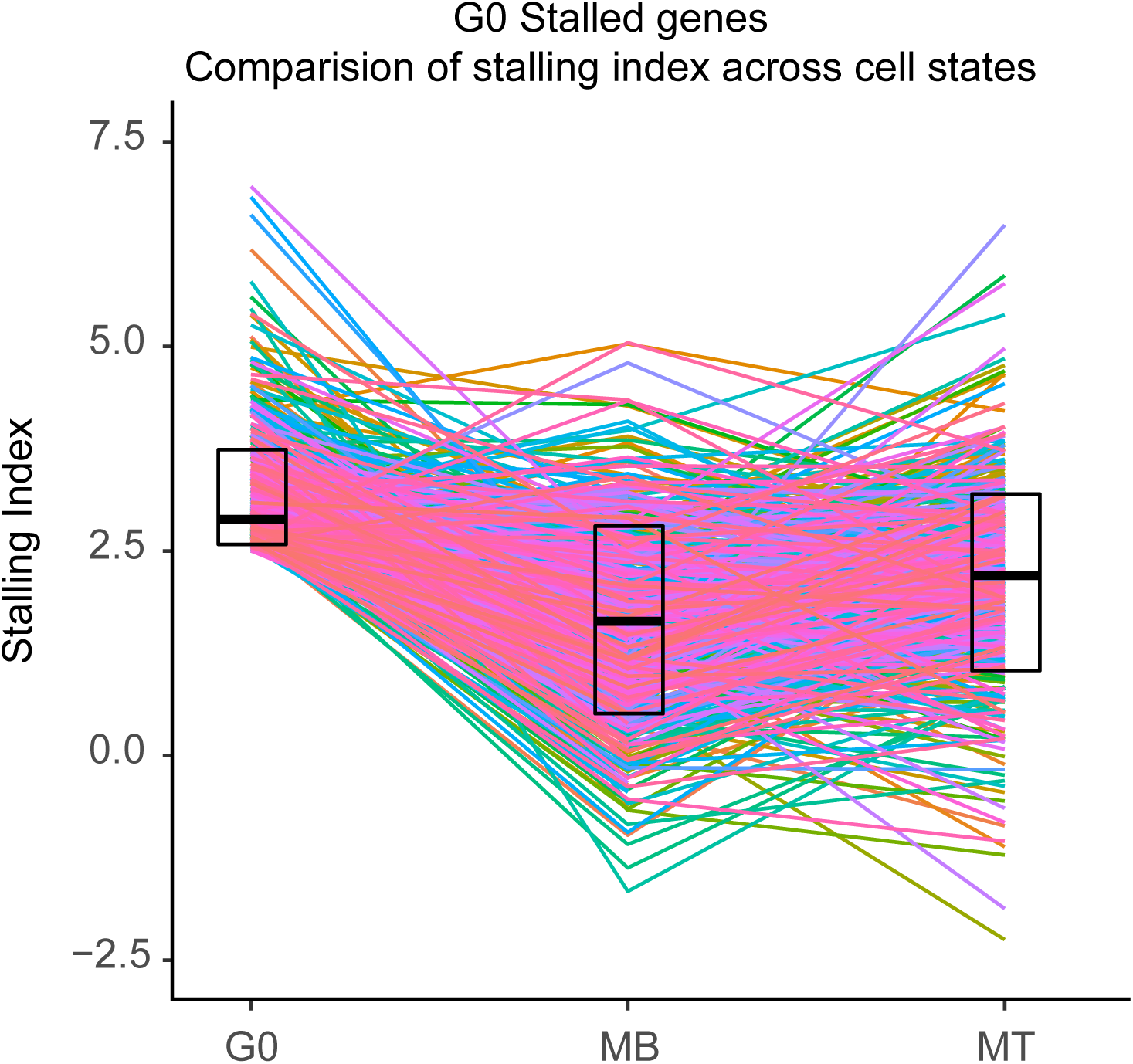
Stalling Index for G0 stalled genes for each of three cellular states. The nested box plot show the distribution for SI (25-75 percentile limits) and the lines trace the SI values across 3 states for each gene.

**Figure 3-figure supplement 2.**
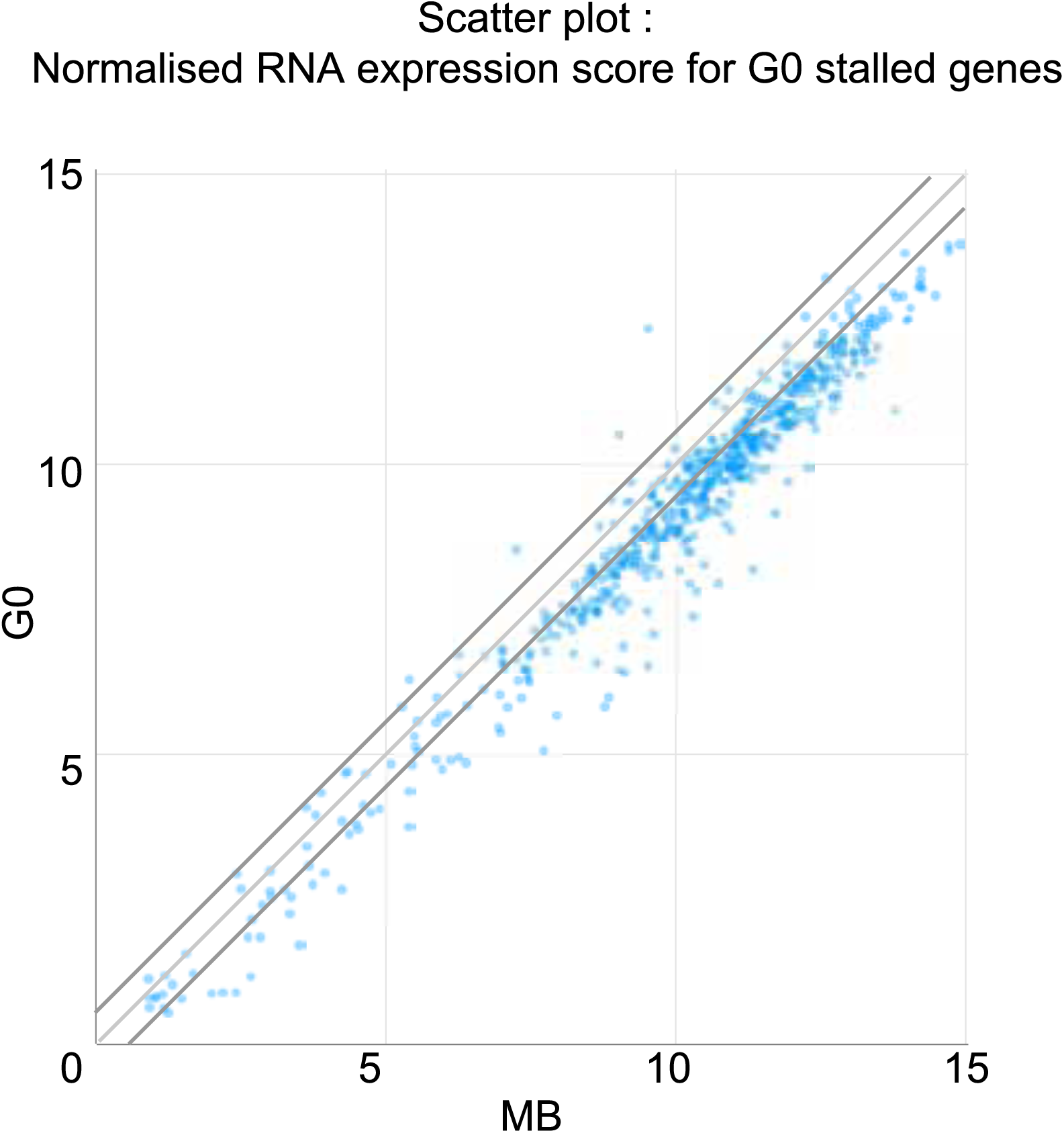
Cell number Normalized RNA levels observed for MB and G0 for set of G0 stalled genes. The thick line depicts the 2 fold change. The G0 stalled genes are observed to be expressed higher in MB than G0

**Figure 3-figure supplement 3.**
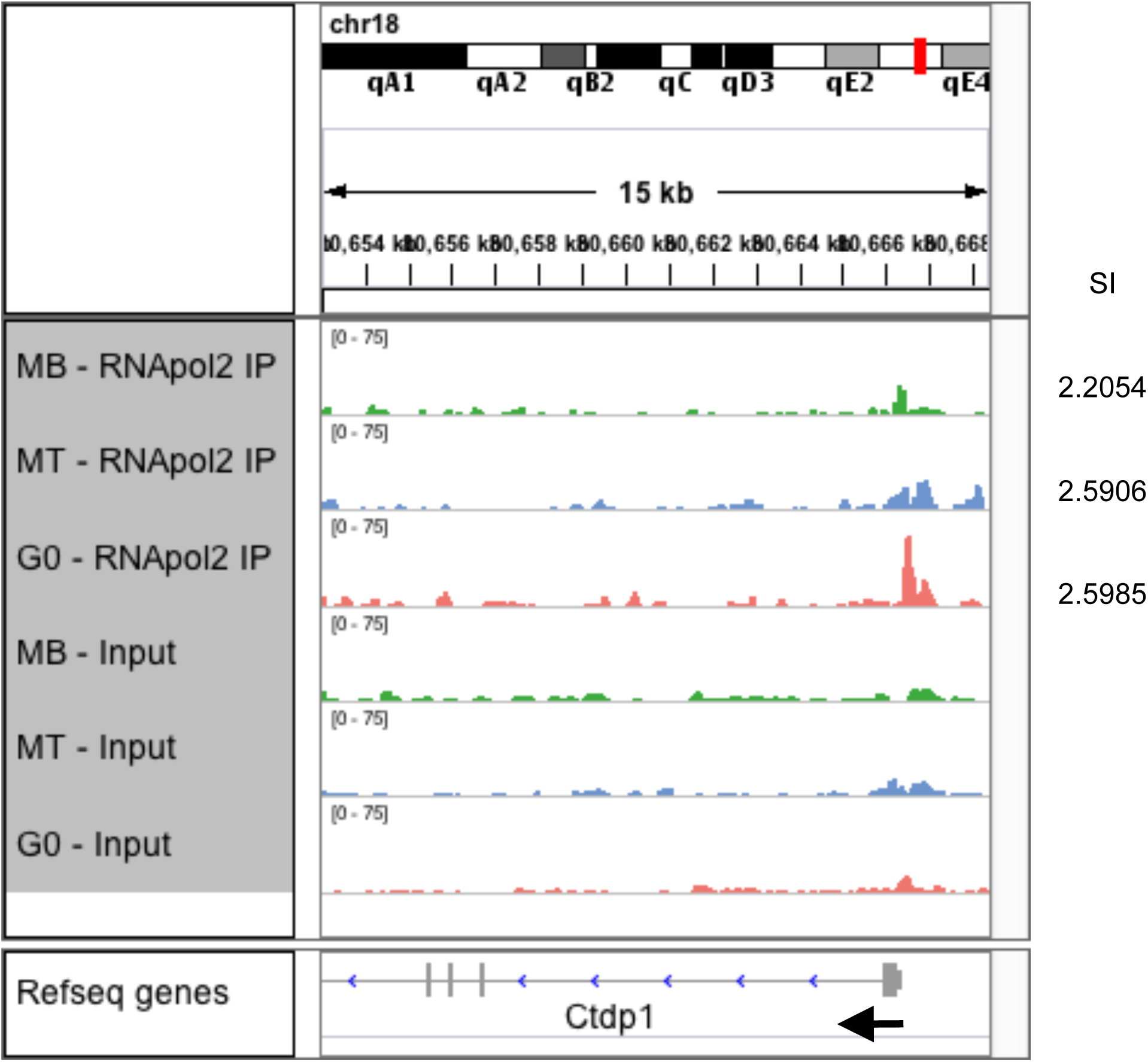
Ctdp1 locus: Genome browser snapshot depicting RNA pol II enrichments. Black arrow indicates the orientation of gene and numbers on right hand side indicate the observed stalling index in respective state.

**Figure 3-figure supplement 4.**
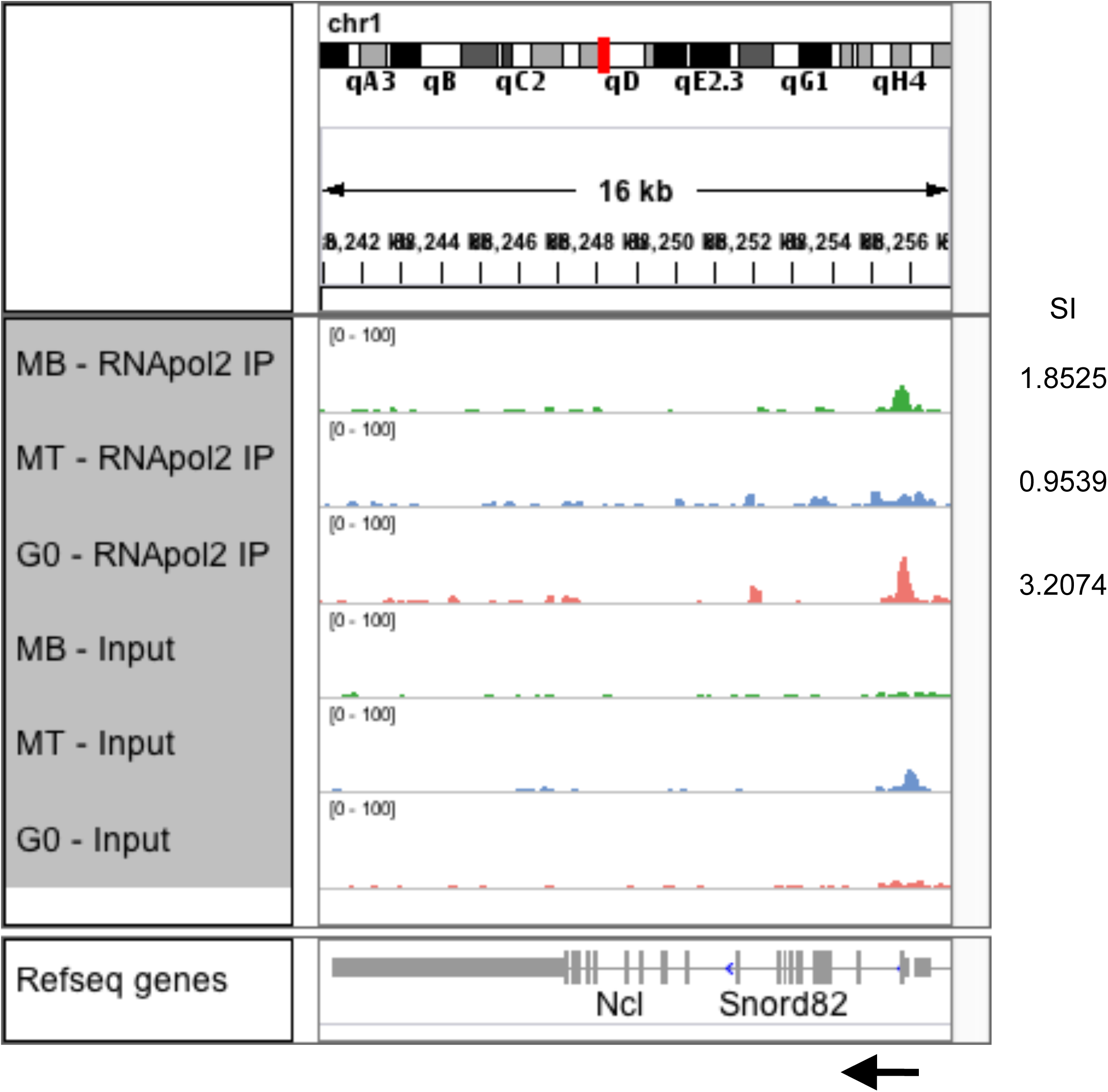
Ncl locus: Genome browser snapshot depicting RNA pol II enrichments. Black arrow indicates the orientation of gene and numbers on right hand side indicate the observed stalling index in respective state. Snord is intronic transcript in the Ncl locus.

**Figure 3-figure supplement 5.**
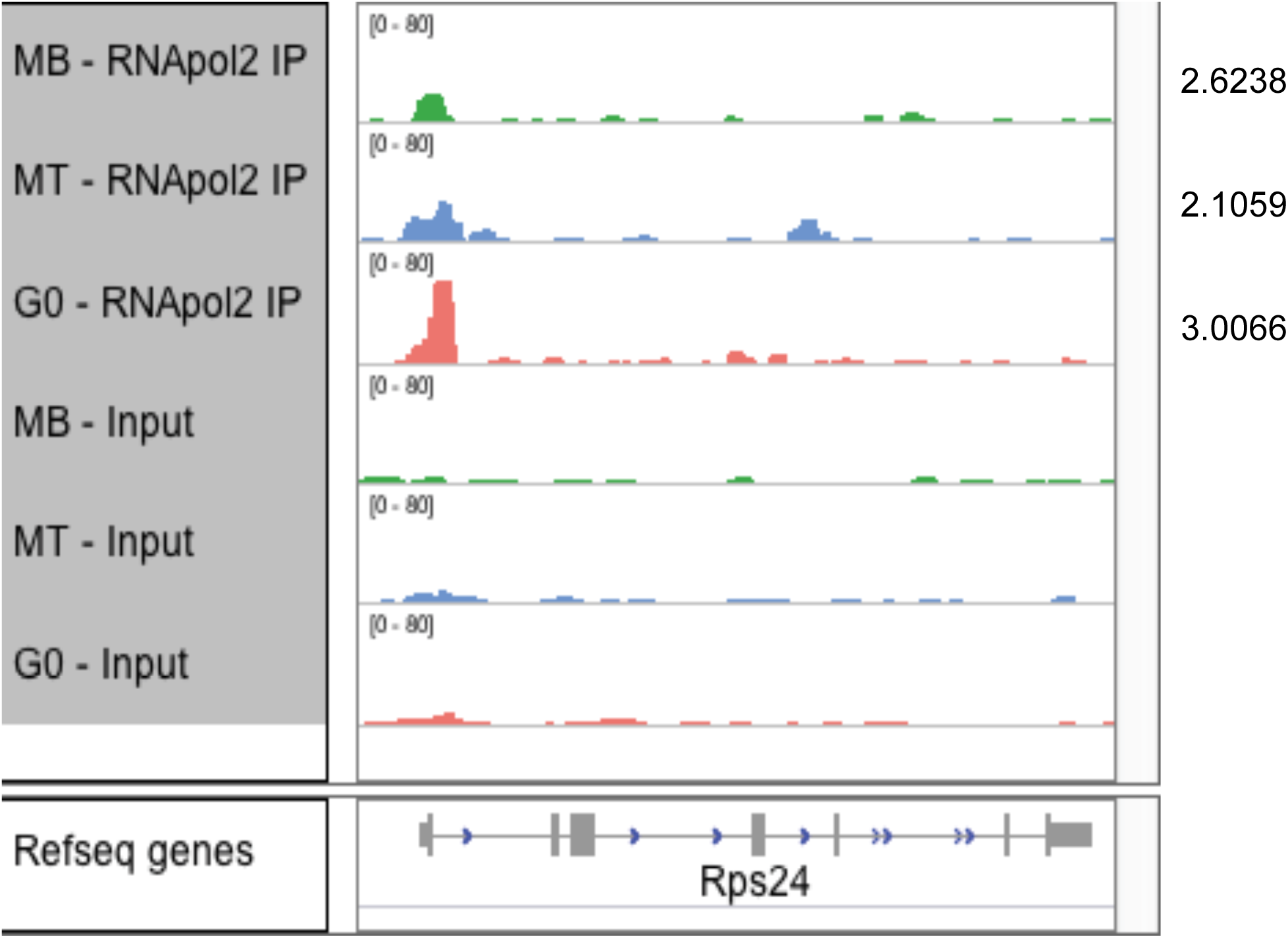
Genome browser snapshot for Rps24, a representative G0-stalled gene. The read density is normalized per million within each sample and plotted at equal scale.

**Figure 3-figure supportplement 6.**
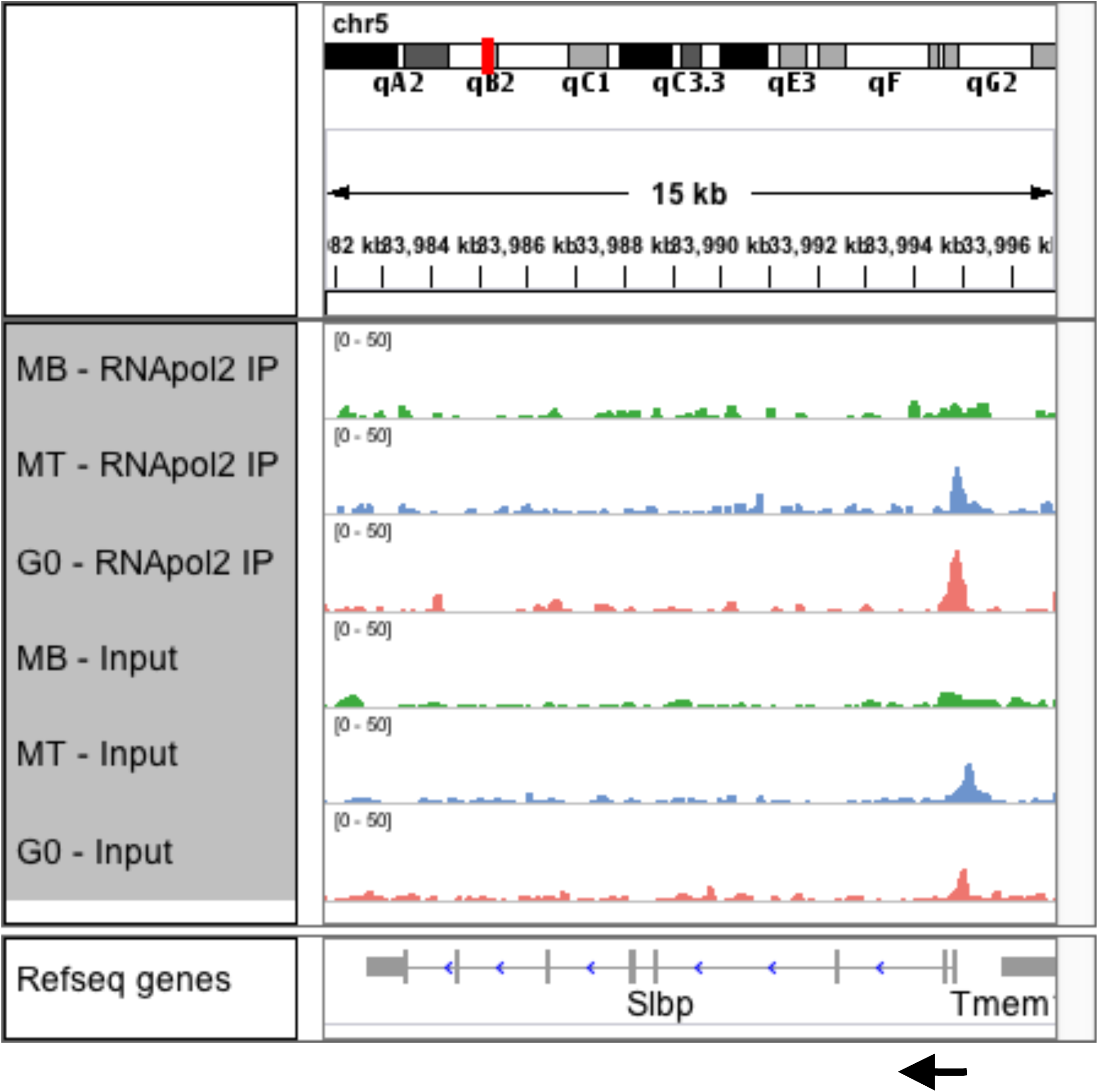
Slbp locus: Genome browser snapshot depicting RNA pol II enrichments. Black arrow indicates the orientation of gene.

**Figure 3-figure supplement 7.**
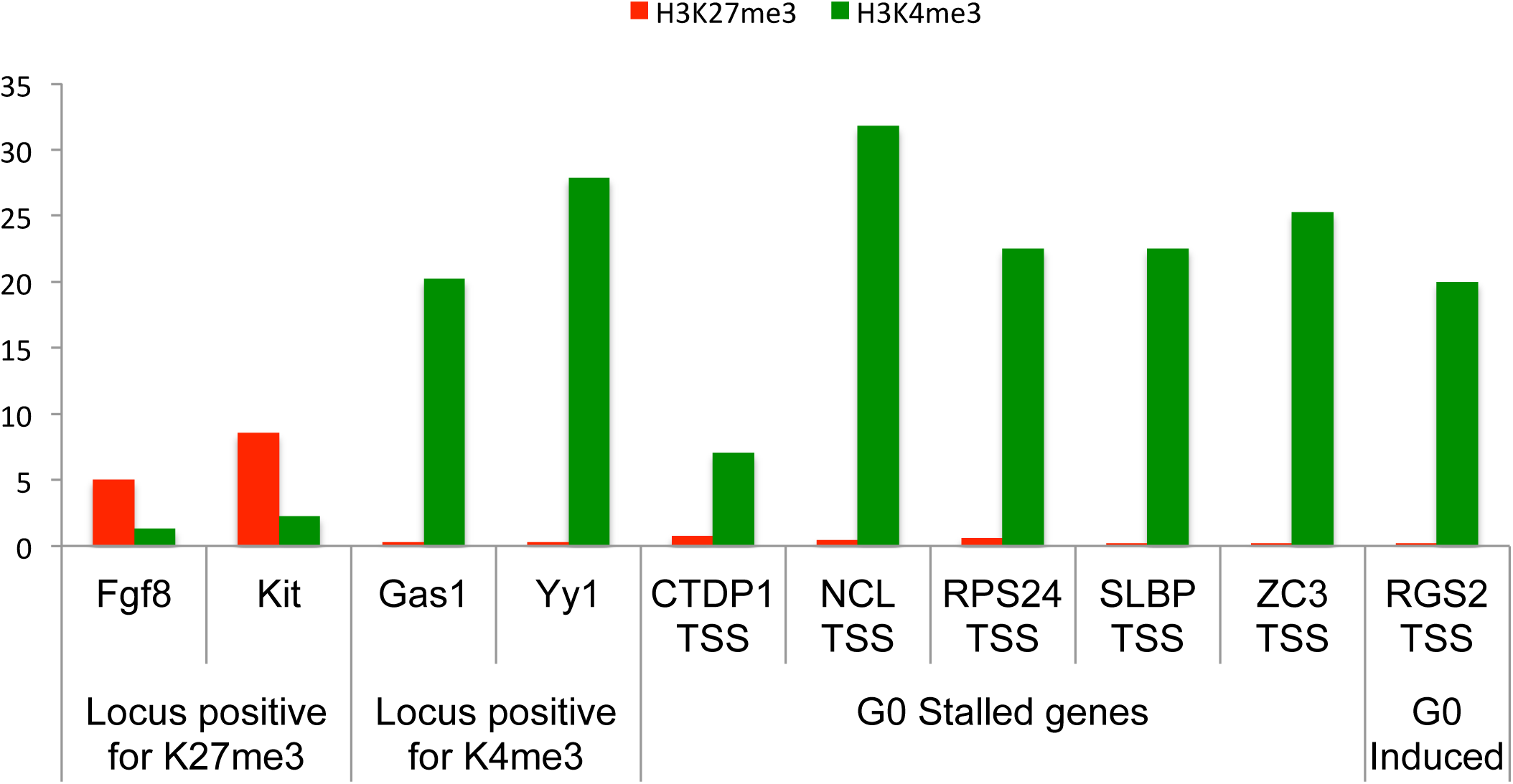
G0 stalled genes (CTDP1, NCL. RPS24, SLBP, ZC3h12a) are marked by H3K4me3 mark but not the H3K27me3 suggesting that they are locus with open chromatin. The Positive controls for only H3K4me3 (Gas1 and Yy1 loci) and H3K27me3 (Fgf8 and Kit loci) are shown for reference.

**Figure 3-figure supplement 8.**
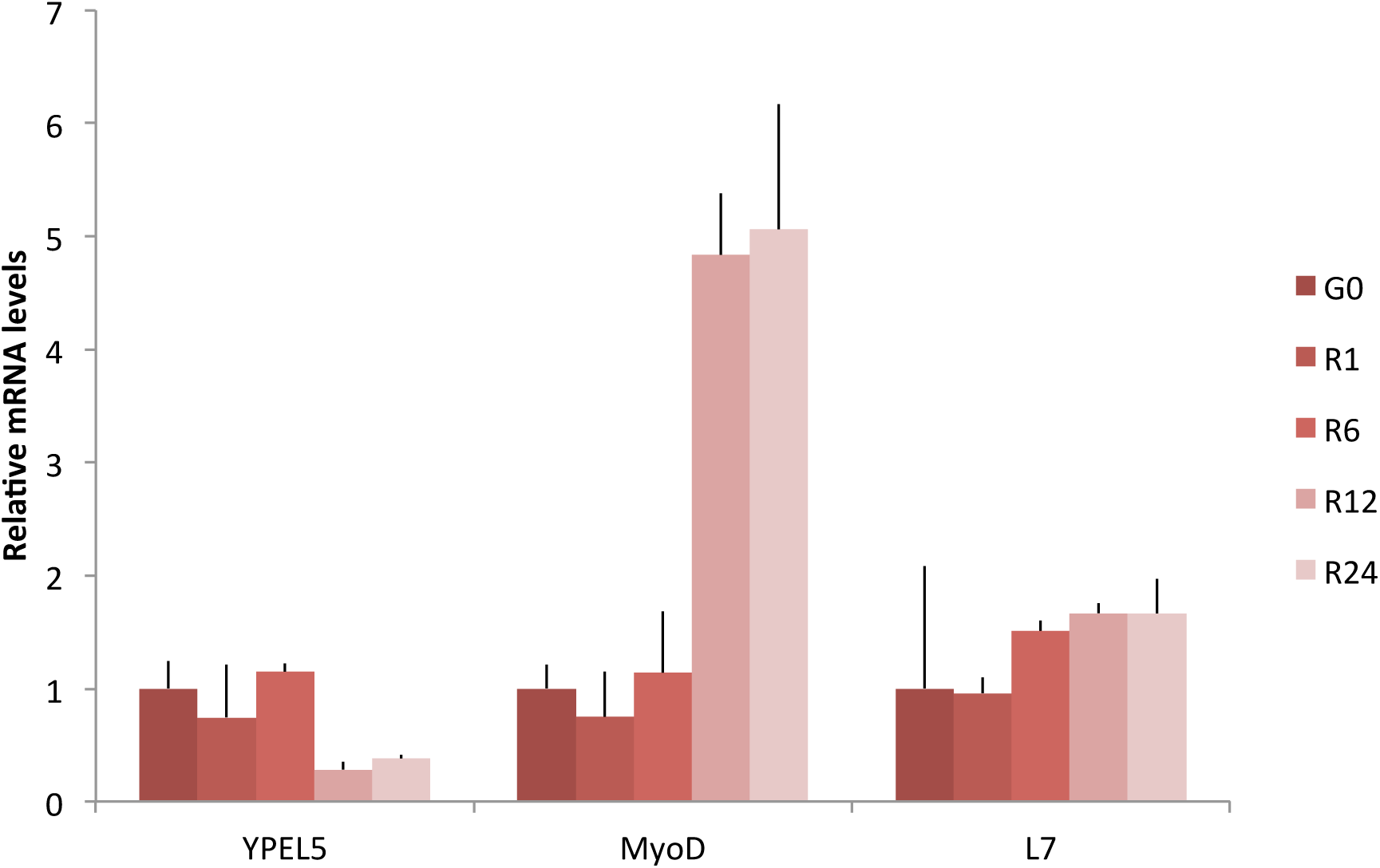
Relative mRNA levels for group of genes repressed and not stalled in G0

**Figure 3-figure supplement 9.**
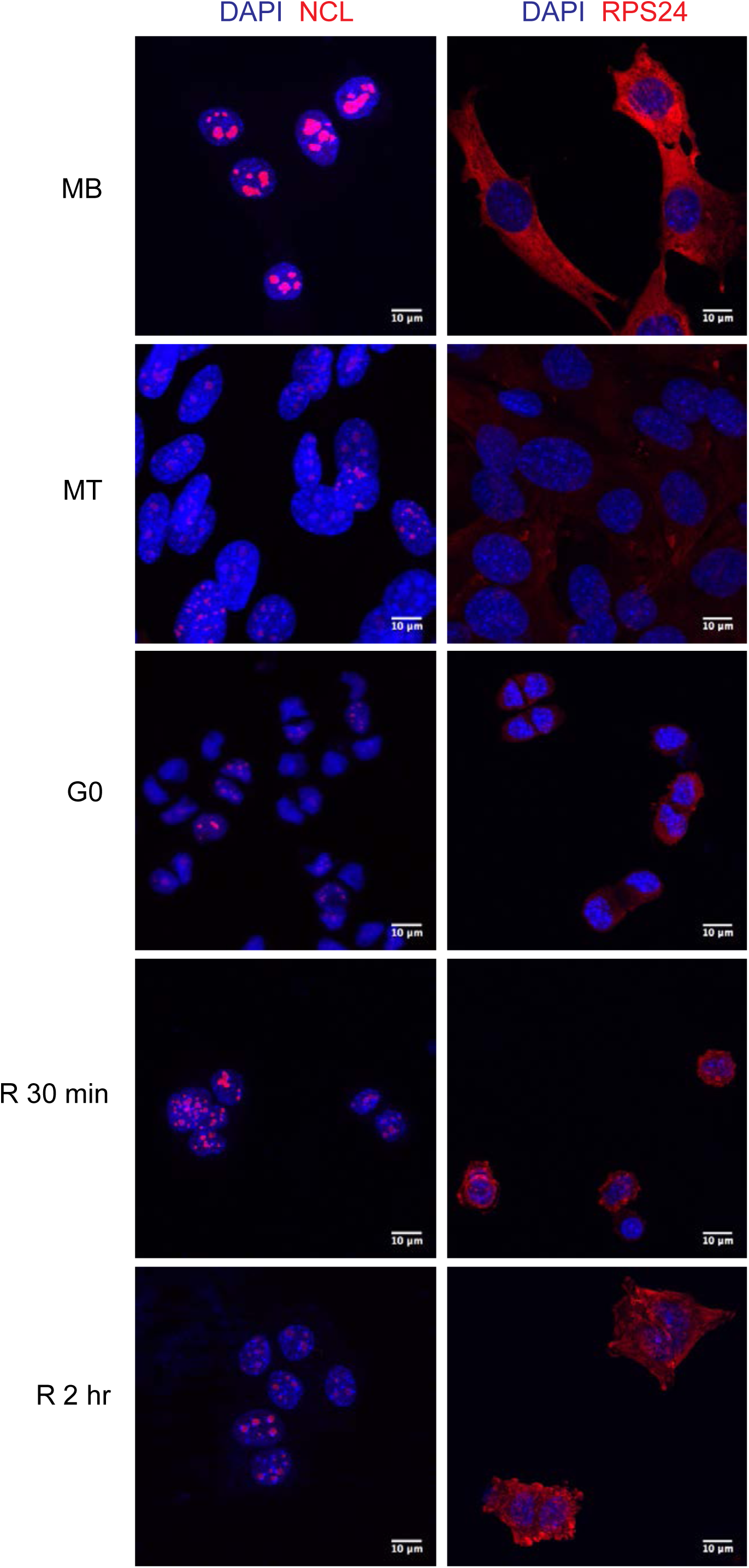
Restoration of expression of G0-stalled genes during cell cycle reactivation. Representative immunofluorescence images for MB, MT, G0, R30min and R2hrs stained for G0 stalled genes (red) and DAPI (Blue). Ncl (left column) shows distinct sub-nuclear puncta (nucleolar) staining whereas and Rps24 (Right column) is found in the cytoplasm. For both these G0 stalled genes MB shows maximum protein expression, whereas the two arrested states (G0 and MT) show very reduced levels. Expression is rapidly induced after reactivation (R30min and R2hrs).

**Figure 3-figure supplement 10.**
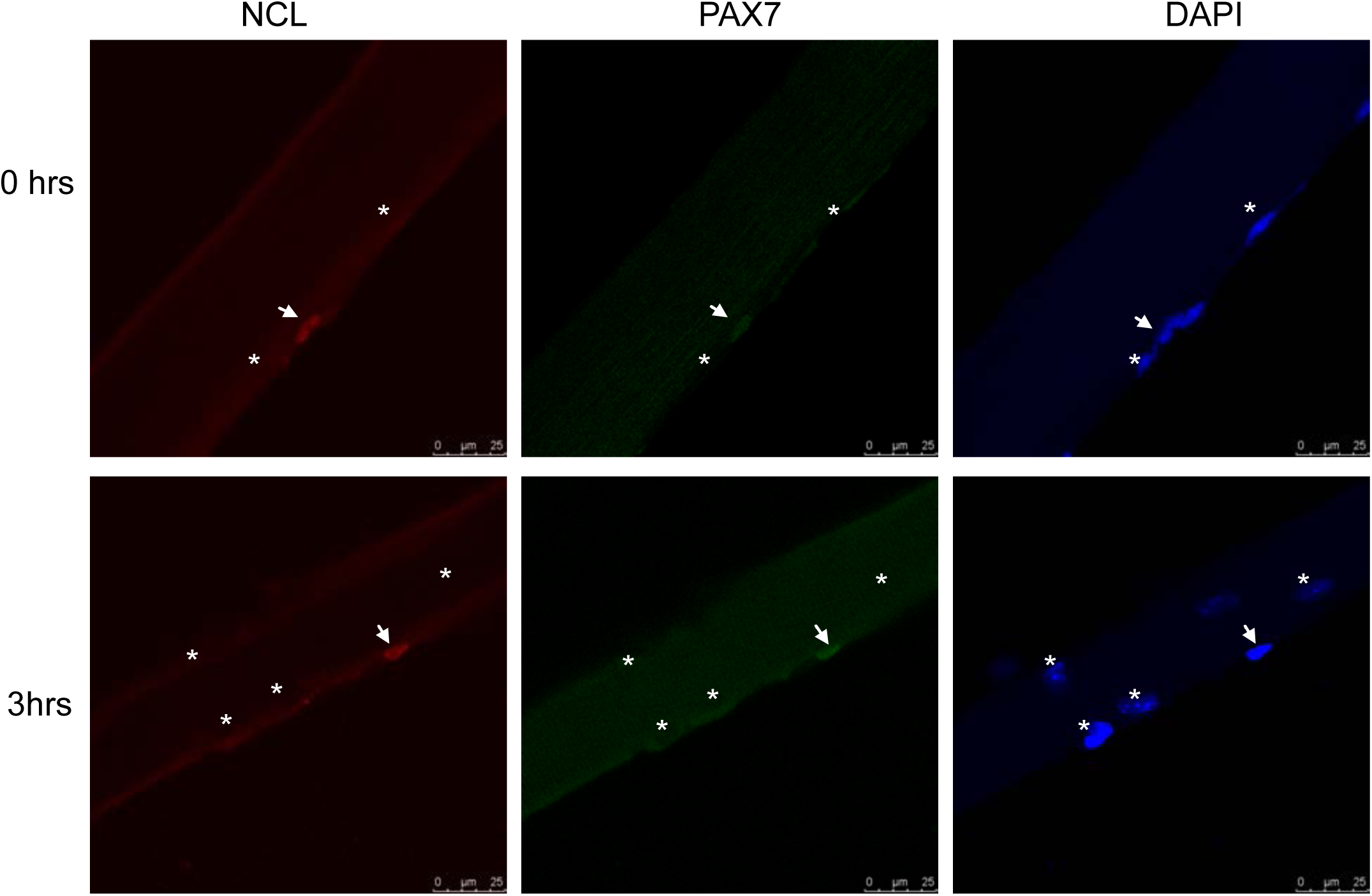
Ncl staining in satellite stem cells (SC) associated with single myofibers cultured ex vivo. SC are marked by Pax7 (arrowhead) can be distinguished from differentiated myonuclei which are Pax7 negative (MN, asterisk) within the underlying myofiber. At 0hrs and 3hrs post isolation, SC shows induced Ncl levels whereas MN shows little staining.

**Figure 3-figure supplement 11.**
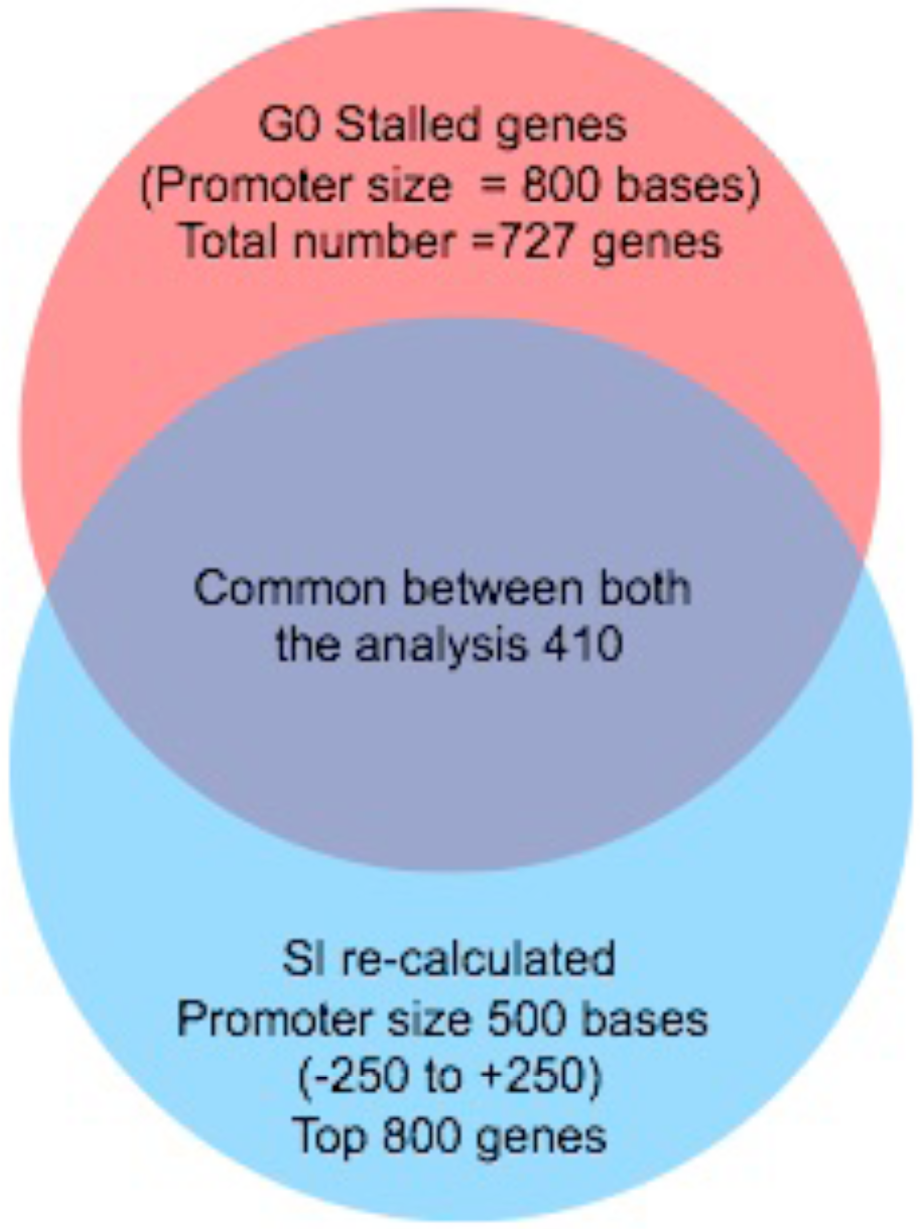
Venn diagram depicting the overlap of G0 stalled genes when calculated with varying the Promoter region (+500 bases or +250 bases)

**Figure 4-figure supplement 1.**
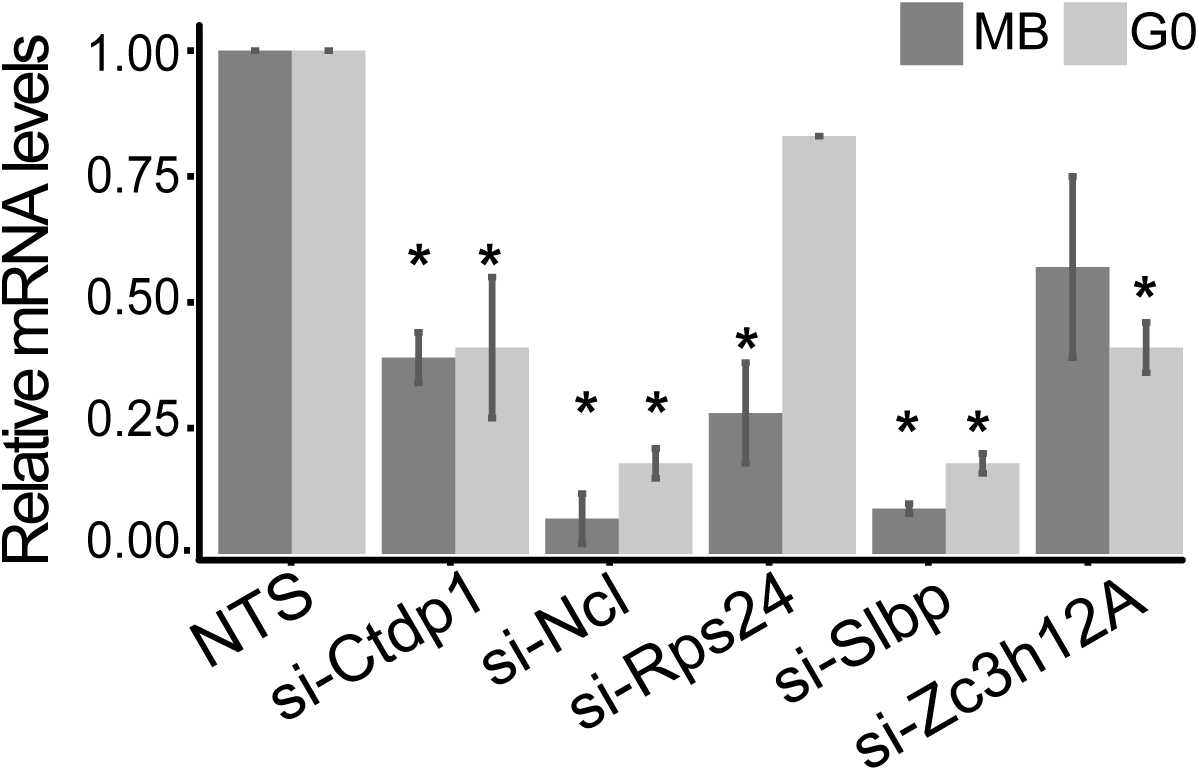
Extent of knockdown of respective mRNA levels after siRNA treatment in MB and G0 estimated using q-PCR (normalized to GAPDH within the treatment and corresponding NTS value) (Values represent mean + SEM n=3, p value <0.05).

**Figure 4-figure supplement 2.**
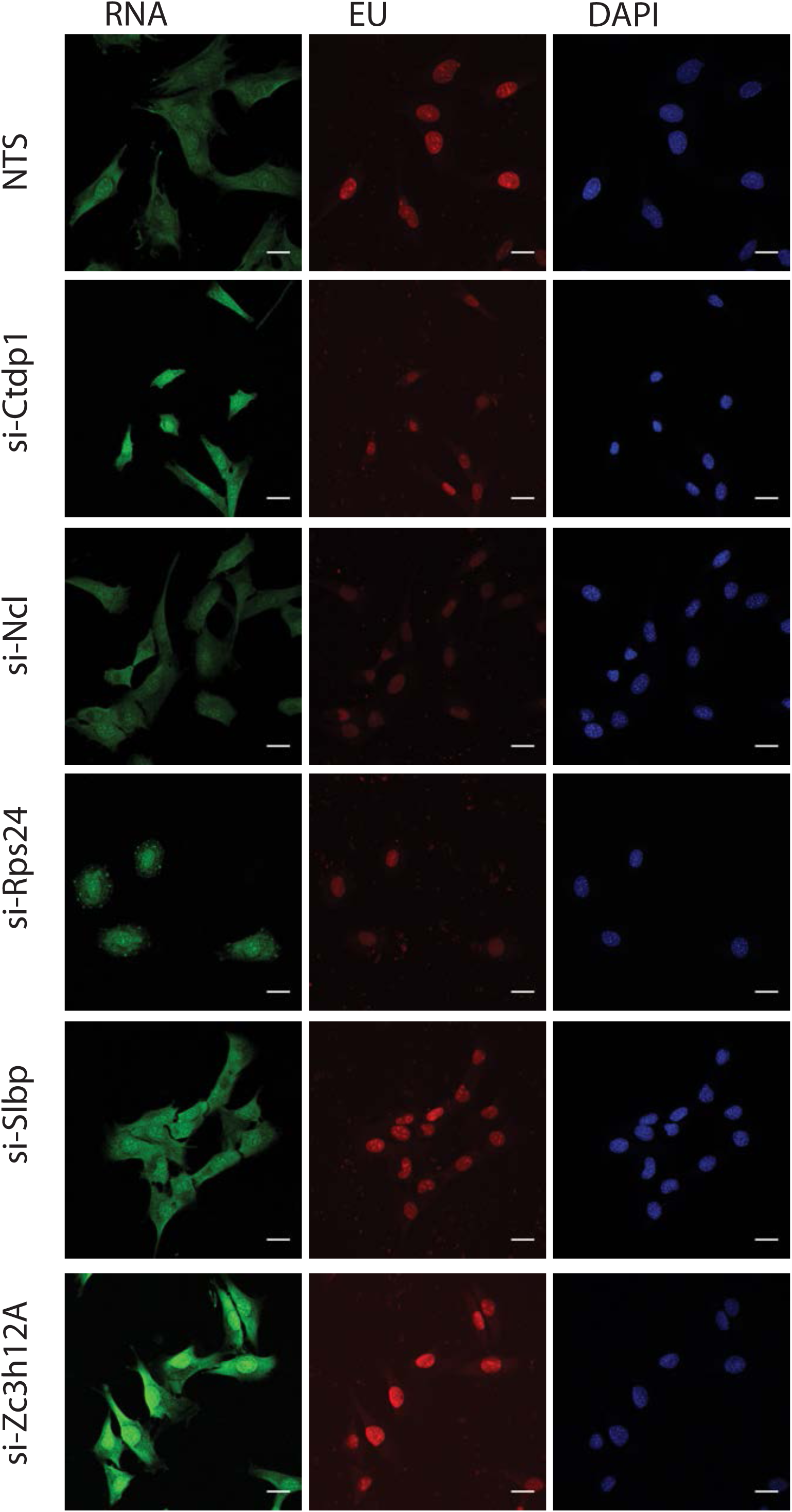
RNA content, active RNA synthesis and nuclear area altered in proliferating MB after siRNA-mediated knockdown of candidate G0-stalled genes (NTS-non targeting control siRNA, Ctdp1, Ncl, Rps24, Slbp and Zc3h12A). After 36hr, cells were co-stained for total RNA (SYTO^®^ RNASelectTM - green) and EU incorporation following a 30‵ pulse (red) (scale bar 10μm).

**Figure 4-figure supplement 3.**
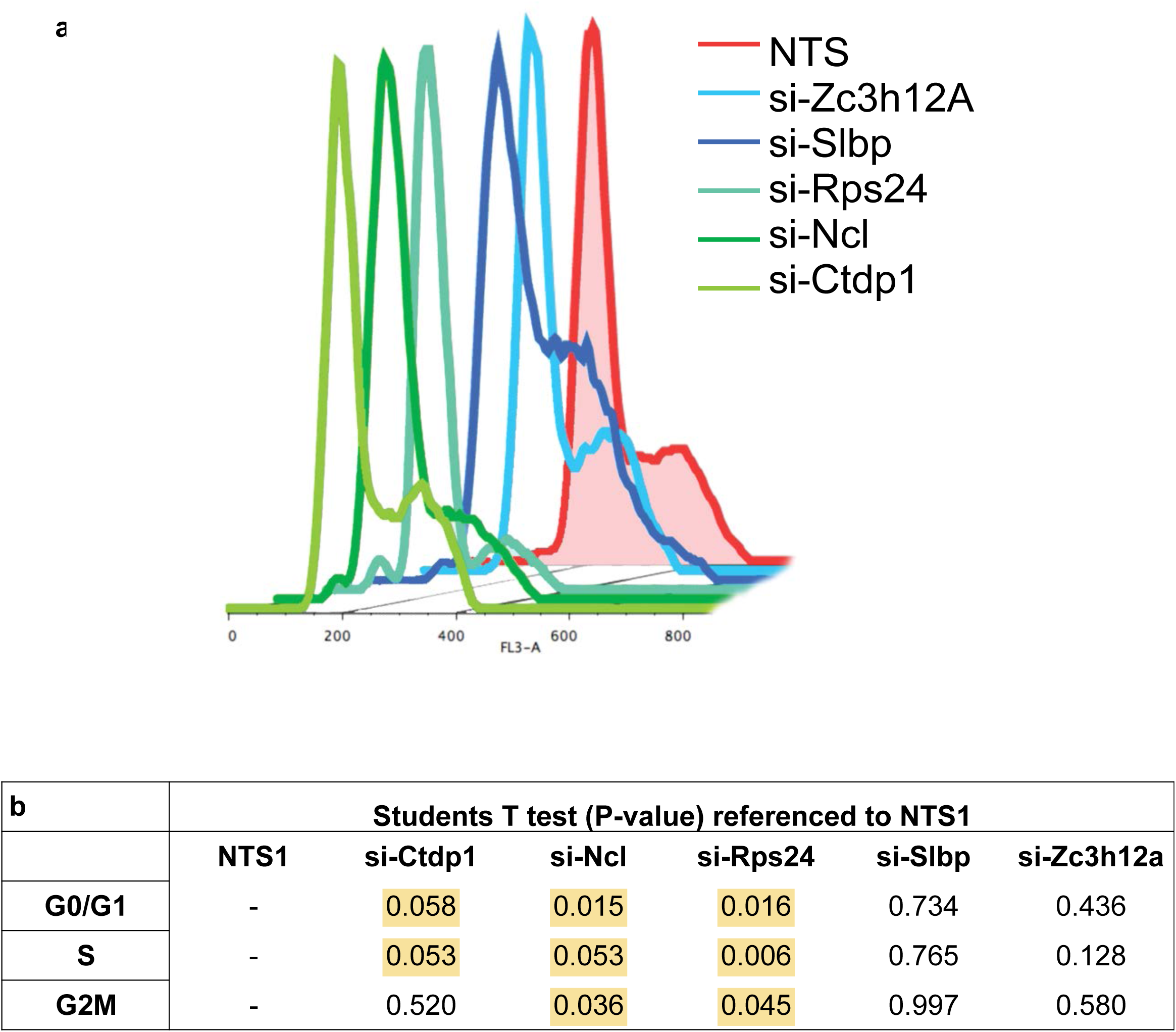
a. Overlayed DNA content traces plotted for the NTS (red line) treated MB. Knockdown of Ctdp1, Ncl and Rps24 show G1 arrest (green shades), for whereas Slbp and Zc3h12A are unchanged (blue shades). The quantification of proportion of cells in each cell cycle state is represented in Figure 4 b and the significance is tabulated in b and highlighted in yellow.

**Figure 4-figure supplement 4.**
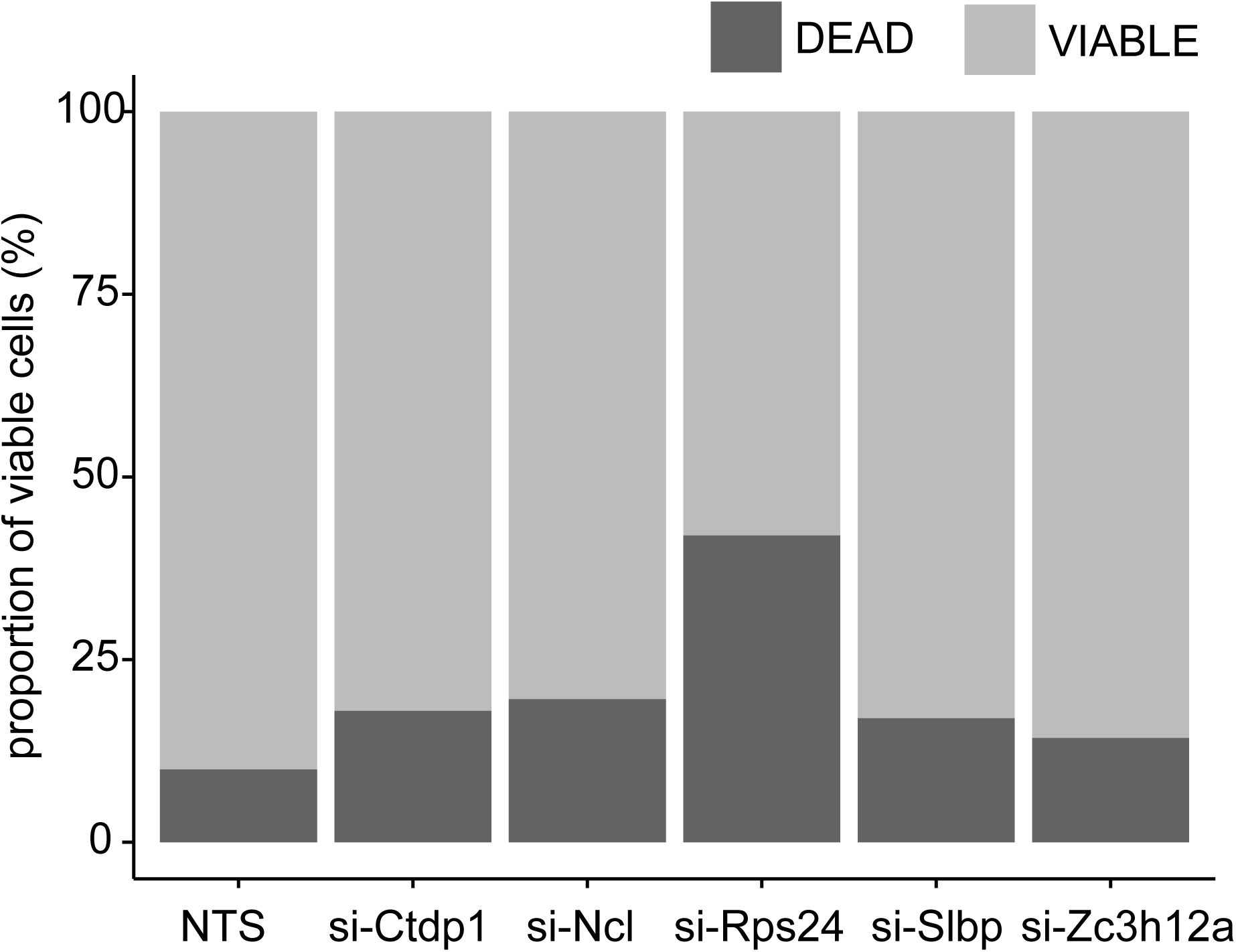
Propidium iodide staining of live cells shows a modest decrease in cell viabity for all the siRNA treated cells maximally Rps24

**Figure 4-figure supplement 5.**
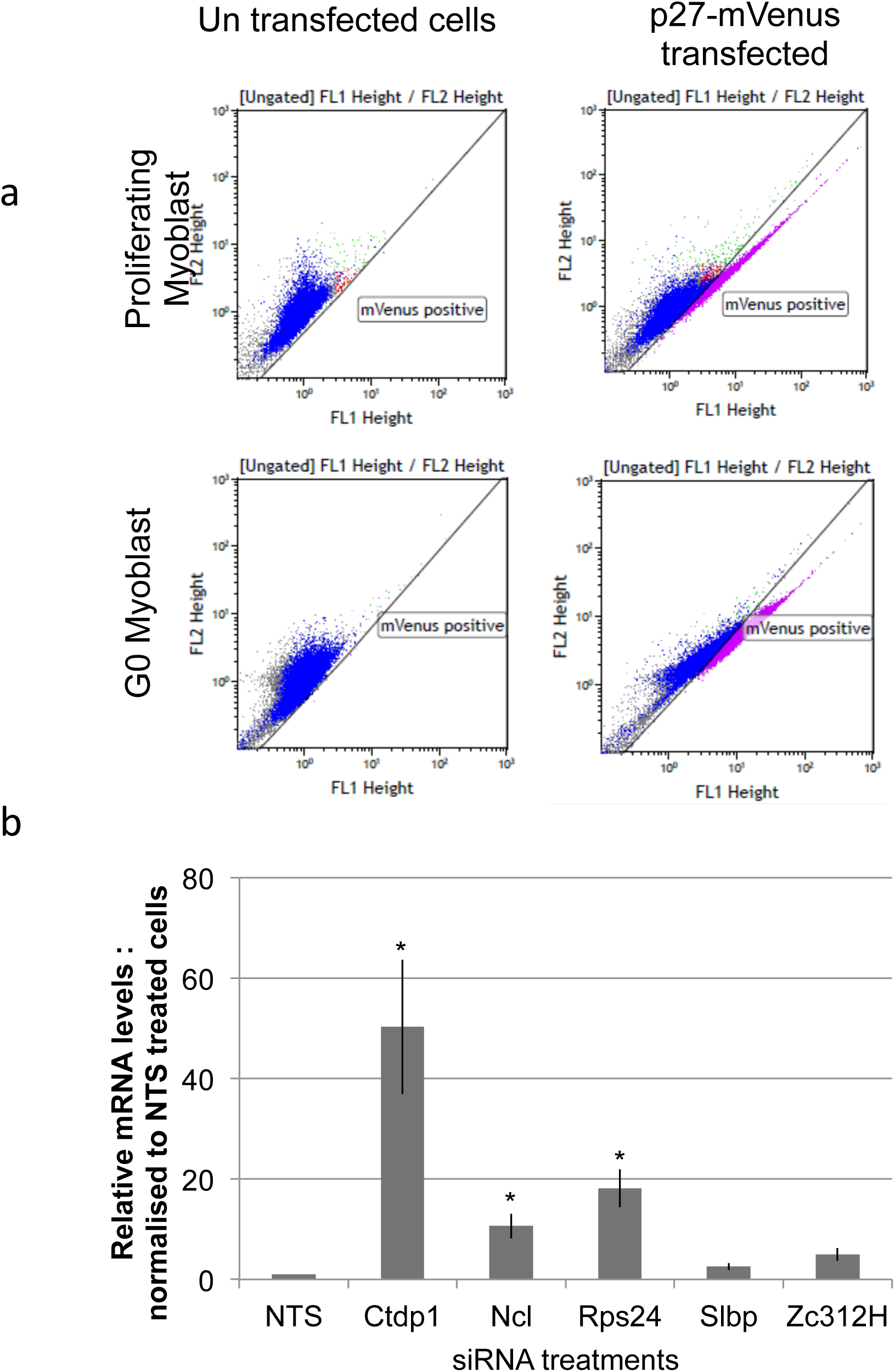
Proportion of P27 positive cells increase in G0 compared to MB and is good indicator for quiescence state. a. Stably expressing P27-Venus (fl-1) plasmid in C2C12 cell line (right side panel), the proportion of P27 positive cells is estimated by flow cytometry. The un- transfected C2C12 MB is used to set reference gates(Left side panel). b. The bar graph below show relative levels of mRNA of p27 marker upon knockdown with G0-stalled genes compared to NTS control during proliferative growth conditions. Knockdowns of Ctdp1, Ncl, and Rps24 show significant induction in p27,a marker of quiescence. (* =pvalue <0.05 Students T-test, N=3)

**Figure 4-figure supplement 6.**
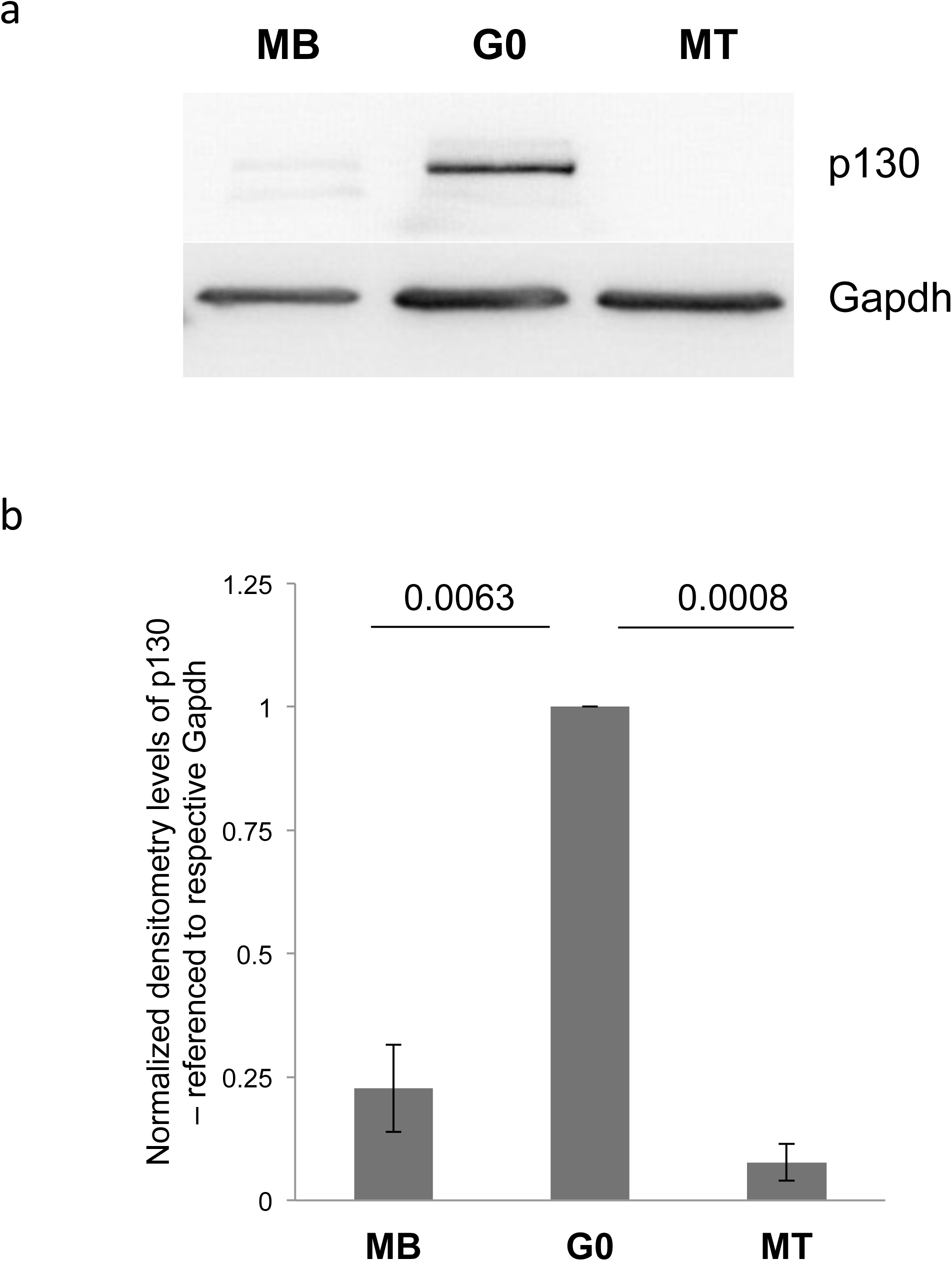
P130 levels as marker for quiescence state Western blots (b) for P130 and Gapdh and densitometry quantification (b) shows G0 specific induction of p130. MB and MT show significantly lower level of P130 compared to G0 and numbers indicate the pvalves, Student‵s T-test.

**Figure 4-figure supplement 7.**
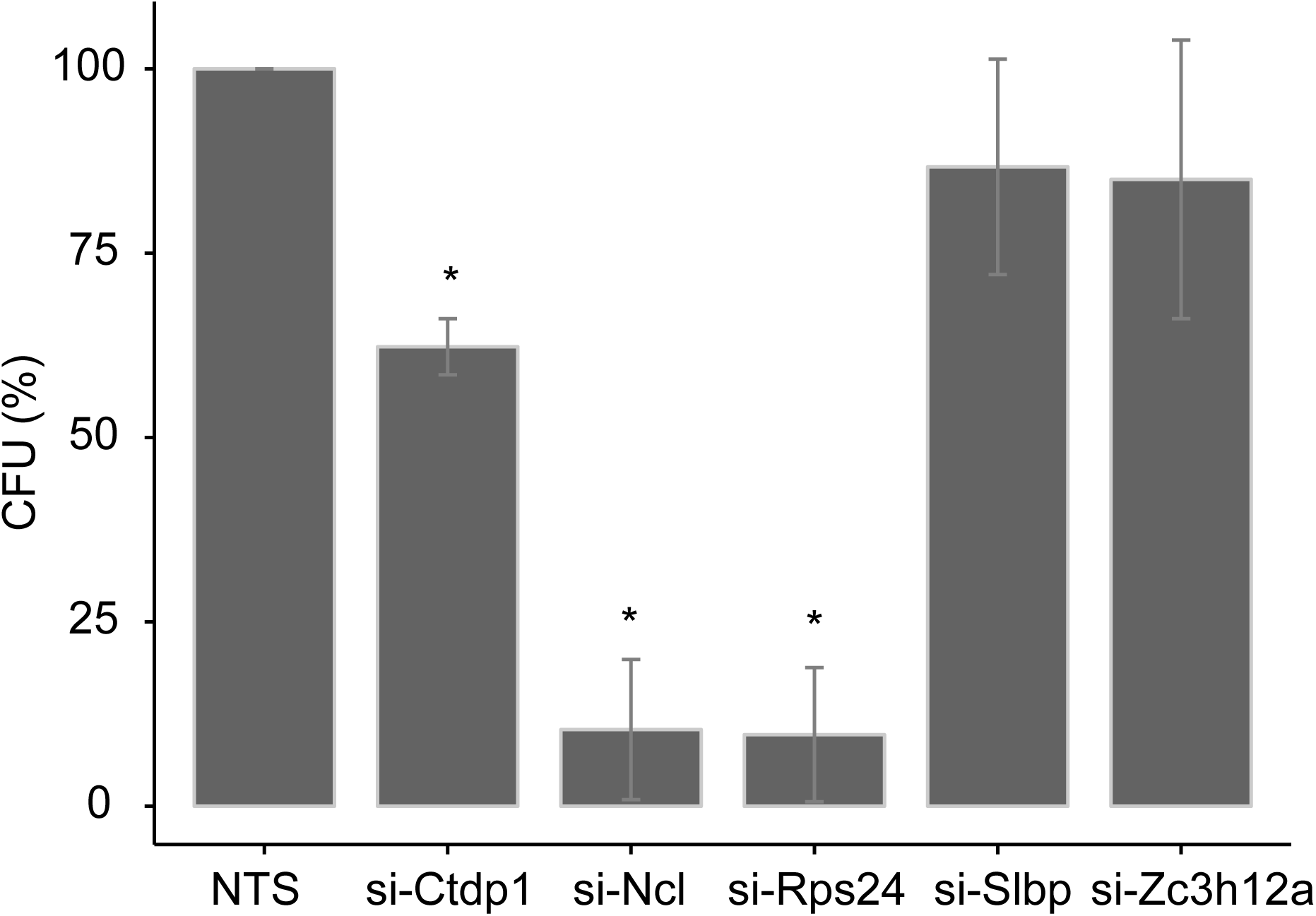
Colony forming assay for G0 cells treated with siRNA for G40 stalled genes represented as % CFU. Reduced self renewal as estimated by colony forming ability for Ncl, Rps24 & Ctdp1 knockdown but not for Slbp and Zc3h12A knock-down (*p-value <0.05, n=3 student‵s T-test)

**Figure 5-figure supplement 1.**
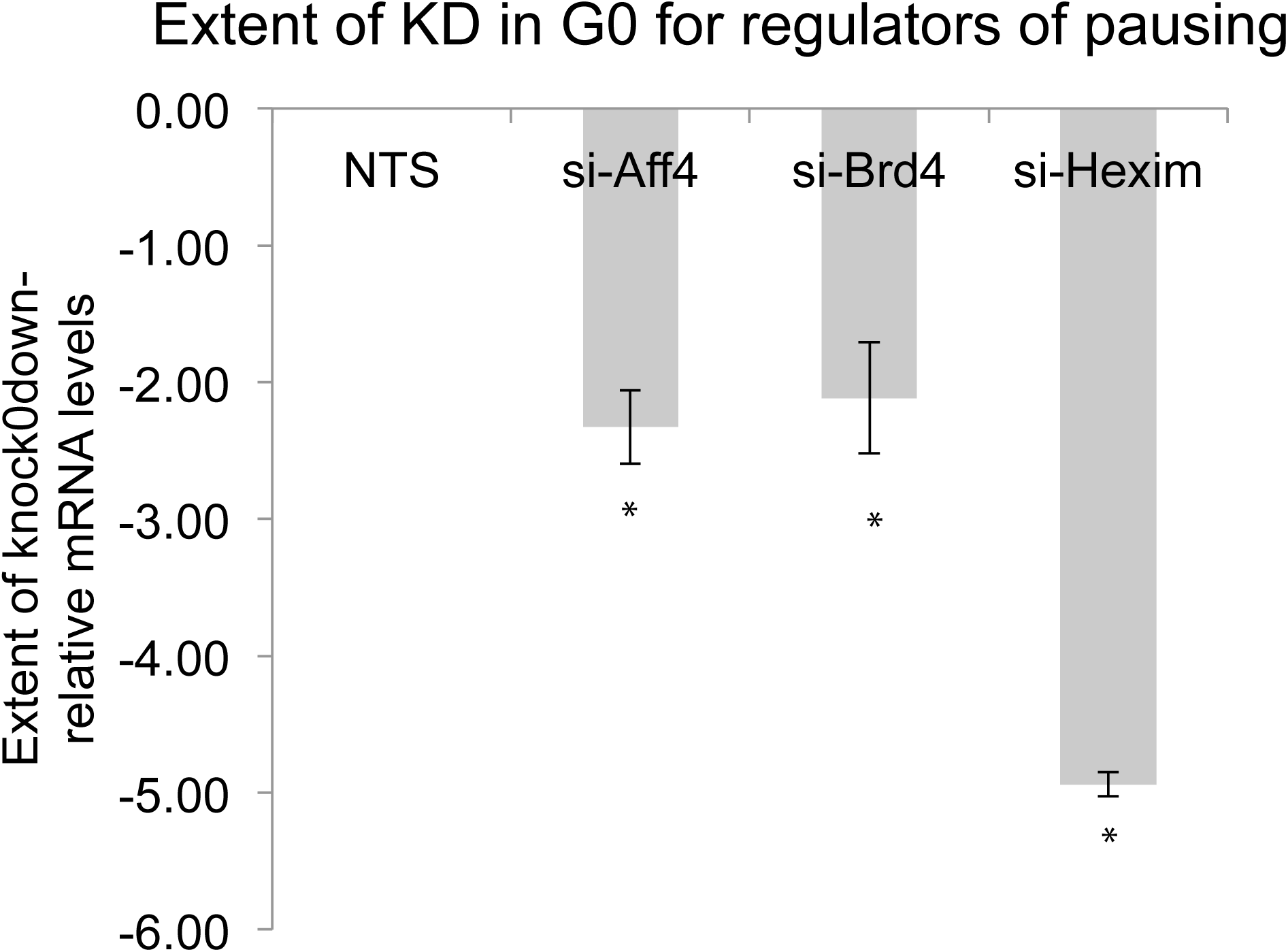
Extent of knockdown in Aff4, Brd4 and Hexim1 in G0 state compared to NTS control. (* ––> pvalue <0.05 Student‵s Ttest, N=3)

**Figure 5-figure supplement 2.**
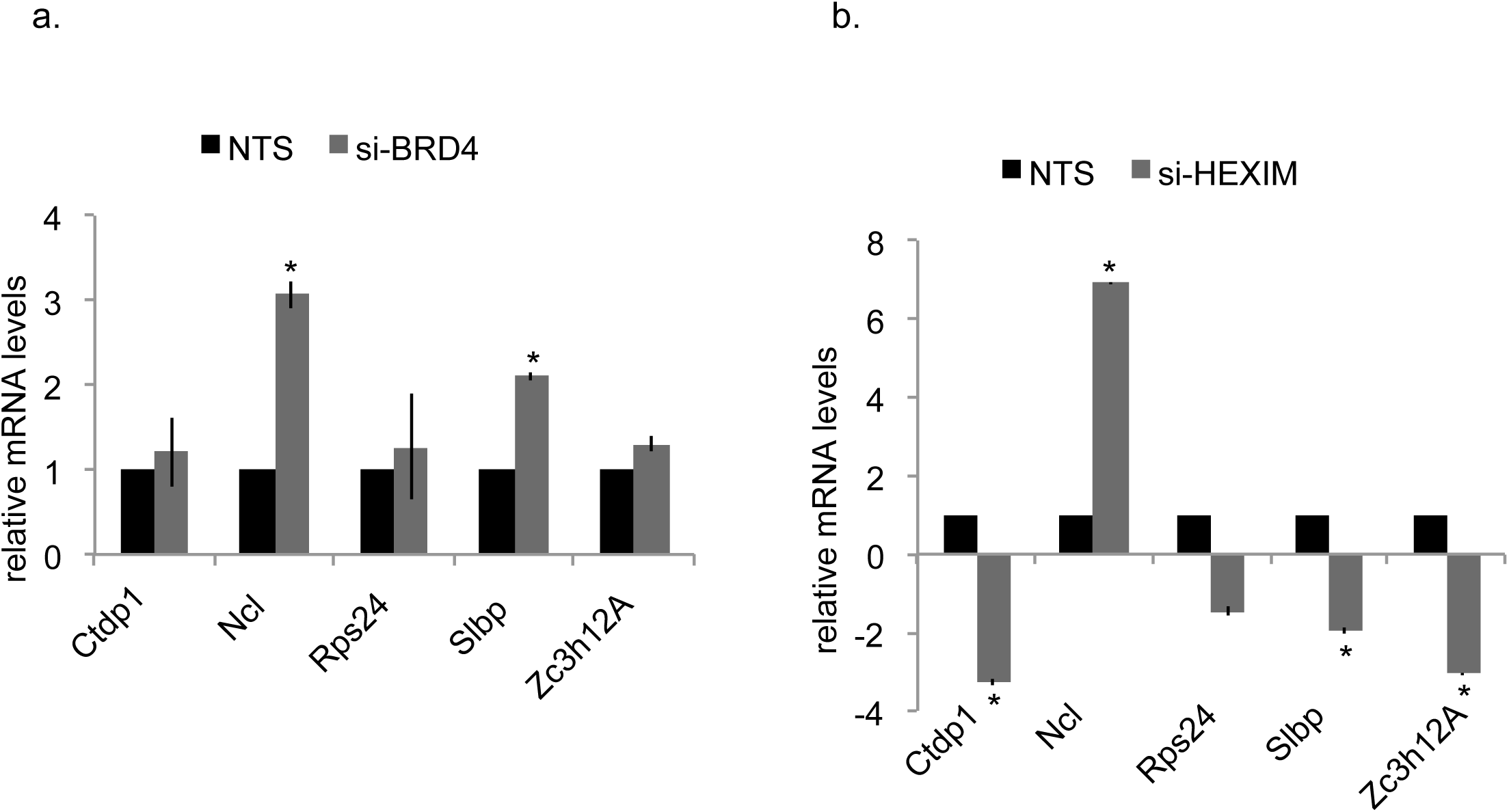
Effect of perturbation of RNA Pol II regulators on expression of G0 stalled genes. Reduced self renewal as estimated by colony forming ability for Ncl, Rps24 and Ctdp1 knockdown. In quiescent cells (G0), knockdown of Brd4 (a) and Hexim1 (b) by siRNA treatment results in altered but not for Slbp and Zc3h12A knockdown (*p-value <0.05, n=3 student‵s T-test) level of expression of G0 stalled genes measured by qRTPCR (n=3 and * =p value <0.05 using students t-test)

**Figure 5-figure supplement 3.**
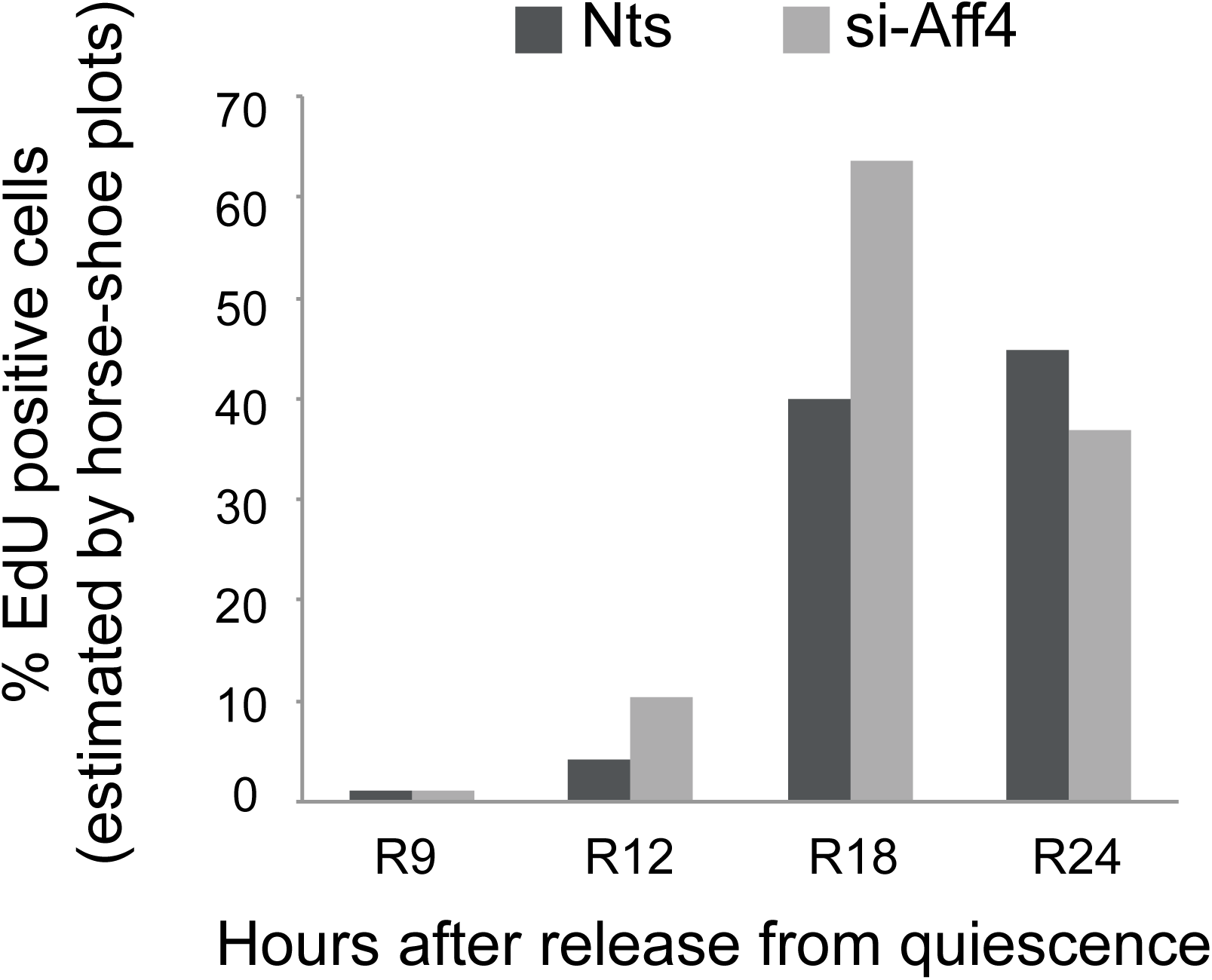
Effect of perturbation of RNA Pol II regulators (Aff4) on cell cycle reentry. The % of EdU positive cells quantified from imaging experiments is observed to be higher in Aff4 knockdown cells at hours R12 and R18 hours after reentry indicative of faster exit from quiescence.

**Figure 5-figure supplement 4.**
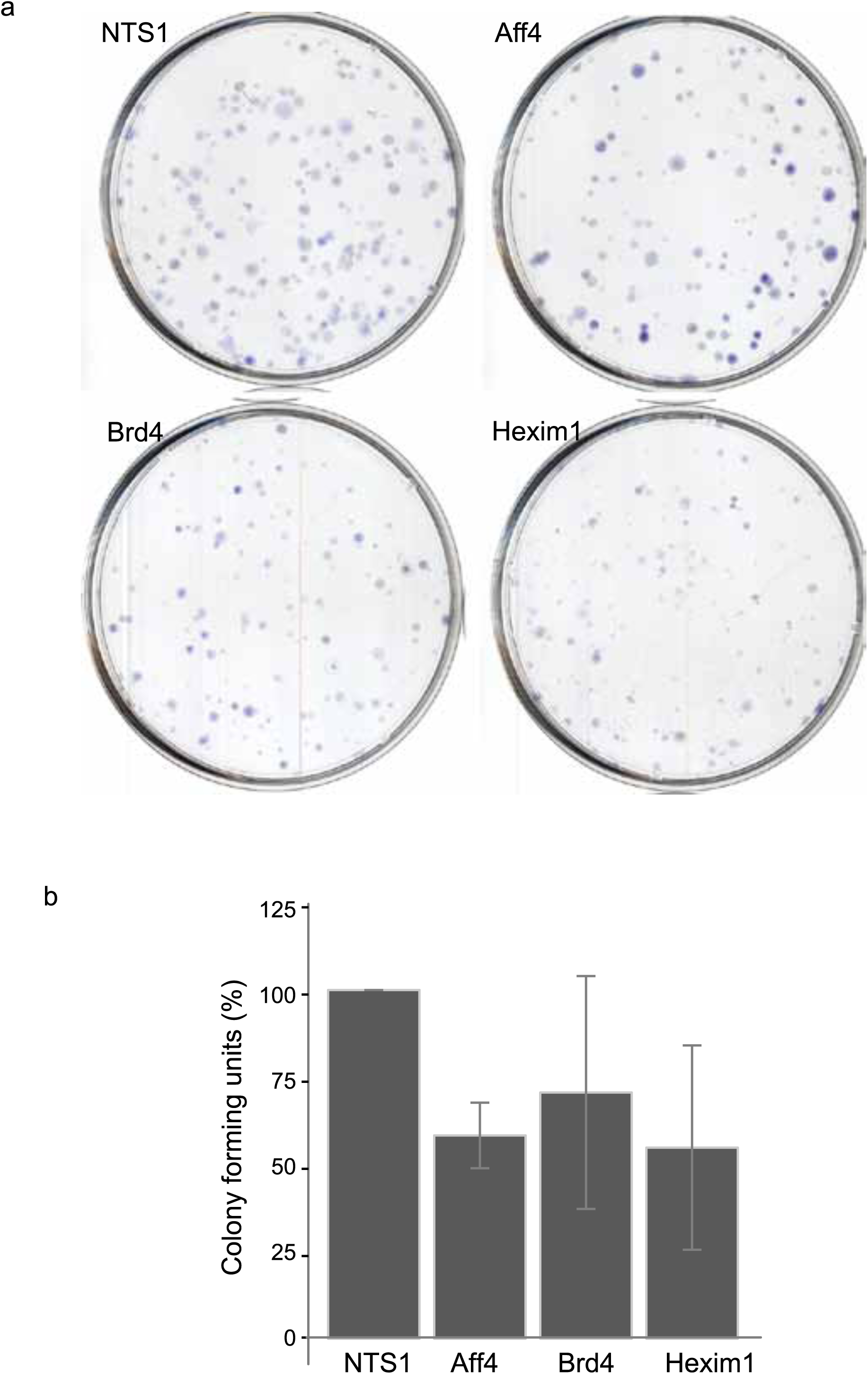
a,b. Knockdown of regulators of RNA Pol II pausing affects self renewal, represented as % CFU. Reduced self renewal as estimated by colony forming ability for Aff4 and Hexim 1 knockdown but not for Brd4 knockdown (*p-value <0.05, n=3 student‵s T-test)

